# Iterative acylation on mature lasso peptides by widespread acetyltransferases for lipolasso production

**DOI:** 10.1101/2024.12.31.630886

**Authors:** Jiang Xiong, Shanquan Wu, Zi-Qi Liang, Shuo Fang, Fen-Yu Tao, Xiao-Tong Gong, Qingfeng Wu, Jiao-Jiao Cui, Kun Gao, Shangwen Luo, Dongsheng Lei, Shi-Hui Dong

## Abstract

The biosynthesis of ribosomally synthesized and post-translationally modified peptides (RiPPs) leverages iterative catalysis to enhance structural and biological diversity. Traditionally, iterative enzymes install multiple post-translational modifications (PTMs) on linear peptides, rather than mature RiPPs with intricate three-dimensional structures, which would require complex changes in substrate binding poses. Here, we present a prolific class of GCN5-related N-acetyltransferases (GNATs) that iteratively and consecutively acylate two Lys residues within the loop and ring motifs of lasso peptides, diverging from the typical iterative modification of linear peptides—an unprecedented function for PTM enzymes. Utilizing high-resolution cryo-electron microscopy and enzymatic reconstitution, we mapped the lasso peptide binding pocket of IatT and pinpointed key residues involved in demarcating the two distinct acetylation steps. Structure-based engineering of IatT’s acetyl group recognition site expanded the cavity to accommodate longer-chain acyl groups, leading to the creation of lipolasso peptides, a novel class of ribosomal lipopeptide. This engineering strategy can be applied to any RiPP BGC encoding GNAT, facilitating the efficient diversification of rare ribosomal lipopeptides.

**Graphic Abstract:** 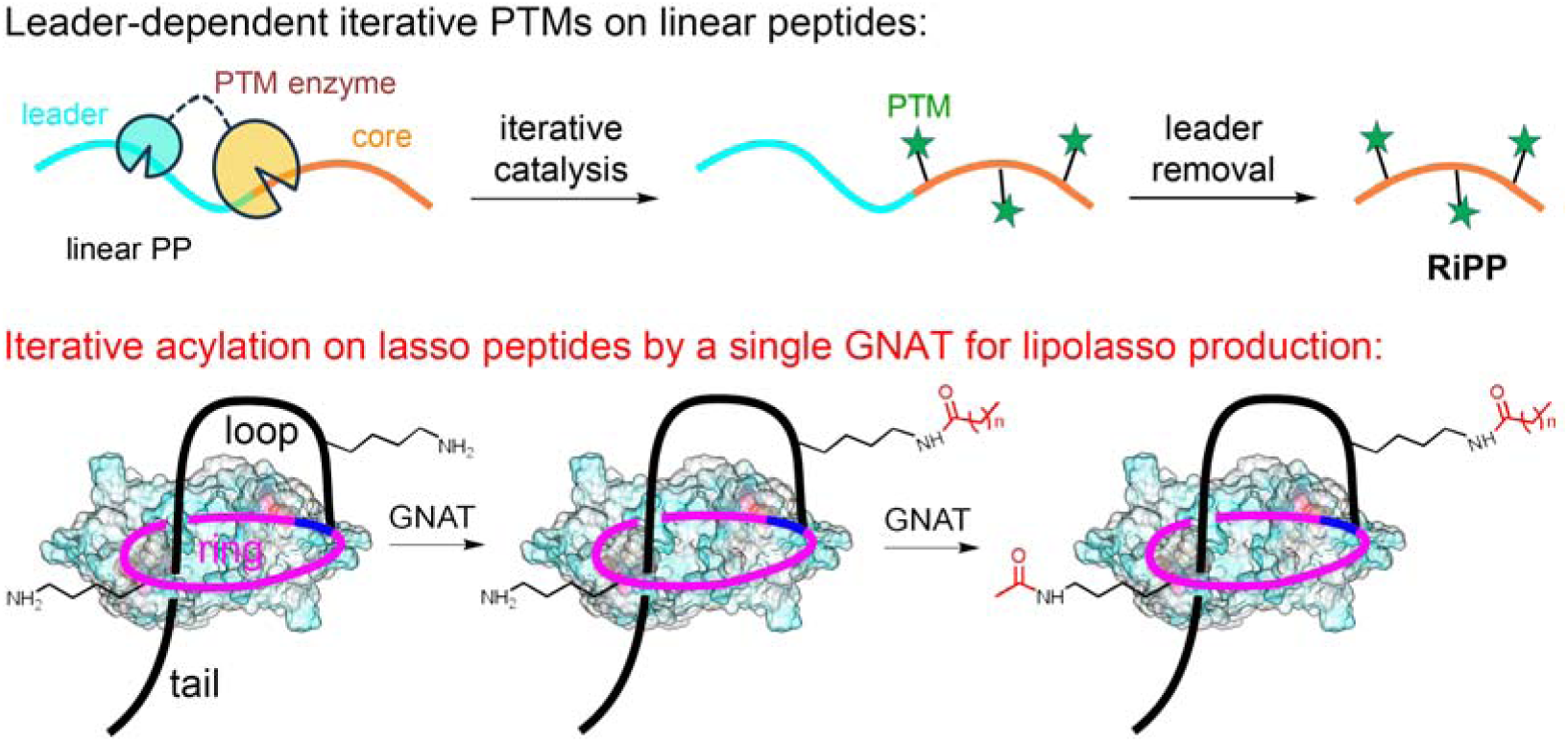

## Introduction

Iterative catalysis, where certain enzymes repeatedly carry out reactions on the same substrates, has been recognized as a hallmark for the biosynthesis of ribosomally synthesized and post-translationally modified peptides (RiPPs) (Fig. 1a), which have been demonstrated as valuable candidates for drug discovery and development^1^. Representative examples of iterative catalysis include lanthipeptide synthases, which construct multiple (methyl)lanthionine residues^2–6^, YcaO enzymes with partner proteins that facilitate the biogenesis of a series of azoles and thioamides^7,8^, and radical SAM enzymes that drive consecutive reactions like C-methylations^9,10^, epimerization^11,12^, and thioether bond formation^13,14^. All reported iterative post-translational modification (PTM) enzymes adhere to the general principle of RiPP biosynthesis: recognizing cognate leader peptides (LPs) and thus iteratively modifying the linear core peptides (CPs) to install the corresponding PTMs (Fig. 1a). However, prior to the current study, no iterative enzyme has been identified that installs multiple PTMs on RiPPs with complex three-dimensional (3D) structures, such as lasso peptides.

**Fig. 1.**
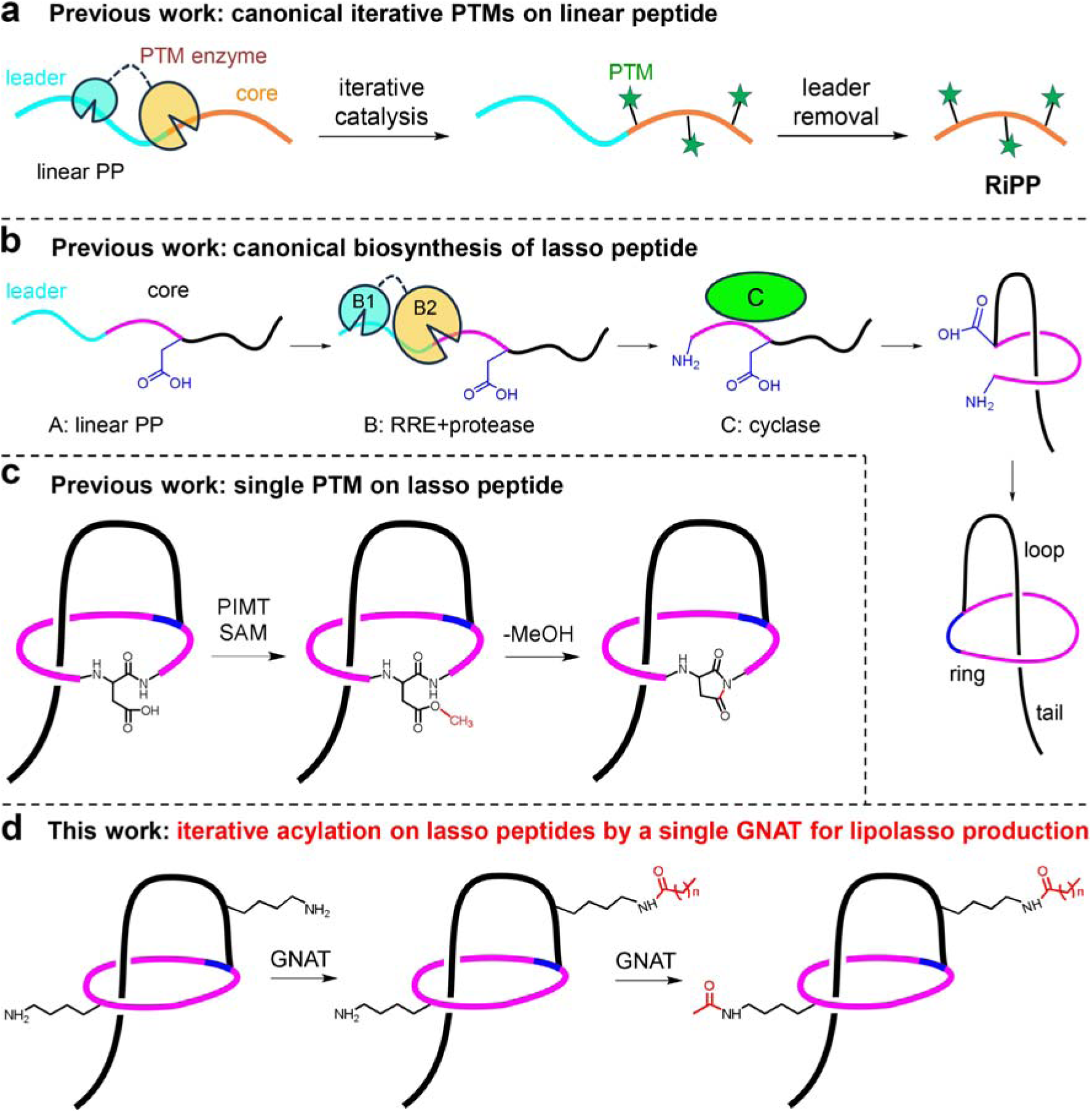
Schematic depiction of representative RiPP PTMs and structures. **a**, Iterative catalysis on linear PPs in RiPP biosynthesis. **b**, General biosynthetic pathway of lasso peptides. **c**, PIMT-catalyzed aspartimide formation with lasso peptide as the substrate. **d**, GNAT-catalyzed iterative acylation on lasso peptides for lipolasso production described in this study.

The unique 3D [1]rotaxane structure, resembling a threaded lariat, is exclusively found in lasso peptides. In the structures of lasso peptides, the C-terminal linear segment is threaded through a macrolactam ring formed between the N-terminus amine and the side chain carboxylate of Asp/Glu residues at position 7-9^2^. As an abundant class of RiPPs, the biosynthesis of lasso peptides begins with the ribosomal synthesis of a precursor peptide (PP, designated as A). This PP is subsequently recognized and processed by a protein complex consisting of a RiPP recognition element (RRE, B1 or E protein), a trans-glutaminase (protease, B2 protein), and a macrolactam synthetase (lasso cyclase) homologous to asparagine synthetase (C protein). In one-third of lasso peptide biosynthetic gene clusters (BGCs), the B1 and B2 proteins fuse into one protein, designated as the B protein. The B1 protein or RRE domain of B protein specifically binds to a well conserved Y(x)_2_PxL(x)_3_G recognition sequence (RS) in the LP of A^15^, which facilitates the B2 protein-catalyzed LP removal, followed by C protein-catalyzed formation of the macrolactam ring in the CP (Fig. 1b). In mature lasso peptides, the C-terminal linear segments are designated as the loop and tail based on their relative positions to the macrolactam ring (Fig. 1b).

Approximately a dozen of secondary PTMs have been characterized during the biosynthesis of lasso peptides, most of which are installed on the linear PPs before the C protein-catalyzed macrolactamization^16^. The only exception is aspartimide formation, catalyzed by a protein L-isoaspartyl methyltransferase (PIMT) homologue^17,18^, which functions solely on the mature lasso structure rather than the linear precursor (Fig. 1c). The molecular mechanism by which this founding member PIMT recognizes the lasso structure remains unclear.

In this work we demonstrate an unprecedented function for a class of GCN5-related N-acetyltransferases (GNATs) that iteratively acylate lasso peptides instead of their linear counterparts (Fig. 1d). We also identified more than 110 lasso peptide BGCs that contain such GNATs. These GNATs can be classified into two classes, GNAT-I and GNAT-II, whose cognate CP sequences encode one and two Lys residues, respectively. Representative GNAT members from both classes were functionally characterized to be able to acetylate both Lys residues on the ring and loop motifs of a lasso peptide. The iterative PTMs at different positions of lasso peptides require significant tuning of the binding poses of lasso peptides in the substrate-binding pocket. We determined the cryo-electron microscopy (cryo-EM) structure of IatT to 2.0 Å resolution and used structure-guided mutational analysis to identify the key residues that function distinctively and cooperatively during the two-step acetylation process. Rational engineering of the acetyl group binding pocket of IatT enabled the production of mono- and diacylated lasso peptides with longer chain-length, resulting in lipolasso peptides, a previously unreported family of ribosomal lipopeptides^19–21^. This engineering strategy for RiPP GNATs will efficiently prepare new-to-nature ribosomal lipopeptides. These abundant GNATs represent the first iterative biocatalysts that function on lasso peptides, thereby expanding the catalytic diversity of such widely distributed GNAT enzyme superfamily.

## Results

### Genome mining of lasso peptide BGCs harboring GNATs

A putative BGC from the strain *Actinosynnema mirum* DSM 43827 first drew our attention (Fig. 2a), which was named as *iat* by the abbreviation of iterative <u>a</u>cetyltransferases. The *iat* BGC displays a similar overall organization to the *alb* BGC that encodes the production of the lasso peptide albusnodin^22^, and consists of genes encoding a PP IatA, an RRE and trans- glutaminase fusion protein IatB, a lasso cyclase IatC, and a putative GNAT IatT (Fig. 2a and Supplementary Table 1). IatA contains a H(x)_2_PxL(x)_4_G sequence in its putative LP, analogous to the known conserved Y(x)_2_PxL(x)_3_G, as the recognition sequence of RRE domain of B protein^23^, along with the highly conserved Thr−2 residue. The putative CP of IatA (IatA_CP_) has Gly1 and Glu8 as potential macrolactamization sites. The most significant difference between IatA_CP_ and AlbA_CP_ is that IatA_CP_ contains two Lys residues at positions 4 and 10, whereas AlbA_CP_ has only one Lys at position 10. The presence of a single Lys residue in AlbA_CP_ corresponds with the characterized structure of albusnodin, which bears a monoacetylation^22^. This variation of CP sequences raises the possibility that IatT may be an unprecedented GNAT capable of iterative acetylation.

**Fig. 2.**
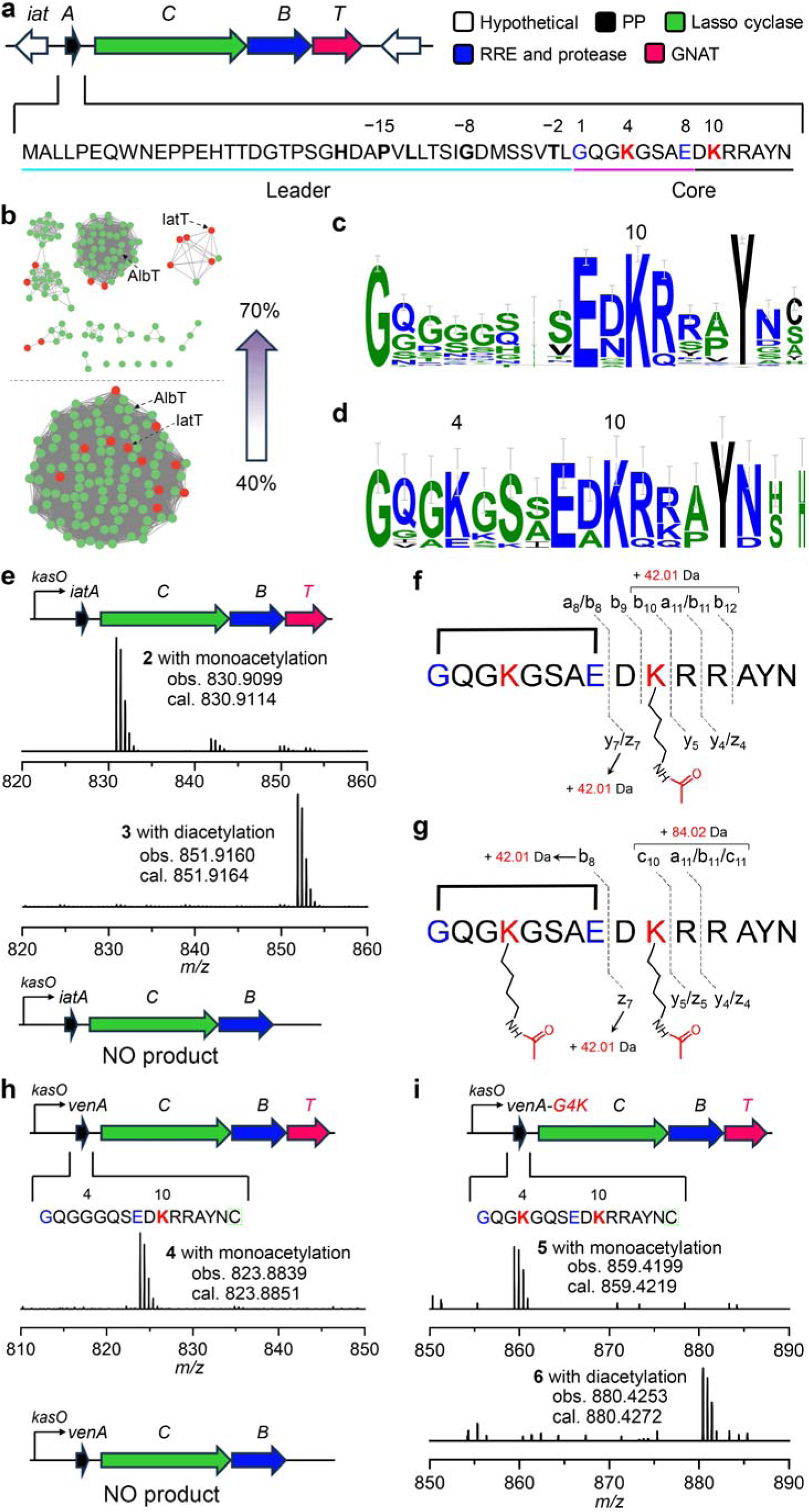
Bioinformatic analysis and heterologous expression of lasso peptide BGCs encoding GNATs. **a**, The *iat* BGC from *A. mirum* DSM 43827. The sequence of PP IatA is shown, with leader and core region and functional residues highlighted. **b**, The IatT-containing SSN cluster with 40% (bottom) and 70% (up) sequence identity threshold. The green and red nodes represent the GNAT sequences of class I and II, respectively. All nodes are nonredundant with identical sequences displayed as a single node. **c**,**d**, Sequence logo plots of the CPs from class I (**c**) and II (**d**) BGCs show the conservation of Lys residues. **e**,**h**,**i**, HRMS data for heterologous expression products of *iat* BGC (**e**), *ven* BGC (**h**), and *ven* BGC with VenA-G4K mutation (**i**). The observed and calculated masses ([M+2H]^2+^) of each product are displayed on the corresponding spectra. The C-terminal Cys residue of VenA highlighted by a green dashed rectangle is not present in **4**-**6**. **f**,**g**, HRMS/MS analysis of **2** (**f**) and **3** (**g**). Detected fragments are indicated. Structures are shown in simplified models with unthreaded conformation for clarity. All assays were run in triplicate and representative results are shown.

To explore the prevalence of IatT-like GNATs, we generated a sequence similarity network (SSN) using the Enzyme Function Initiative Enzyme Similarity Tool (EFI-EST)^24,25^ with IatT sequence as the query. The IatT-containing cluster was readily separated from others using a threshold of 40% sequence identity in the resulting SSN (Supplementary Fig. 1a). This cluster was then analyzed using the EFI Genome Neighborhood Tool (EFI-GNT) to retrieve the co-occurring genes for each GNAT sequence. The resulting Genome Neighborhood Networks (GNNs) and Genome Neighborhood Diagrams (GNDs) revealed that only GNAT sequences in the IatT-containing cluster have neighbors of lasso peptide biosynthetic genes (Supplementary Fig. 1b). Careful analysis of GNDs corresponding to the IatT-containing cluster led to the identification of 112 fully-sequenced GNAT-encoding lasso peptide BGCs (Fig. 2b), all of which are encoded by the Actinomycetes class (Supplementary Fig. 1c).

Sequence comparison of all the lasso peptide PPs enabled us to classify these BGCs into two classes, with *alb* and *iat* as representatives of class I and class II, respectively. Class I encodes CPs with one Lys residue, and class II encodes CPs with two Lys residues. Class I PPs contain 103 members, and sequence alignment demonstrated the presence of a completely conserved Lys10 (Fig. 2c and Supplementary Fig. 1d). In contrast, the 13 sequences from class II contain a highly conserved Lys4 in addition to Lys10 (Fig. 2d and Supplementary Fig. 1e), as observed in IatA. Notably, two exceptions in class II PPs, JiaA from *Jiangella alba* DSM 45237 and AlkA from *Jiangella alkaliphila* DSM 45079, contain the first Lys at position 5 instead of position 4.

To investigate whether the GNAT sequence variations correlate with the observed differences between PP sequences of the two classes, we gradually increased the sequence identity threshold of the GNAT SSN from 40% to 70%, resulting in the dissociation of the original cluster into several smaller ones (Fig. 2b). In the updated SSN with a 70% sequence identity threshold, class II GNATs, shown as red nodes, spread across various clusters along with green class I members, indicating that the similarities in GNAT sequences do not reflect the sequence divergence of their cognate PPs. A Maximum likelihood (ML) tree was also constructed for all GNATs, and sequences from both classes were interspersed, as shown in the SSN (Extended Data Fig. 1), further supporting the above analysis.

### IatT-like enzymes are iterative GNATs in vivo

Bioinformatic analysis suggested that the IatT-like GNATs might be iterative enzymes involved in lasso peptide biosynthesis. To explore this, we investigated the products of the *iat* BGC through heterologous expression of both the full-length and 6.*iatT* BGC under identical conditions^26^. Specifically, the respective DNA fragments were cloned into pSET-kasO vector with a strong *kasO* promoter upstream of the cloning site^26–28^. The two resulting plasmids were individually transformed into *E. coli* ET12567 (pUZ8002) and introduced into *Streptomyces* hosts via *E. coli*/*Streptomyces* conjugation (Supplementary Tables 2 and 3). After screening a set of host strains and culture media, successful expression of the *iat* BGC was achieved in *Streptomyces coelicolor* M1154^29^ using ISP2 media. The metabolites from the M1154-*iat* and M1154-*iat*-6.*iatT* strain cultures were analyzed for signals within the expected mass ranges by matrix-assisted laser desorption/ionization time-of-flight mass spectrometry (MALDI-TOF MS) and liquid-chromatography high-resolution mass spectrometry (LC-HRMS) (Fig. 2e and Supplementary Fig. 2).

The theoretical mass of unacetylated lasso peptide (mirusin A, compound **1**) from the *iat* BGC is 809.9058 ([M + 2H]^2+^), calculated based on the CP sequence of IatA (Fig. 2a). The M1154-*iat* sample displayed two groups of mass peaks with [M + 2H]^2+^ of 830.9099 and 851.9160, which correspond to the mono- and diacetylated lasso peptides (Fig. 2e), named as mirusins B and C (compounds **2** and **3**), respectively. Conversely, M1154-*iat*-6.*iatT* strain did not produce any related products within the same mass range, underscoring the necessity of *iatT* in the biosynthesis of mirusins (Fig. 2e and Supplementary Fig. 2). A similar phenomenon was observed with the heterologous expression of the *alb* BGC, where the removal of *albT* completely abolished the production of albusnodin^22^.

Mirusins were subsequently enriched by chromatographic fractionation and purified by high-performance liquid chromatography (HPLC). The ^1^H NMR signals of both mirusins were highly broad (Supplementary Fig. 3), and their ^1^H-^1^H COSY NMR spectra displayed only a limited number of cross-peaks (Supplementary Fig. 4), precluding NMR-based structural analysis. Therefore, two winding strategies were employed to support the presence of the threaded topology for mirusins. The unlassoed counterpart of a lasso peptide is defined as a branched cyclic peptide, which exhibits distinct physiochemical properties. We first synthesized the branched cyclic peptide corresponding to mirusin B (**2**), named as mirusin B* (**2**\*), for comparison. The structure of **2**\* was confirmed by HRMS/MS analysis (Supplementary Fig. 5). Compound **2**\* displayed sharper and clearer signals in its ^1^H NMR spectrum compared to those of **2** (Supplementary Fig. 3), enabling the acquisition of high quality 2D NMR data and the subsequent signal assignment of **2**\* (Supplementary Fig. 6 and Table 4). Comparison of the ^1^H NMR spectra and elution profiles of **2** and **2**\* suggested that **2** was not the unlassoed branched cyclic peptide (Supplementary Figs. 3 and 7). The broadening of the ^1^H NMR signals of lasso peptides **2** and **3** likely resulted from the conformational exchange of its lariat structure without a dominant conformer.

Additionally, we provide evidences for the existence of the threaded lasso structures using carboxypeptidase Y digestion, a widely used method for this purpose^22,30–32^. Compounds **2**, **2**\*, **3**, and IatA PP were incubated with carboxypeptidase Y for 16 h, followed by MALDI-TOF MS analysis. The PP was completely consumed, and **2**\* was degraded into the cyclic octapeptide and other related shorter branched cyclic peptides, while **2** and **3** remained intact (Supplementary Fig. 7). Moreover, compounds **2** and **3** exhibited similar resistance to carboxypeptidase Y degradation even after 16 h of 95 °C heat treatment (Supplementary Fig. 7). This resistance to carboxypeptidase Y digestion under different conditions supported the presence of the threaded topology for mirusins as lasso peptides. Collectively, these data demonstrate the role of IatT as the first iterative acetyltransferase involved in lasso peptide biosynthesis.

As shown by the sequence alignment of PPs in Figs 2c and 2d, as well as the structure of albusnodin^22^, the iterative acetylation was proposed to occur sequentially on Lys10 and Lys4, which was supported by the HRMS/MS analysis of **2** and **3**. The fragment ions of **2** and **3** were mapped onto their structures (Figs. 2f and 2g and Supplementary Figs. 8 and 9). Differences between the b_9_/b_10_/b_11_ and b_8_ ions of **2** supported monoacetylation on Lys10 (Fig.2f), and the diacetylation of **3** was similarly assigned to Lys10 and Lys4, considering the physiochemical properties of the residues within the macrolactam ring (Fig. 2g).

A class I BGC from *Streptomyces venezuelae* DSM 40230, designated as *ven* (Fig. 2h), was also selected for heterologous expression following a similar protocol. The *ven* BGC is highly similar with *alb* BGC, as supported by the identical sequence of VenA and AlbA^22^. Thus, an identical product to albusnodin was expected from the *ven* BGC. A monoacetylated product with the same mass signals as the albusnodin structure (compound **4**) was successfully detected in the culture of the M1154-*ven* strain but not in that of the M1154-*ven*-6.*venT* strain (Fig. 2h and Supplementary Fig. 10), consistent with observations from the heterologous expression of *iat* and *alb* BGCs. Notably, the C-terminal Cys residue from the CP of VenA was missing in the final product, as was the case for albusnodin derived from AlbA^22^. The main difference between IatA and VenA/AlbA is the presence of Lys4 in IatA, whereas VenA and AlbA have Gly4. This prompted the construction of a mutant *ven* BGC encoding VenA- G4K. Heterologous expression of this mutant *ven* BGC resulted in the production of both mono- and diacetylated products, named as albusnodin B and C (compounds **5** and **6**), respectively (Fig. 2i and Supplementary Fig. 10). These data are consistent with both class I and II GNATs acting as biocatalysts for iterative acetylation involved in lasso peptide biosynthesis, an activity that has not been observed for any other member of this enzyme superfamily.

### IatT-like enzymes are iterative GNATs on lasso peptides in vitro

The heterologous expression experiments could not rule out the possibility that IatT-like GNATs function on the linear PPs as substrates to deliver mono- and diacetylated peptides for the subsequent formation of lasso structure. To test this possibility, we first used AlphaFold-Multimer^33^ to calculate the IatT-IatA complex structure to investigate their potential binding mode. The multimer model was compared with structurally homologous GNATs involved in spermidine/spermine biosynthesis, which are bound with spermine (SpeG from *Bacillus thuringiensis*, PDB code 6VFN^34^) or acetyl-CoA (AcCoA) (SpeG from *Vibrio cholerae*, PDB code 4R57^35^, hereafter designated as vcSpeG). The IatA_CP_ was predicted to occupy the putative substrate binding pocket of IatT, which is analogous to the way SpeG accommodates spermine as the acetyl acceptor, consistent with the role of IatT as a peptidyl acetyltransferase (Extended Data Fig. 2a). The LP of IatA folds as a long loop, which occupies part of the AcCoA space, and extends away from the binding pocket towards the solvent (Extended Data Fig. 2b). The computational binding pattern disagreed with the characteristic binding mode of PTM enzymes to PPs, where the LPs contribute to recognition by the PTM enzymes. Thus, the LP of IatA is unlikely involved in the iterative acetylation.

Next, we overexpressed the PP His_6_-SUMO-IatA alone and together with IatT in *Escherichia coli* to determine whether IatT could acetylate IatA in vivo (Supplementary Fig. 11 and Tables 2 and 3). A tobacco etch virus (TEV) protease recognition site was located between SUMO and IatA, and digestion by TEV protease leads to a tripeptide scar (SNA) fused to the N-terminus of IatA. After Ni-affinity chromatography purification and TEV protease digestion, the LC-HRMS analysis showed that both systems only yielded unmodified IatA (Supplementary Fig. 12). In vitro reactions using purified IatA and His_6_-IatT along with AcCoA and other necessary components resulted in similar mass profiles without any acetylated peptide detected (Fig. 3a). Up to this point, these data have ruled out the linear PP as the substrate for IatT, supporting the hypothesis that IatT may function on lasso peptides at the late stage of the biosynthesis of mirusins.

**Fig. 3.**
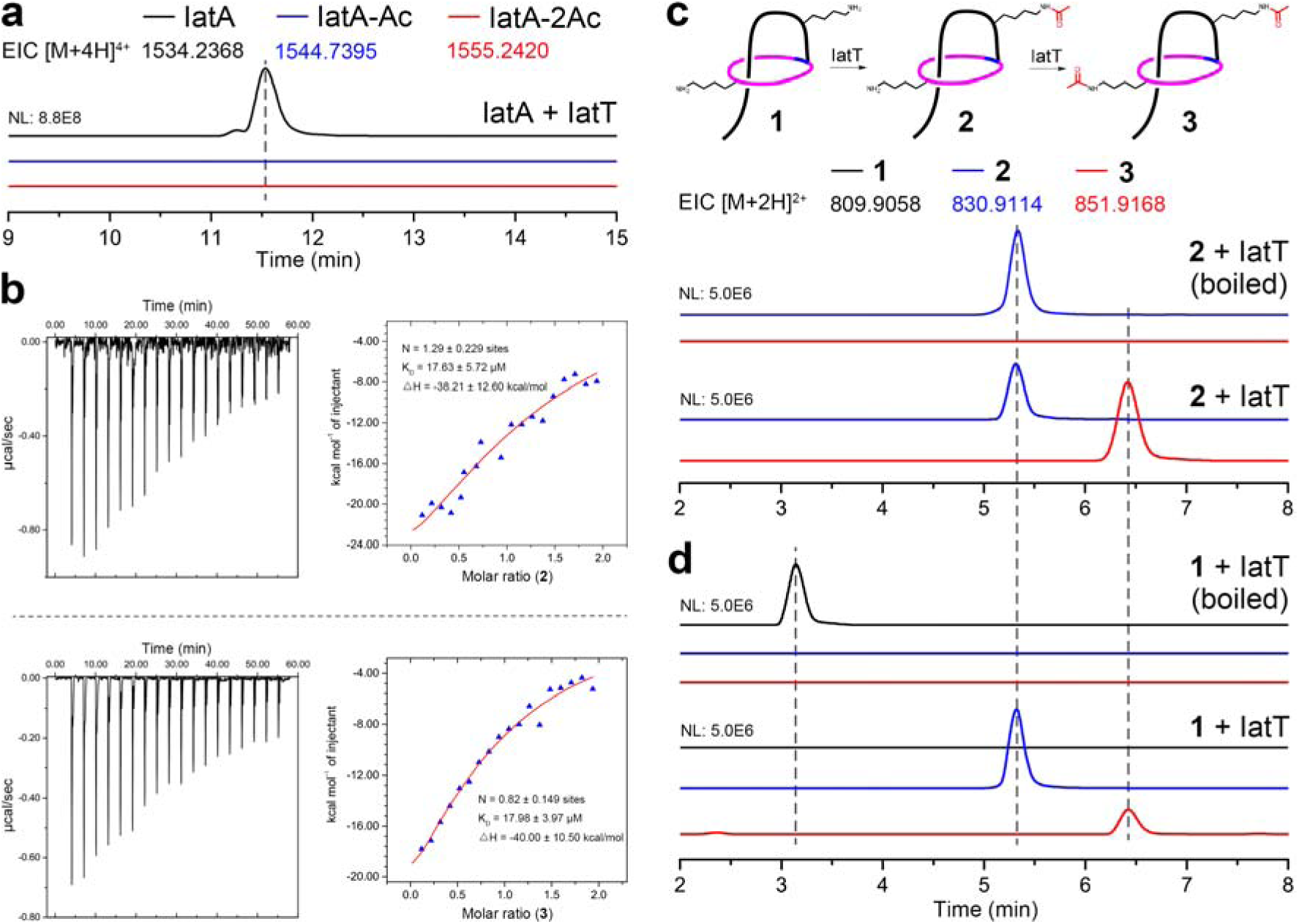
Characterization of IatT in vitro. Enzymatic activity assays of IatT using IatA (**a**), **2** (**c**), and **1** (**d**) as substrates in vitro via LC-HRMS analysis. Shown are extracted ion chromatograms (EICs) for the substrate (black) and monoacetylated (blue) and diacetylated (red) peptides. **b**, ITC data for titration of **2** (upper) and **3** (lower) into IatT solution. Errors represent fitting residuals (standard deviation). All assays were run in triplicate and representative results are shown.

Given that the binding of substrate to IatT is required for catalysis, we subsequently studied the molecular interactions between IatT and its potential substrate and product, including the linear peptide IatA and lasso peptides **2** and **3**, using isothermal titration calorimetry (ITC) (Fig. 3b and Extended Data Fig. 2c). The titration of IatA to IatT revealed no detectable binding, consistent with the predicted model. Conversely, the ITC analysis of lasso peptides indicated that IatT can bind about one molar equivalent of **2** and **3** with similar micromolar affinity, demonstrating a direct interaction between IatT and the lasso peptides.

As the unacetylated mirusin was not available from heterologous expression of *iat* BGC and IatT was proposed to function iteratively, we tested whether IatT could acetylate **2** to form **3** in vitro. Similar in vitro reactions of IatT were carried out using **2** as substrate instead of IatA. The subsequent LC-HRMS analysis revealed a new peak with a yield of about 73.2%, showing the identical mass and retention time with the standard **3** (Fig. 3c). When the branched cyclic peptide **2\*** was subjected to the IatT in vitro reaction, less than 10% of acetylated **2\***, namely **3\***, was produced (Extended Data Fig. 3a). These in vitro reactions clearly demonstrate the function of IatT to acetylate the mature lasso peptide, rather than linear PP or branched cyclic peptide.

To reinforce the iterative catalysis feature of IatT on lasso peptides, the unacetylated lasso peptide mirusin A (**1**) was required. We sought to manipulate the acetyl group binding pocket of IatT to reduce its catalytic efficiency for the production of **1** based on the complex structure of IatT bound with AcCoA, which is discussed in the following sections. A hextuple mutant of IatT was designed, named IatT-HM1, to contain T86E, F99S, I101S, T117R, L135S, and Y148S variations. The corresponding heterologous expression plasmid was constructed to contain the sequence of *iaT*-HM1, and was introduced into the same *Streptomyces* host to yield the M1154-*iat*-HM1 strain. Cultivation of this strain successfully produced **1** in addition to **2** and **3** (Supplementary Fig. 13). The purified **1** was then subjected to in vitro reactions of IatT under same conditions, which completely transformed **1** to **2** and **3** with a ratio of approximately 10:2 (Fig. 3d), consolidating the utilization of lasso peptides as substrates by IatT.

Having characterized the iterative acetylation reactions of IatT on lasso peptides, several additional representative GNATs from both classes of lasso peptide BGCs were selected for in vitro enzymatic transformations (Extended Data Fig. 1 and Supplementary Table 1). The class I GNATs AlbT^22^, VenT, EmbT from *Embleya scabrispora* NF3, and NocT from *Nocardiopsis alborubida* ATCC 23612 were evaluated using **1** and **2** as substrates. The subsequent LC-HRMS analysis demonstrated that these GNATs successfully acetylated both lasso peptides (Extended Data Figs. 3b-3e). The class II GNAT XiaT of from *S. xiamenensis* strain 318 was also tested and found capable of acetylating **1** and **2** (Extended Data Fig. 3f), as expected given the high sequence identity (93%) between the CPs of XiaA and IatA including the Lys4 and Lys10. Additionally, two outliers of class II GNATs, JiaT and AlkT (Extended Data Fig. 1), which are neighbored by cognate PPs containing Lys5 and Lys10 instead of Lys4 and Lys10 as in IatA and XiaA, were also tested. In vitro reactions confirmed that JiaT and AlkT could acetylate **1** and **2** with Lys4 and Lys10 in the lasso structures (Extended Data Figs. 3g and 3h). Collectively, the successful biochemical reconstitution of multiple GNATs from both classes using mirusin A (**1**) and B (**2**) as substrates clearly demonstrates that these GNATs, with over 110 members, are unprecedented lasso peptide iterative acetyltransferases, which serve as the final steps for the maturation of acetylated lasso peptides (Fig. 1d).

### Structural basis for GNAT-mediated iterative acetylation on lasso peptides

To understand how a single GNAT recognizes different lasso peptides for iterative catalysis, we determined the structures of IatT in the presence of CoA and lasso peptide **1**, as well as in the presence of AcCoA and lasso peptide **2** using cryo-EM. After data processing, two final density maps with average resolutions of 2.2 Å and 2.0 Å were obtained, respectively (Fig. 4a, Extended Data Figs. 4 and 5, and Supplementary Table 5). The density maps are of sufficient quality to allow unbiased de novo model building for IatT and CoA or AcCoA (Fig. 4b). The lasso peptides could not be confidently modeled in the final structures. Both structures of IatT-CoA and IatT-AcCoA are highly similar. Thus, the IatT-AcCoA complex, with higher resolution and more homogeneous density map, was primarily discussed in the following sections. The final atomic models of IatT-AcCoA and IatT-CoA appear to be tetradecameric, with overall dimensions of approximately 115 Å × 115 Å × 85 Å (Fig. 4b), which includes 2,646 amino acids with 189 residues from each chain (residues 5–193). The IatT tetradecamer can be described as a dimer of heptamers, where the two heptamers stack on top of each other to form a cylinder-like architecture.

**Fig. 4.**
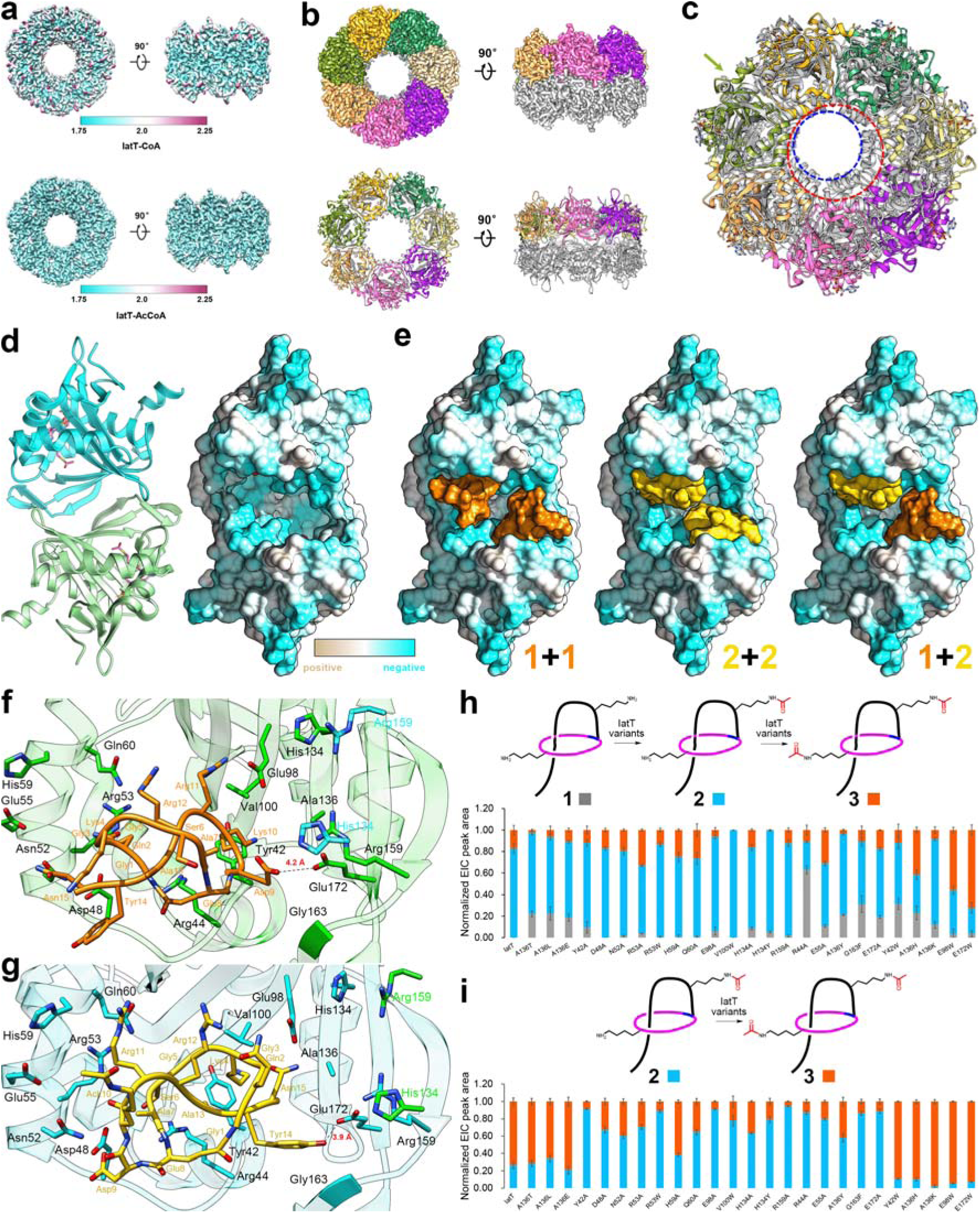
Cryo-EM structures and mutagenesis studies of IatT. **a**, Density maps depicting the local resolution across the tetradecamer structures of the IatT-CoA (up) and IatT-AcCoA (bottom). **b**, Cartoon representation of the overall cylinder structure of IatT-AcCoA tetradecamer in surface (up) and ribbon (bottom) models. **c**, Comparison of the structures of the IatT tetradecamer with colored monomer and vcSpeG (grey PDB code 4R57^35^) dodecamer. A IatT monomer (arrow) is superimposed with a monomer of vcSpeG to show the different sizes of the interior dimensions of the IatT tetradecamer (red dashed circle) and vcSpeG dodecamer (blue dashed circle). **d**, Structures of IatT stacked dimers showing the lasso peptide binding pocket. Left, a ribbon model with one monomer in cyan and the other in green, and AcCoA in pink sticks. Right, a surface model showing electrostatic potential map. **e**, Representative molecular docking models of two molecules of **1** (orange) and/or **2** (yellow) in the IatT substrate binding pocket. The surface models of **1** are derived from conformation **1a**. The two identical lasso models in the same binding pocket (**1**+**1** or **2**+**2**) are obtained through simple flip to display C2 symmetry. **f**,**g**, Key residues of the lasso peptide binding pocket in the IatT- AcCoA structure modelled with **1** (conformation **1a**) (**f**) and **2** (**g**). Two IatT monomers are shown in green and cyan, respectively. The His134 and Arg159 from one monomer point toward the other monomer. **h**,**i**, Comparison of in vitro reaction results of IatT and its variants using **1** (**h**) and **2** (**i**) as substrates. Bar graphs represent normalized relative EIC area means (nL=L3 biologically independent experiments) of **1**-**3**, and the error bars areLstandard deviation of defined species.

A PDB search using DALI server^36^ revealed that IatT is structurally homologous to many spermidine N-acetyltransferases, despite low sequence identity (Supplementary Fig. 14). These spermidine GNATs typically form dodecamers with smaller overall sizes compared to the tetradecameric IatT. The closest structural homolog, vcSpeG, has overall dimensions of approximately 103 Å × 103 Å × 72 Å^35^. Structural superimposition of the IatT tetradecamer and vcSpeG dodecamer clearly shows that the inner dimension of the IatT cylinder is larger than that of vcSpeG by about 12 Å (Fig. 4c), reflecting the larger size of lasso peptides compared to spermidine/spermine as substrates.

The AcCoA binding poses of IatT and vcSpeG are mostly conserved with slight shift, but there are variations in the position of the adenine moiety (Supplementary Fig. 15). Additionally, the acetyl groups display a rotation of about 90 degrees. Following the direction of the AcCoA chain, a large pocket for the acetyl group acceptor is observed between the two adjacent, inverted monomers with C2 symmetry, each belonging to one of the stacked heptamers (Fig. 4d). The electrostatic potential analysis shows that the binding pocket surface is relatively polar, consistent with the structures of **1** and **2** containing more polar residues.

The abovementioned ITC experiments suggested a 1:1 ratio of substrate lasso peptides with IatT, which indicated that two lasso peptide molecules are expected to be accommodated in the binding pocket formed by two IatT monomers. To validate the structural information, we performed molecular docking for compounds **1** and **2** with the IatT-AcCoA complex structure. The lasso conformations of **1** and **2** were initially predicted using the LassoHTP method^37,38^, which were further optimized through enhanced sampling to yield preferential conformations. Considering the nucleophilic mechanism of the acetylation reaction, we applied three screening criteria to the molecular docking results: 1) the distances from ε-amine of Lys10 in **1** or Lys4 in **2** to the carbonyl carbon of the acetyl group in AcCoA were no more than 5.5 Å; 2) the dihedral angels between the C-N and C-O bonds in the emerging intermediates were less than 36°; 3) The two docking lasso structures in the same binding pocket displayed no overlap. Applying these screening criteria resulted in 49 docking models of two molecules of **1** and/or **2** within the predicted binding pocket of IatT-AcCoA dimeric complex. The results support the authenticity and the adequate space of the binding pocket in IatT for accommodating different combinations of two lasso peptide molecules (Fig. 4e).

### Site-directed mutational and mechanistic analysis of IatT

In these 49 IatT-lasso peptide models, which feature various combinations of lasso conformations, lasso peptide **1** displayed eight different conformations, designated as conformations **1a** to **1h** (Fig. 4f and Extended Data Figs. 6 and 7). In contrast, only one conformation was obtained for lasso peptide **2** (Fig. 4g). Conformation **1a** was the most prevalent, appearing in 13 models (Fig. 4f). The second, third, and fifth most prevalent conformations were **1b**, **1c**, and **1e**, which appeared in 12, 9, and 3 models, respectively, and displayed a similar overall layout compared to conformation **1a**. Therefore, conformations **1a**, **1b**, **1c**, and **1e** are likely to be the dominant conformations for initial acetylation.

Although lasso peptides **1** and **2** could be modeled into the same pocket in the IatT-AcCoA structure, they displayed completely inverted binding poses in order to deliver Lys10 on the loop of **1** and Lys4 on the ring of **2** to the reaction site (Figs. 4f and 4g). A set of residues with various physicochemical properties surrounding **1** and **2** was identified, including Tyr42, Arg44, Asp48, Asn52, Arg53, Glu55, His59, Gln60, Glu98, Val100, His134, Ala136, Arg159, Gly163, and Glu172. These 15 residues showed various degrees of conservation across 112 IatT-like GNATs (Supplementary Fig. 16). To identify the key determinants of IatT for the iterative acetylation of lasso peptides, particularly those controlling the binding pose change after the first acetylation, each of these residues was mutated to Ala or to residues with opposite physicochemical properties. The catalytic efficiencies of these IatT variants were evaluated in vitro using **1** and **2** as substrates (Figs. 4h and 4i). The activity of the variants was quantitively assessed using an endpoint LC-HRMS assay.

The 25 IatT variants of the 15 residues had varying effects on the transformation of **1** to **2** (first acetylation) and/or **2** to **3** (second acetylation) (Figs. 4h and 4i and Supplementary Table 6), allowing the classification of all variants into five categories based on their transformation ratios. 1) Unaffected second acetylation but reduced first acetylation efficiency: variants IatT- A136T, A136L, and A136E showed a significant decrease in the first acetylation step, leaving 19.9% to 21.9% of **1** untransformed compared to the complete turnover of **1** by wild type IatT, whereas the second acetylation remained similarly effective (Supplementary Fig. 17). 2) Unaffected first acetylation but reduced second acetylation efficiency: variants including IatT-Y42A, D48A, N52A, R53A, R53W, H59A, Q60A, E98A, V100W, H134A, H134Y, and R159A did not significantly affect the first acetylation but showed reduced conversion ratios for the second acetylation (Supplementary Fig. 18). 3) Reduced activity for both acetylation steps: variants IatT-R44A, E55A, A136Y, G163F, and E172A exhibited significantly reduced activity for both the first and second acetylation reactions (Supplementary Fig. 19). 4) Opposite effects on two acetylation steps: variants such as IatT-Y42W, A136H, and A136K displayed decreased activity for the first acetylation but increased activity for the second (Supplementary Fig. 20). 5) Improved activity for both steps of iterative acetylation: two stand out variants, IatT-E98W and E172W, showed highly enhanced efficiency for both acetylation steps compared to wild type IatT (Supplementary Fig. 21). IatT-E98W produced 52.3% and 94.6% of **3** from **1** and **2**, respectively. The E172W variant was even more efficient, yielding 83.3% and 92.5% of **3** from **1** and **2**, respectively. The larger aromatic side chain of Trp compared to Glu may create a more contracted yet fitting binding pocket for the substrate lasso peptides. Given that Glu98 and Glu172 are the most conserved residues in all IatT-like GNATs (Supplementary Fig. 17), their corresponding Trp mutations could serve as a general strategy for generating improved iterative biocatalysts for lasso peptides. These mutational studies further demonstrated the significance of these residues in the putative lasso peptide binding pocket and suggested that they play different roles in the two-step acetylation process.

The side chain carboxyl group of IatT Glu172 points toward the side chain carboxyl group of Asp9 in **1** (4.2 Å; Fig. 4f) and the side chain hydroxy group of Tyr14 in **2** (3.9 Å; Fig. 4g). The hydrogen bonding could facilitate the proper locking of each substrate lasso peptide, aligning with the reduced activity observed for the IatT-E172A variant in both acetylation steps (Figs. 4h and 4i). When Glu172 of IatT is mutated to Trp, the aromatic side chain of Trp may have a strong π-π interaction with Tyr14 of **2**, one of the completely conserved residues in all identified CPs (Figs. 2c and 2d), which explains the significantly improved efficiency of the IatT-E172W variant. Lys10 of **1** and Lys4 of **2**, as the acetyl group acceptors, are surrounded by the Tyr42, Glu98, Val100, and Ala136 residues of IatT, and the mutations of these IatT residues indeed affected both steps of acetylation (Figs. 4h and 4i).

The structural models of IatT revealed that residues Arg44, Asp48, Asn52, Arg53, Glu55, His59, and Gln60 formed a cavity, which could accommodate Gln2 of **1** or Arg11 of **2** (Figs. 4f and 4g and Extended Data Figs. 6a-6c and 7d). The IatT-D48A, N52A, R53A, H59A, and Q60A variants only reduced the efficiency of the second acetylation, whereas R44A and E55A exhibited reduced activity for both acetylation steps (Figs. 4h and 4i). Sequence alignment of all IatT-like GNATs revealed the conservation of Arg44 (with major substitutions of Asn or Thr), Asp48 (with major substitutions of Glu or Gln), Asn52 (with major substitutions of Ala, Val, or Ser), Arg53, Glu55 (with major substitutions of Ala or Gly), His59 (with major substitutions of Arg or Ala), and Gln60 (Supplementary Fig. 17). Additionally, Arg11 of IatA is one of the most conserved residues in all bioinformatically identified CP sequences, with Gln being the only alternative that might serve as a similar function (Figs. 2c and 2d). Gln2 of IatA is also the most conserved at this position, with Gly, Ser, and Asn as frequent replacements (Figs. 2c and 2d). Collectively, these data showed that this cavity displayed different interactions with Gln2 of **1** and Arg11 of **2**, which may facilitate the transformation of the binding poses of **1** and **2** required for the iterative catalysis of lasso peptides.

### Engineering of IatT for the production of lipolasso peptides

Given that known lipopeptides predominantly originate from nonribosomal pathways^39,40^, the acylation of RiPPs among the diverse and rapidly growing PTMs has garnered significant interests, which produces ribosomal lipopeptides to expand the structural and biological diversity. To broaden the border of ribosomal lipopeptides, we aimed to build lipolasso peptides by engineering the acetyl group binding pocket of IatT to accommodate larger acyl groups. In the IatT-AcCoA complex structure, the acetyl group points toward a small hydrophobic pit, which is surrounded by residues Phe99, Ile101, Thr117, Leu135, and Tyr148 (Fig. 5a). Additionally, Thr86 reinforces the back of this hydrophobic pit. Consequently, the IatT-HM2 variant, carrying T86A, F99A, I101A, T117A, L135A, and Y148A mutations, is expected to possess a larger cavity suitable for longer chain acyl groups (Figs. 5b and 5c).

**Fig. 5.**
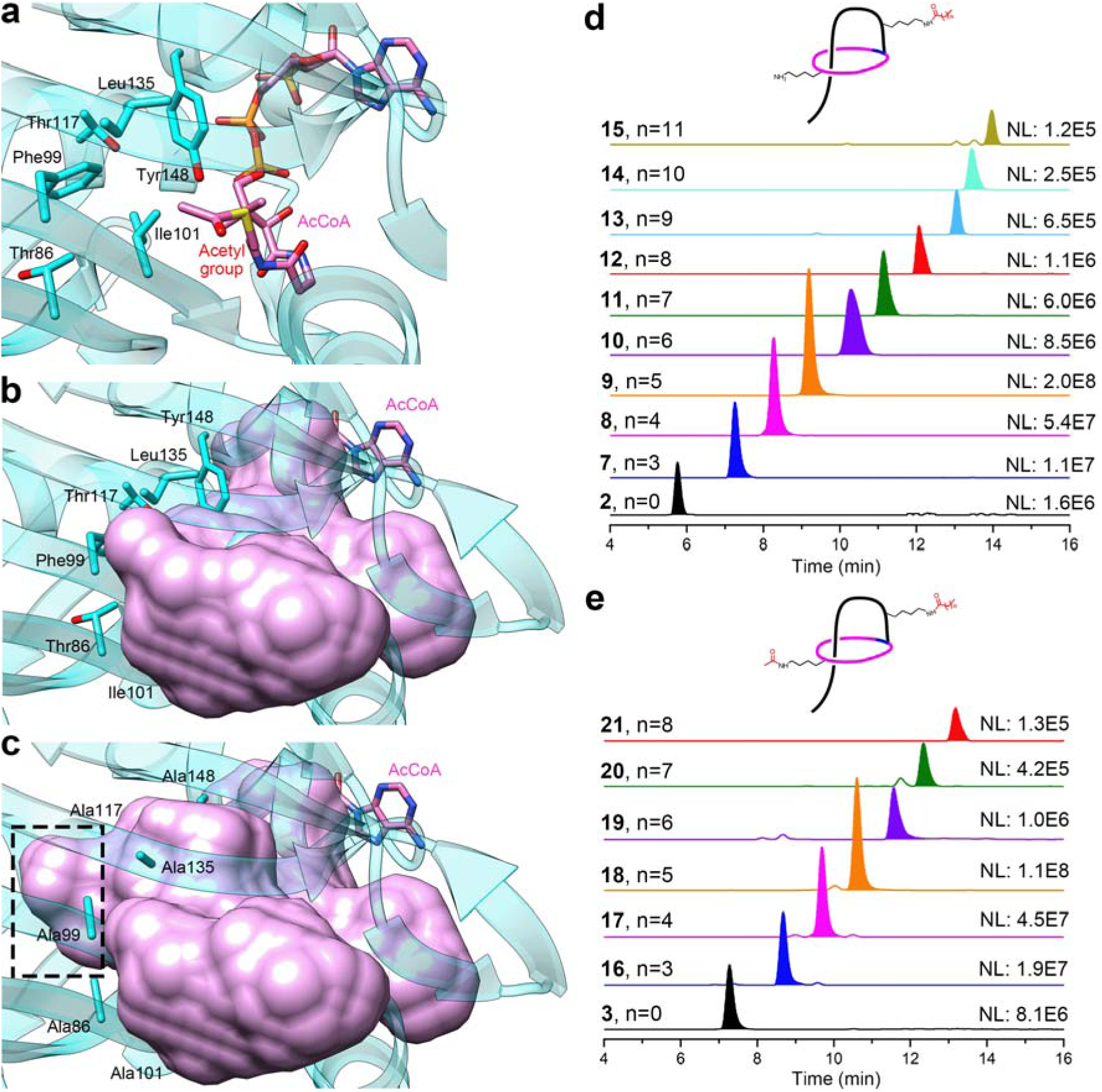
Engineering studies of IatT. **a**, Key residues of the acetyl group binding pocket in the IatT- AcCoA structure. **b**,**c**, Comparison of the AcCoA binding pocket sizes in IatT (**b**) and IatT-HM2 (**c**). The AcCoA binding pockets are shown in pink surfaces. The enlarged portion of the acyl group binding pocket in IatT-HM2 is indicated by a black dashed rectangle. The Ile101 in IatT, as well as Ala99 and Ala117 in IatT-HM2, are blocked by the binding pocket surfaces. **d**,**e**, LC-HRMS analysis of monoacylated (**d**) and diacylated (**e**) lipolasso peptides from the M1154-*iat*-HM2 metabolites. Shown are EICs of target compounds with less than 5 ppm of mass errors. All assays were run in triplicate and representative results are shown.

To test the above hypothesis, the *iatT*-HM2 gene was integrated into the same heterologous system to produce the M1154-*iat*-HM2 strain. Metabolite analysis of this strain using LC-HRMS revealed the production of 15 lipolasso peptides, in addition to compounds **2** and **3**. Nine monoacylated lipolasso peptides (compounds **7**-**15**) were identified, with acyl chain lengths ranging from C5 to C13 (Fig. 5d). Tandem MS analysis of representative members confirmed the monoacylation at Lys10 (Supplementary Figs. 22-24), consistent with the characterized order of acylation. Additionally, six diacylated lipolasso peptides were produced (Fig. 5e), demonstrating the conservation of iterative catalysis capability of IatT- HM2. Interestingly, tandem MS analysis showed that the acyl groups on Lys10 ranged from C5 to C10, while only the acetyl group was attached to Lys4 (Supplementary Figs. 25-27). Possible explanations for the acyl group selectivity observed during the first and second rounds of acylation may include: 1) the natural abundance of different acyl CoAs restricts the diversity of the second acylation, and/or 2) after the initial acylation of Lys10, longer chain acyl groups induce significant conformation changes that hinder the binding of larger acyl CoAs for subsequent acylation.

## Discussion

Although iterative catalysis has been widely documented for significantly expanding the structural diversity and complexity of RiPPs, iterative PTM enzymes classically modify linear PPs by anchoring LPs and sliding CPs through their active sites^2^. Here, however, we demonstrate a novel and unprecedented iterative catalytic mode for a class of GNATs with over 110 members on lasso peptides, where LPs have been cleaved and CPs have been reshaped into a lariat conformation (Fig. 1). Interestingly, the unique iterative catalytic feature on lasso peptides is not exclusive to class II GNATs co-occurring with PPs containing two Lys residues in their cores; class I GNAT members also exhibit iterative functionality. These biochemical findings are consistent with the phylogenetic analysis of lasso peptide GNATs.

The presence of a single Lys residue in the CPs, rather than the GNATs themselves, is the key factor in the lack of discovery of iteratively acetylated lasso peptide.

High-resolution cryo-EM structures of the IatT-AcCoA complex revealed a distinct overall architecture compared to orthologous GNATs that use small molecules as substrates. The most obvious difference lies in their self-assembly states: IatT, and potentially all lasso peptide GNATs, oligomerize into a tetradecamer, whereas small molecule GNATs form at most a dodecamer. This evolution of oligomerization states results in the formation of cylinder-like structures with varied dimensions, reflecting the disparate sizes of their cognate substrates, such as lasso peptides with at least 15 residues or spermines.

The two stacked monomers of IatT with C2 symmetry constitute a large cavity, the entrance of which faces the interior space of the IatT cylinder. Additionally, the acetyl groups of two AcCoA molecules bound with the two IatT monomers point towards the cavity, supporting the cavity as the pocket to accommodate substrate lasso peptides. Computational studies successfully modeled two lasso peptide molecules with no and/or monoacetylation in various conformations into this pocket, supporting the proposed role of the cavity for lasso peptide binding and rotation required by iterative catalysis.

Structure-based mutational studies identified several key residues surrounding the lasso peptide binding pocket, each functioning differently in the two-step consecutive acetylation processes. All mutants of IatT Ala136 resulted in compromised activity for the first acetylation, while their effects on the second acetylation varied according to the physicochemical features of the mutant residues. The residues Tyr42, Asp48, Asn52, Arg53, His59, Gln60, Glu98, Val100, His34, and Arg159 were primarily involved in the second acetylation, while Arg44, Glu55, Gly163, and Glu172 likely participated in both acetylation steps. Two superior variants, IatT-E98W and E172W, were characterized as more efficient in transforming non-acetylated lasso peptides to the diacetylated form. Substitution by Trp is expected to occupy more space and thus squeeze the binding pocket size, potentially allowing for more appropriate binding of the monoacetylated lasso peptide after conformational movement. Additionally, the potential π-π interaction between IatT Trp172 and Tyr14 of **2** may increase their affinity and thus enhance the second acetylation.

Currently, known lipopeptides are predominantly biosynthesized by nonribosomal pathways^39,40^, as exemplified by clinical antibiotics daptomycin, polymyxin, and echinocandin etc. So far, only three classes of ribosomal lipopeptides have been discovered recently, namely lipolanthines^19,20^, selidamides^21^, and lipoavitides (Figs. 6a-6c), which feature hybridization with polyketide/lanthipeptide, fatty acid/proteusin, and fatty acid/aminovinylmethyl-cysteine (AviMeCys)-containing peptide, respectively. Enzymes responsible for the condensation of these RiPPs and fatty acyl units all belong to the GNAT superfamily, highlighting the potential of GNATs in the divergent biosynthesis of ribosomal lipopeptides. Notably, the functions of all characterized GNATs are limited to a single transfer of the acyl group. The scarcity of ribosomal lipopeptides motivated us to engineer the acetyl group binding pocket for a deeper tunnel, leading to the production of lipolasso peptides as a new class of ribosomal lipopeptides (Fig. 6d). Moreover, several GNATs have been shown to catalyze N-acetylation of other classes of RiPPs, including linear azol(in)e-containing peptides^41^ and graspetides^42^. Our strategy of GNAT engineering illustrates an efficient approach to creating new-to-nature classes of ribosomal lipopeptides, thus expanding the diversity of lipopeptides.

**Fig. 6.**
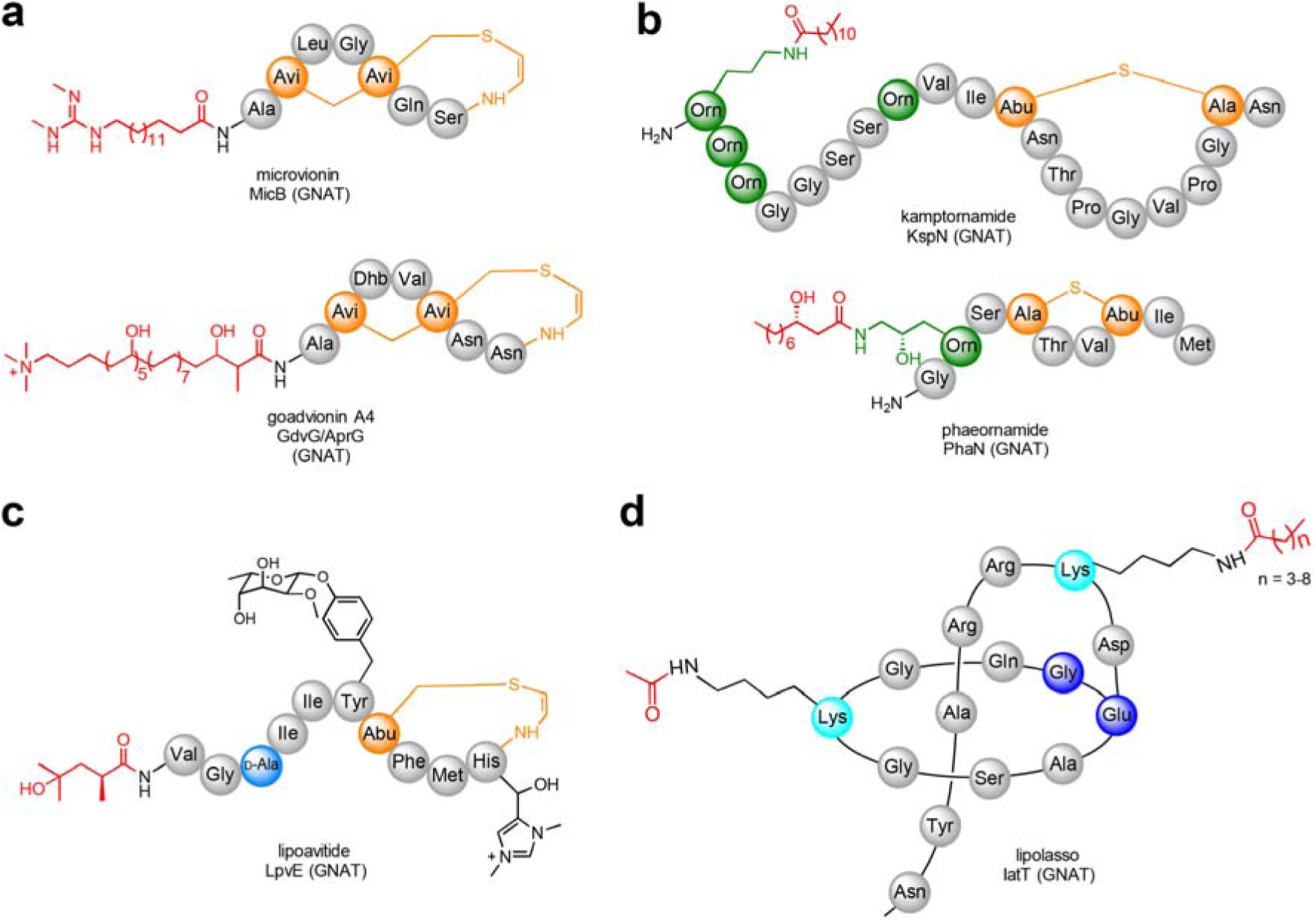
Structures of four classes of ribosomal lipopeptides. **a**, lipolanthines. **b**, selidamides. **c**, lipoavitides. **d**, lipolasso peptides. Avi, Avionin; Dhb, dehydrobutyrine; Orn, ornithine. Names of the compounds and enzymes for acyl group condensation with RiPPs are indicated below the structures.

In conclusion, we have characterized a large repertoire of GNATs capable of catalyzing iterative acetylation on mature lasso peptides instead of linear PPs. Our results not only provide insights into the molecular basis of such unprecedented catalytic feature, but also showcase how to enhance the efficiency of iterative catalysis and accelerate the divergent biosynthesis of ribosomal lipopeptides through the engineering of substrate binding pockets.

### Online content

Any methods, additional references, Nature Portfolio reporting summaries, source data, extended data, supplementary information, acknowledgements, peer review information; details of author contributions and competing interests; and statements of data and code availability are available at XXX.

## Methods

### General materials and methods

Oligonucleotides were synthesized by General Biol (Anhui) Co., Ltd. Restriction endonucleases were obtained from New England Biolabs and Thermo Fisher Scientific. Phanta Max Super-Fidelity DNA Polymerase and ClonExpress Ultra One Step Cloning Kit V2 were obtained from Vazyme. Amicon Ultra-4 centrifugal filters were purchased from EMD Millipore. Chemical reagents were purchased from Sigma Aldrich. All reagents and solvents were commercially available and used without further purification unless otherwise stated. Synthetic peptides were obtained from Nanjing Yuanpeptide Biotech Co., Ltd. (Nanjing, Jiangsu, China). *Escherichia coli* DH5α and *E. coli* BL21(DE3) cells were used for plasmid maintenance and protein overproduction, respectively. DNA sequencing was performed by Xi’an Qingke Biotechnology Co., Ltd. pET N-terminal His6 TEV LIC, pET His6 SUMO TEV LIC, and pET His6 MBP TEV LIC vectors were gifts from Scott Gradia. All NMR data was acquired on a Bruker AVANCE NEO 600 spectrometer, with chemical shifts referenced to the solvent peak of DMSO-*d*_6_ (Shanghai Haohong Scientific Co., Ltd.) at 2.50 ppm for ^1^H NMR analysis.

### Molecular biology techniques

The *embT*, *nocT*, *jiaT*, and *alkT* genes were codon optimized and synthesized by Xi’an Qingke Biotechnology Co., Ltd. Other genes were cloned from genomic DNA of the corresponding strains. DNA fragments of target genes were PCR-amplified using Phanta Max Super-Fidelity DNA Polymerase. Amplification products were confirmed by 1.2% agarose gel electrophoresis and purified using spin columns. The vectors were digested with the indicated restriction enzymes for 3 h in a 37 °C water bath. The resulting DNA products and linearized vector were assembled using the Ready-to-Use Seamless Cloning Kit. *E. coli* DH5α cells were transformed with 5 μL of the assembled products by heat shock, and cells were plated on LB agar plates supplemented with appropriate antibiotics and grown overnight at 37 °C. Several colonies were picked and used to inoculate separate 5 mL cultures of LB medium. The cultures were grown at 37 °C for 16 h before plasmid extraction. Mutagenesis was performed using the PCR-based seamless cloning method. All constructs were verified by DNA sequencing. The protein information is shown in Supplementary Table S1. The primers used in this study are listed in Supplementary Table S2. All plasmids used in this study were listed in Supplementary Table S3.

### Bioinformatics

The SSN of IatT was generated using “Sequence BLAST” option in EFI-EST Tools^25,43–45^, with the IatT sequence as the query. Sequences with 100% identity were conflated into a single node, and other parameters were kept as default. The SSN was visualized using the organic layout within Cytoscape^46^. A 40% sequence identity threshold was applied to separate the IatT-like GNAT cluster, and the resulting SSN was subjected to EFI-GNT analysis. The resulting GNNs and GNDs revealed GNAT-encoding lasso peptide BGCs, which were further manually confirmed. All sequences of the IatA-like PPs and IatT-like GNATs were collected and organized into FASTA files. These sequences were input into Mega11^47^ for sequence alignment using ClustalW^48^ and ML tree analysis. Sequence conservation logo figures were produced by submitting sequence alignment results to the online WebLogo 3 tool^49^.

### Heterologous expression of *iat* and *ven* BGCs

All heterologous expression plasmids listed in Supplementary Table S3 were introduced into *S. coelicolor* M1154 through *E. coli* ET12567/pUZ8002-*Streptomyces* conjugation, resulting in the corresponding recombinant strains.^26^

The cultivation of M1154 strain seed cultures was all carried out in TSB media (5.95 g tryptone, 1.05 g peptone, 0.875 g glucose, 1.75 g NaCl, 0.875 g K_2_HPO_4_, 350 mL deionized H_2_O adjusted to pH 7.3) supplemented with 10 μg/mL of apramycin and 25 μg/mL nalidixic acid with a total volume of 3.5 L (10 ×350 mL). The cultures were agitated at 220 rpm and 30 °C for approximately 3 days. The obtained seed cultures of M1154-*iat*-related strains were then evenly used to inoculate freshly prepared ISP2 media (1.4g Yeast extract, 3.5 g Malt extract, 1.4g Dextrose, 350 mL deionized H_2_O adjusted to pH 7.3) with a total volume of 30 L (100 × 350 mL). For *ven* BGC, the M1154-*ven* strain was cultured using M15 media (10.5 g glucose, 0.35 g peptone, 1.75 g beef extract,1.75 g NaCl, 0.875 g CaCO_3_, 350 μL trace elements, 350 mL deionized H_2_O adjusted to pH 7.3) with a total volume of about 1 L (3 × 350 mL). The incubation lasted for 7 days at 220 rpm and 30 °C. The resultant cultures were harvested by centrifugation.

### Isolation of mirusins

After cultivation, the M1154-*iat* cultures were harvested via centrifugation at 6,000 rpm for 20 min. The cell pellets were soaked with methanol multiple times until no target signals were detected in MALDI-TOF MS analysis. The methanol extracts were concentrated under reduced pressure, resulting in 24 g of cell crude extract. The supernatant, after filtration, and the cell crude extract were subjected to a column filled with HP20 resins. A gradient of H_2_O/MeOH (100/0, 70/30, 50/50, 20/80, 0/100, *v*/*v*) was used to elute the target compounds, yielding six fractions (A1- A6). Compounds **2** and **3** were found in fraction A2, as revealed by MALDI-TOF MS analysis.

The A2 fraction underwent further purification using a reversed-phase silica gel column, eluted with H_2_O/MeOH (100/0, 92/8, 80/20, 70/30, 50/50, 20/80, 0/100, *v/v*), yielding seven fractions (B1-B7). Fractions B3-B4, containing **2** and **3**, were combined for further purification using a silica gel column (80-120 mesh) eluted with CH_2_Cl_2_/MeOH mixture (100/0, 90/10, 80/20, 70/30, 60/40, 50/50, 0/100, *v/v*) and H_2_O with 0.1% TEA, yielding eight fractions (C1-C8). Compounds **2** and **3** were detected in fractions C7 and C8, which were combined and concentrated under reduced pressure.

Further purification was achieved using Sephadex LH-20 column eluted with 50% H_2_O/MeOH. The resulting sample was finally purified by HPLC equipped with a C18 reversed-phase column (Shimadzu, C18, 10 mm × 250 mm, 5 μm) The elution condition of **2** was: 5%–20% acetonitrile in H_2_O with 0.1% trifluoroacetic acid for 30 min; flow rate of 3 mL/min; t_R_ = 22.6 min. The elution condition of **3** was: 5%–25% acetonitrile in H_2_O with 0.1% trifluoroacetic acid for 30 min; flow rate of 3 mL/min; t_R_ = 20.1 min. Finally, 7.6 and 9.2 mg of **2** and **3** were purified from 21 L cultures, respectively.

Isolation of compound **1** from the M1154-*iat*-HM2 strain followed similar protocol. The corresponding cells were extracted with methanol to give 18 g sample, which was subjected to a column with HP20 resins and eluted using a gradient of H_2_O/MeOH (100/0, 90/10, 70/30, 50/50, 30/70, 10/90, 0/100, *v/v*), yielding seven fractions (D1-D7). All fractions were analyzed by LC-HRMS. Compound **1** was detected in fractions D5-D7, which were combined for further purification using Sephadex LH-20 column in MeOH. The resulting sample containing compound **1** was finally purified by HPLC equipped with an identical column (5%–50% acetonitrile in H_2_O with 0.1% formic acid for 30 min; 1 mL/min; t_R_=5 min). Finally, 2.4 mg of **1** was purified from 30 L cultures.

### Expression and purification of proteins

The plasmids containing the genes of interest were transformed into *E. coli* BL21(DE3) cells for protein expression. Cells were grown for 24 h on Luria-Bertani (LB) agar plates containing 50 µg/mL ampicillin or kanamycin at 37 °C. Single colonies were picked to inoculate 10 mL of LB containing appropriate antibiotic and grown overnight at 37 °C. This culture was used to inoculate 2 L of LB containing appropriate antibiotic, which was grown to an optical density at 600 nm (OD_600_) of 0.6-0.8, then placed on ice for 10 min. Isopropyl β-D-1- thiogalactopyranoside (IPTG) was added to a final concentration of 0.5 mM, followed by induction at 18 °C for 18 h. Cells were harvested via centrifugation at 6,000 rpm for 20 min and resuspended in 30 mL suspension buffer (500 mM NaCl, 20 mM Tris, pH 8.0, 10% glycerol (*v/v*)).

Cells were lysed by sonication (1s on, 2s off) at 50% amplitude in an ice water bath. Insoluble cell material was removed by centrifugation at 12,000 rpm for 30 min at 4 °C. The resultant supernatant was loaded onto a pre-equilibrated NiNTA-HisTalon μSphere column (GE; 5 mL). The column was washed with 40 mL wash buffer containing 1 M NaCl, 20 mM Tris, pH 8.0, and 30 mM imidazole. Elution was performed with 40 mL of elution buffer containing 1 M NaCl, 20 mM Tris, pH 8.0, and 250 mM imidazole, collecting 5ml fractions.

The resultant fractions were examined visually by Coomassie-stained sodium dodecyl sulfate polyacrylamide gel electrophoresis (SDS-PAGE) gel. A buffer exchange with protein storage buffer (20 mM Tris, pH 8.0, 300 mM NaCl, 10% glycerol (*v/v*), 0.5 mM β-mercaptoethanol) was performed using overnight dialysis in 4 °C fridge. For Cryo-EM structure determination and ITC experiments, IatT protein was further purified using a Superdex 200 size-exclusion column (GE Healthcare Lifesciences) with a buffer containing 300 mM KCl and 20 mM HEPES, pH 7.5. A single peak from the 280 nm absorbance trace, corresponding to the desired protein, was collected. The proteins were then concentrated using a 10 kDa molecular weight cut-off Amicon Ultra centrifugal filter. Protein concentrations were determined based on their absorbance at 280 nm, with theoretical extinction coefficients calculated using the ExPASy ProtParam tool (http://web.expasy.org/protparam/). Protein purity was visually inspected by Coomassie-stained SDS-PAGE (Supplementary Fig. 11).

### Expression and purification of linear IatA

The IatA expression plasmid (pET His6 SUMO TEV LIC vector) was transformed or co-transformed with the pRSFDuet-*iatB1*-*iatT* plasmid into *E. coli* BL21(DE3) for the expression of PP. The cultivation and purification processes were similar to those used for other proteins. Specifically, the cultures were shaken at 37 °C for 3 h for IatA expression and at 30 °C for 18 h for IatA-IatB1-IatT co-expression after induction by IPTG before cell harvest. Additionally, glycerol was removed from the storage buffer. The resulting SUMO-IatA solutions were treated with TEV for 14 hours in a 37 °C water bath to remove the SUMO tag. The cleaved SUMO was precipitated by heat treatment at 95 °C for 10 min. Subsequently, the supernatant was obtained through centrifugation and dried by lyophilization. The samples were then directly dissolved in methanol for in vitro IatT assays and LC-HRMS analysis.

### Carboxypeptidase Y and heat resistance

Although the peptides used in this study cannot be dissolved in water at high concentration for NMR analysis, they can be dissolved in a water-based buffer at concentration exceeding 100 μM. Therefore, linear ItaA at this concentration was treated with 0.5 units of carboxypeptidase Y in a solution containing 50 mM MES and 1.0 mM CaCl_2_ at pH 6.75 for 16 h at 25 °C. The same procedures were applied to lasso peptides **2** and **3** before and after incubation at 95 °C, as well as the branched cyclic peptide **2\***. The samples were then analyzed by MALDI-TOF MS or LC-HRMS.

### ITC

All ITC experiments were conducted using a MicroCal iTC200 instrument, featuring a cell volume of 200 μl and a 40-μl microsyringe. The experiments were carried out under the following conditions: cell temperature at 20 °C, stirring at 500 rpm, peptide concentration at 200 μM, IatT concentration at 20 μM, and an ITC buffer consisting of 300 mM KCl and 20 mM HEPES, pH 7.5. The system was equilibrated to 20L°C, with an additional delay of 60 s applied. The titrations began with an initial control injection of 0.4 μl, followed by 18 identical injections of 2 μl, with a duration of 0.8 s per injection and 180 s intervals between injections. The reference power was set at

1.2 μcal/s. Background data from the buffer sample were subtracted prior to data analysis. The data

were analyzed using the MicroCaliTC200 Origin7 analysis software package provided by the manufacturer.

### Cryo-EM sample preparation, data collection, and data processing

A 4 μl aliquot of the IatT- CoA (0.75 mg/mL) and IatT-AcCoA (1.00 mg/mL) sample was applied to a glow-discharged Quantifoil grid (Au 2/2, 200 mesh). The grid was blotted and plunge-frozen in liquid ethane using a Vitrobot mark IV (Thermo Fisher, Inc.) at 4 °C and 100% relative humidity.

Cryo-EM data were collected at 300 kV with a Titan Krios G3i (Thermo Fisher, Inc.) electron microscope equipped with a K3 direct electron detector with a Bio-Quantum energy filter (Gatan, Inc.). The microscope was operated at a nominal magnification of ×105,000 (pixel size 0.83 Å) with the detector in super-resolution mode. Each movie (40 frames) was acquired using a total exposure time of 3 s and a total dose of 60Le^−^/Å^2^.

Micrograph movies were imported into cryoSPARC4.0^50^ for image analysis. Imported movies were motion-corrected and dose-weighted. Contrast transfer function (CTF) estimation was performed using CTFFIND4^51^. A total of 13802 and 8723 micrographs with good CTF and astigmatism were selected for IatT-CoA and IatT-AcCoA, respectively. Approximately 500 particles were manually picked and subjected to 2D classification, with the best 2D classifications selected as templates for particle picking. A total of ∼6,240,000 particles of IatT-CoA and ∼5,190,000 particles of IatT-AcCoA were automatically picked and then subjected to multiple rounds of 2D and 3D classification. Finally, ∼1,440,000 particles of IatT-CoA and ∼4,350,000 particles of IatT-AcCoA corresponding to the best 3D class were subjected to multiple rounds of 3D auto-refinement with D7 symmetry applied. For all reconstructions, the resolution was assessed using the gold standard FSC 0.143 criterion. The image processing workflows for the two data sets are summarized in Extended Data Fig. 4.

### Model building and refinement

The predicted structure of IatT, generated using AlphaFold2^52^, was rigid-body fitted into cryo-EM density maps using UCSF Chimera^53^. The coordinates of CoA and AcCoA were derived from the structure of PseH-AcCoA (PDB code 4RI1)^54^. The two ligands were fitted into the density maps using Coot^55^, with geometry restraints generated using eLBOW of Phenix^56^. Both IatT-CoA and IatT-AcCoA models were refined with Phenix^56^ and then manually corrected with Coot^55^. The final models were assessed using MolProbity^57^. The visualization and graphic representation of the structures were generated using UCSF Chimera^53^. The data collection and refinement statistics are summarized in Supplementary Table 5.

### In vitro assays of GNATs

The complete reactions were performed in a 50 μl × 4 reaction system containing 50 μM peptide, 2 mM TCEP, 3 mM AcCoA, 50 mM Tris (pH 8.0), 100 mM NaCl, 2 mM MgCl_2_, and 10 μM IatT or its mutants. The reactions were incubated at 30 °C water bath overnight. Control reactions used boiled IatT or its mutants. All reactions were terminated by adding acetonitrile with 0.1% formic acid in a 1:1 (*v*/*v*) ratio. Subsequently, the reaction mixtures were centrifugated at 12,000 rpm for 1 min. The resulting supernatants were dried using a centrifugal concentrator and then re-dissolved in 100 μL of methanol for MAIDL-TOF MS or LC-HRMS analysis.

### LC-HRMS data acquisition

All LC-HRMS analysis was carried out using a ThermoFisher Scientific Q Exactive mass spectrometer. Separation was performed on an AcclaimTM 120 C18 3μm 120Å (3*150mm) column at a flow rate of 0.3 mL/min. The mobile phases used were solvent A (0.1% formic acid in water) and solvent B (0.1% formic acid in methanol). The following gradient was used for all in vitro experiments: 30-100% B over 15 min. For heterologous expression of *iat*, *ven*, and *iat*- HM2 BGCs, gradients were set to 5-100% B over 12 min, 10min, and 15 min respectively. The mass spectrometry instrument was configured to operate in positive mode, with a mass range of 200-3000 m/z. For MS/MS analysis, a Bruker maXis plus QTOF mass spectrometer was used, with the collision energy set to 55 eV.

### Lasso peptide modelling

Lasso-HTP^37^ was utilized to model the structure and topology of lasso peptide **1** with free Lys4 and Lys10. To generate the structure and topology model of the monoacetylated lasso peptide **2**, a three-step process was employed: 1) the AMBER parameters for acetylated Lys, referred to as ACK, were obtained from an open-source database (http://pc164.materials.uoi.gr/dpapageo/amberparams.php); 2) the name of Lys10 in the coordinate file of lasso peptide **1** was changed to ACK; 3) the AMBER parameters of ACK was added to the tLeap script of lasso peptide **1** generated by Lasso-HTP. Subsequent operation of the tLeap program in AmberTools23^58^ produced the structure and topology model of lasso peptide **2**. Conformation sampling was conducted using GROMACS 2023.4^59^. Each modelled lasso peptide was solvated in a cubic box containing approximately 1000 water molecules, and sodium chloride was used to neutralize the system. A steepest decent energy minimization was performed prior to the replica exchange molecular dynamics (REMD) simulation. The REMD simulation consisted of a 200 ps equilibration and a 2 ns production run in an isothermal-isobaric ensemble (NPT ensemble). Temperature coupling was achieved using the V-rescale method, while Berendsen and Parrinello-Rahman methods were used for pressure coupling during equilibration and production, respectively. Temperature points were set between 295K and 350K, resulting in replica exchange probabilities of approximately 20%. Trajectories at 300K were used for clustering based on heavy atoms, and the middle conformations of each cluster were selected as the final lasso peptide models.

### Molecule docking

All modelled conformations of lasso peptides **1** and **2** were used in the molecule docking with MEGADOCK 4.0^60^. The IatT residues at positions 5-15, 75-82 and 180-193, located on the periphery surface, were blocked from docking to enrich the correct docking conformation within the lasso peptide binding pocket. Docking conformations were initially filtered based on the distances between the ε-amine of the corresponding Lys in the lasso peptides and the carbonyl carbon of the acetyl group in AcCoA, which was set to be no more than 5.5 Å. The angle deviation of nucleophilic substitution direction was compared with the theorical Bürgi–Dunitz angle and required to be less than 36 °C. Filtered conformations were then populated by rotation according to the C2 symmetry of the IatT dimmer. Finally, the pairwise overlapping volume of all remaining lasso peptide conformations was estimated, and pairs without overlapping were collected. The final 49 docking coordinate files of lasso peptides **1** and/or **2** are provided as source data.

### In situ energy minimization

For each of the collected docking conformations containing two lasso peptide molecules, the topology models of the complexes were created using the tLeap program. An energy minimization was then performed using the sander module in AmberTools23^58^. During energy minimization, the backbone atoms of IatT and the carbonyl carbon in AcCoA were strongly restrained

with a restraint weight of 100. The GBneck2 model (igb=8) was employed as the implicit solvent model. The coordinates obtained after energy minimization were used for further analysis.

## Reporting summary

Further information on research design is available in the Nature Portfolio Reporting Summary linked to this article.

## Data availability

The data that support the findings of this study are available within the main text and its Supplementary Information files. Data are also available from the corresponding author upon request. Source data are provided with this paper. The cryo-EM maps of IatT in complex with CoA and AcCoA have been deposited into the EM Database with accession code EMD-60983 and EMD-60984, respectively. The corresponding coordinates have been deposited into the Protein Data Bank with accession code 9IY3 and 9IY4, respectively.

## Acknowledgments

This work was supported by the National Natural Science Foundation of China (No. 22077056, 22377046, 32171300, 22107040, and 21907046) and the Science and Technology Major Program of Gansu Province of China (22ZD6FA006 and 23ZDFA015).

## Author contributions

J.X. and S.-H.D. carried out bioinformatic, genetic and biochemical work and performed metabolic analysis, compound isolation and structure elucidation. S.W. and D.L. carried out cryo-EM data collection and structure elucidation. Z.-Q.L. started the project and performed heterologous expression and isolation of mirusins. X.-T.G., S.F., F.-Y.T., and J.-J.C. participated in the heterologous expression and cloning experiments. S.L. and S.-H.D. designed, conceived, and supervised the project. S.L., D.L., and S.-H.D. analyzed data and wrote the manuscript with input from all authors.

## Additional information

Extended data is available for this paper at XXX.

## Supplementary information

The online version contains supplementary material available at XXX.

## Correspondence and requests for materials

should be addressed to Shangwen Luo, Dongsheng Lei, or Shi-Hui Dong.

---

**Extended Data Fig. 1.**
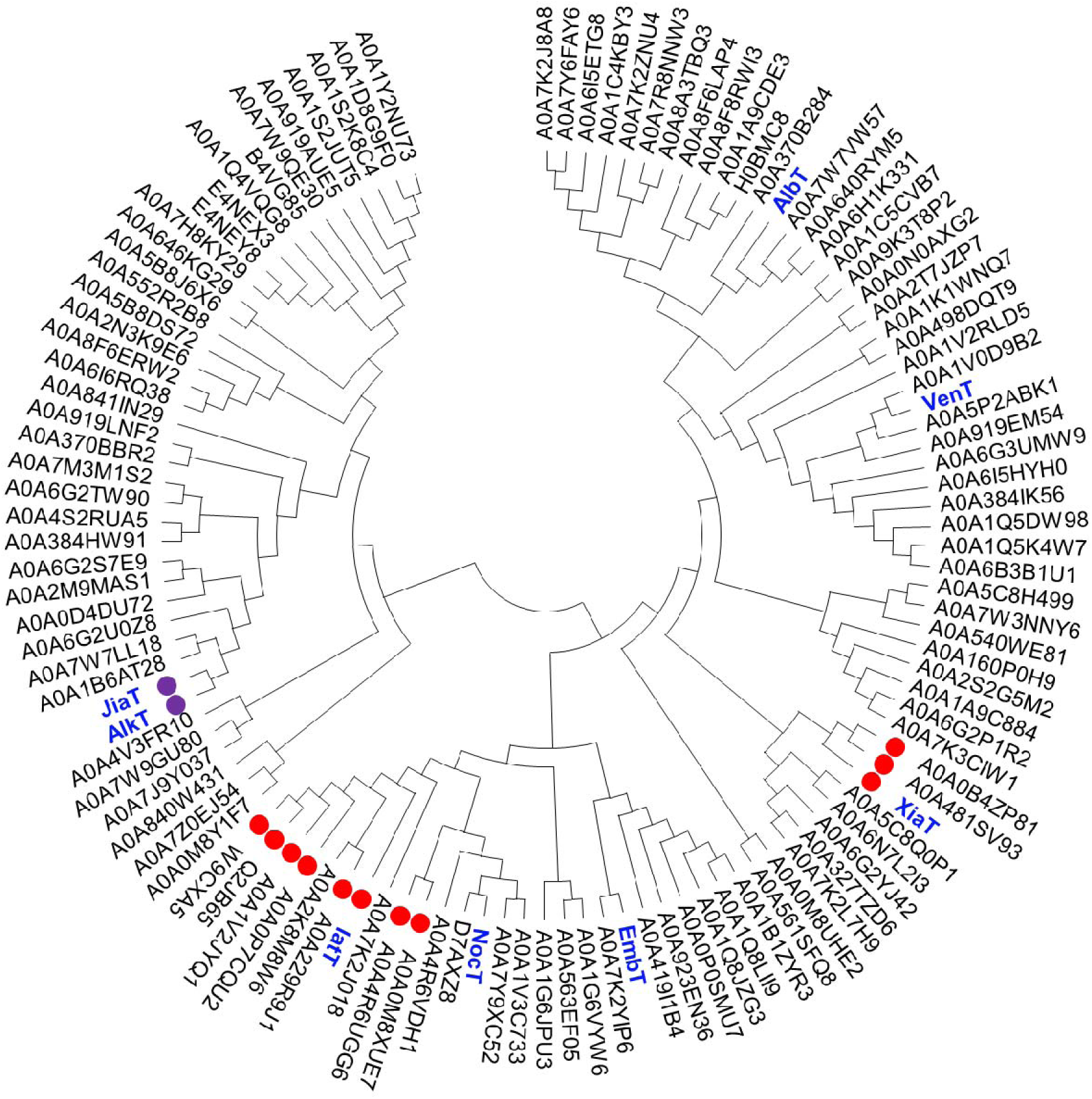
The ML tree of lasso peptide GNATs. The class II members are indicated by solid red circles, whereas the two exceptions with a Lys5 instead of Lys4 are denoted by purple circles. The characterized lasso peptide GNATs are displayed in blue bold names. Other IatT-like GNATs are indicated by their UniProt IDs.

**Extended Data Fig. 2.**
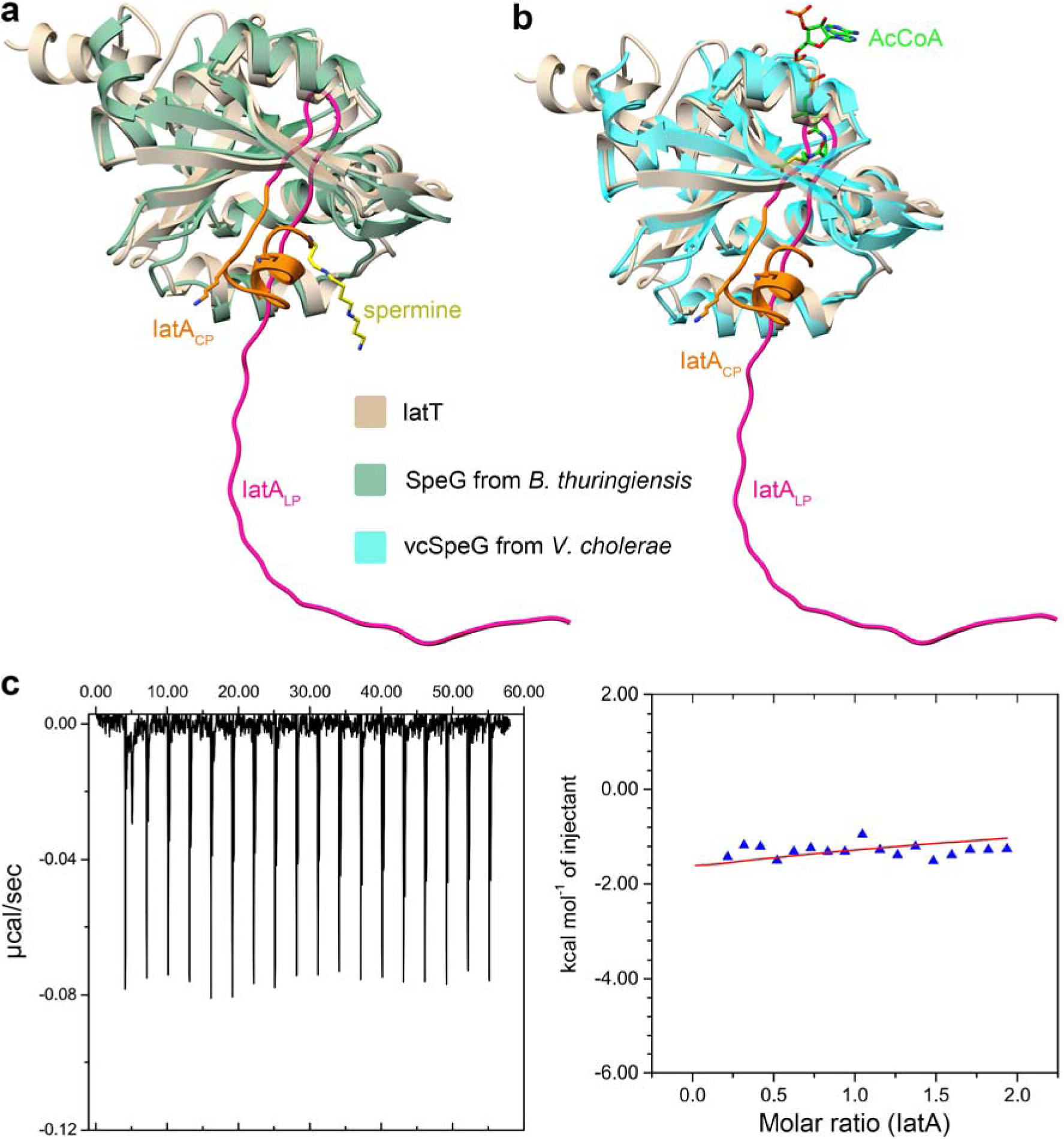
Binding analysis of IatT-IatA complex. **a**,**b**, Structural comparison of IatT- IatA AlphaFold-multimer model with SpeG in complex with spermine related products (**a**) and PseH bound with AcCoA (**b**). **c**, ITC data for titration of IatA into IatT solution. No obvious binding is observed. All assays were run in triplicate and representative results are shown.

**Extended Data Fig. 3.**
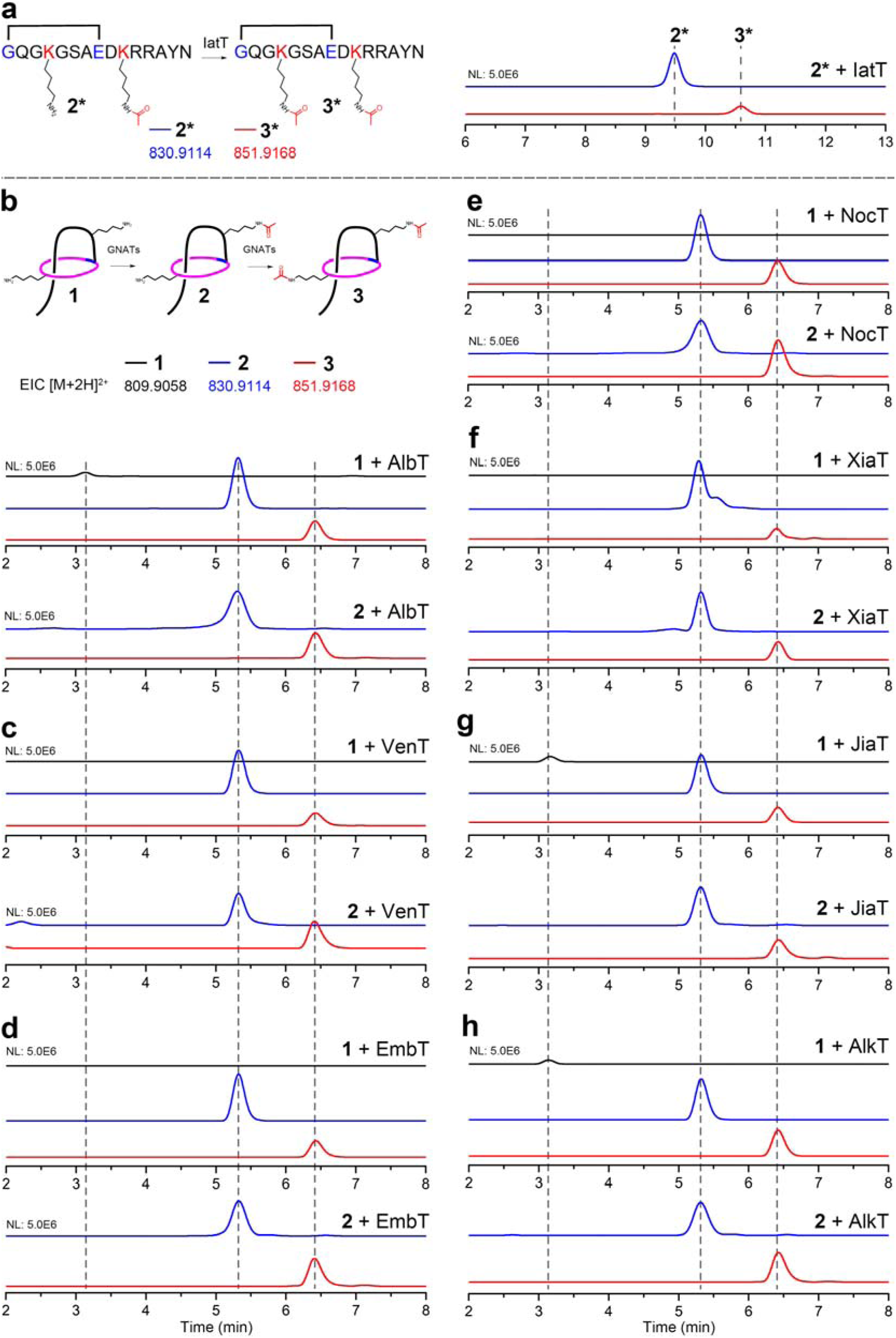
Enzymatic activity assays of IatT and homologous GNATs in vitro via LC-HRMS analysis. **a**, EICs for assays of IatT using **2\*** as substrate. **b**-**h**, EICs for assays of AlbT (**b**), VenT (**c**), EmbT (**d**), NocT (**e**), XiaT (**f**), JiaT (**g**), and AlkT (**h**) using 1 and 2 as substrates. All assays were run in triplicate and representative results are shown.

**Extended Data Fig. 4.**
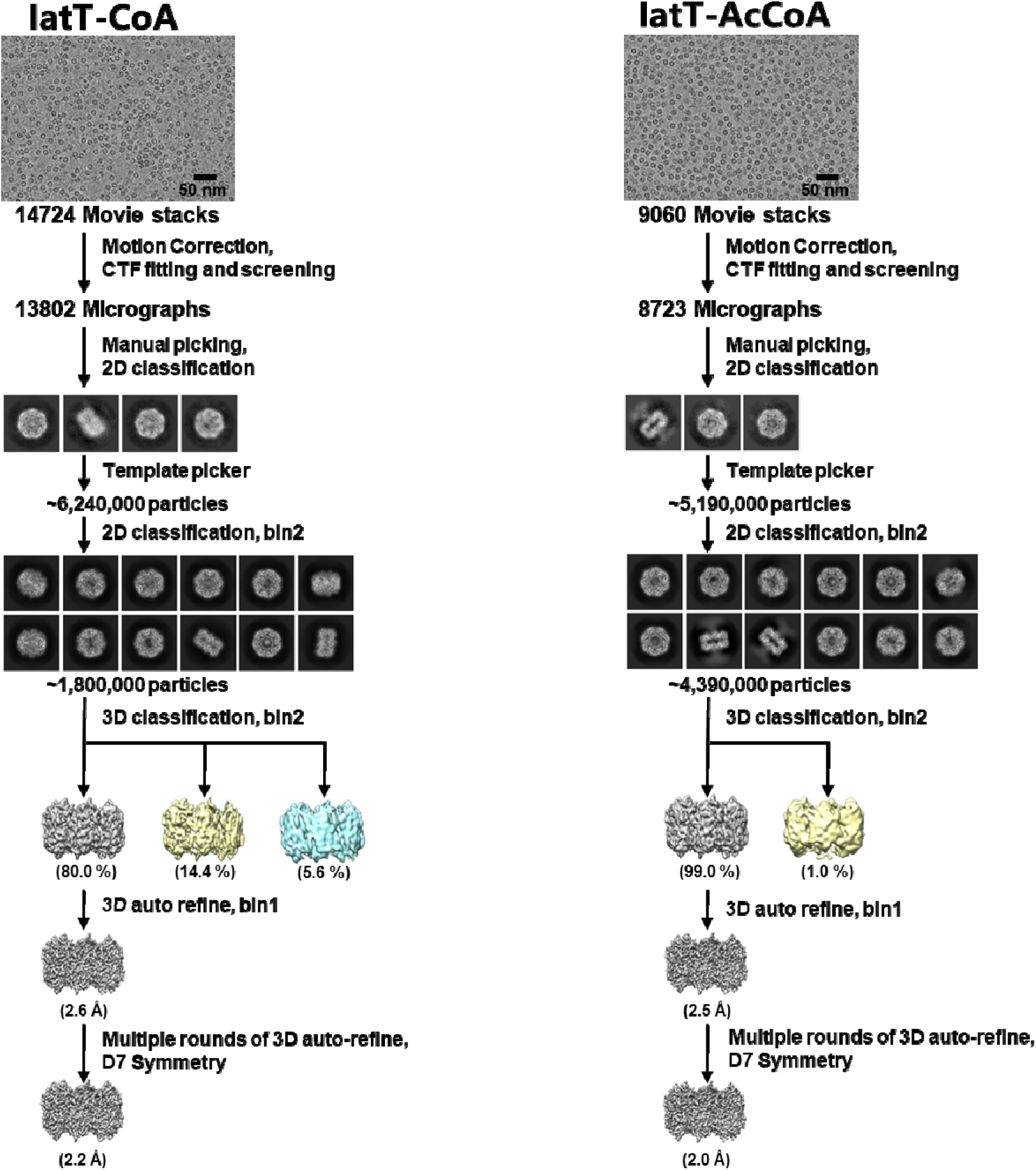
Flow-chart for cryo-EM data processing. **a**,**b**, Shown the data processing procedures for entire protein complex of IatT-CoA (**a**) and IatT-AcCoA (**b**). Details can be found in the Methods section.

**Extended Data Fig. 5.**
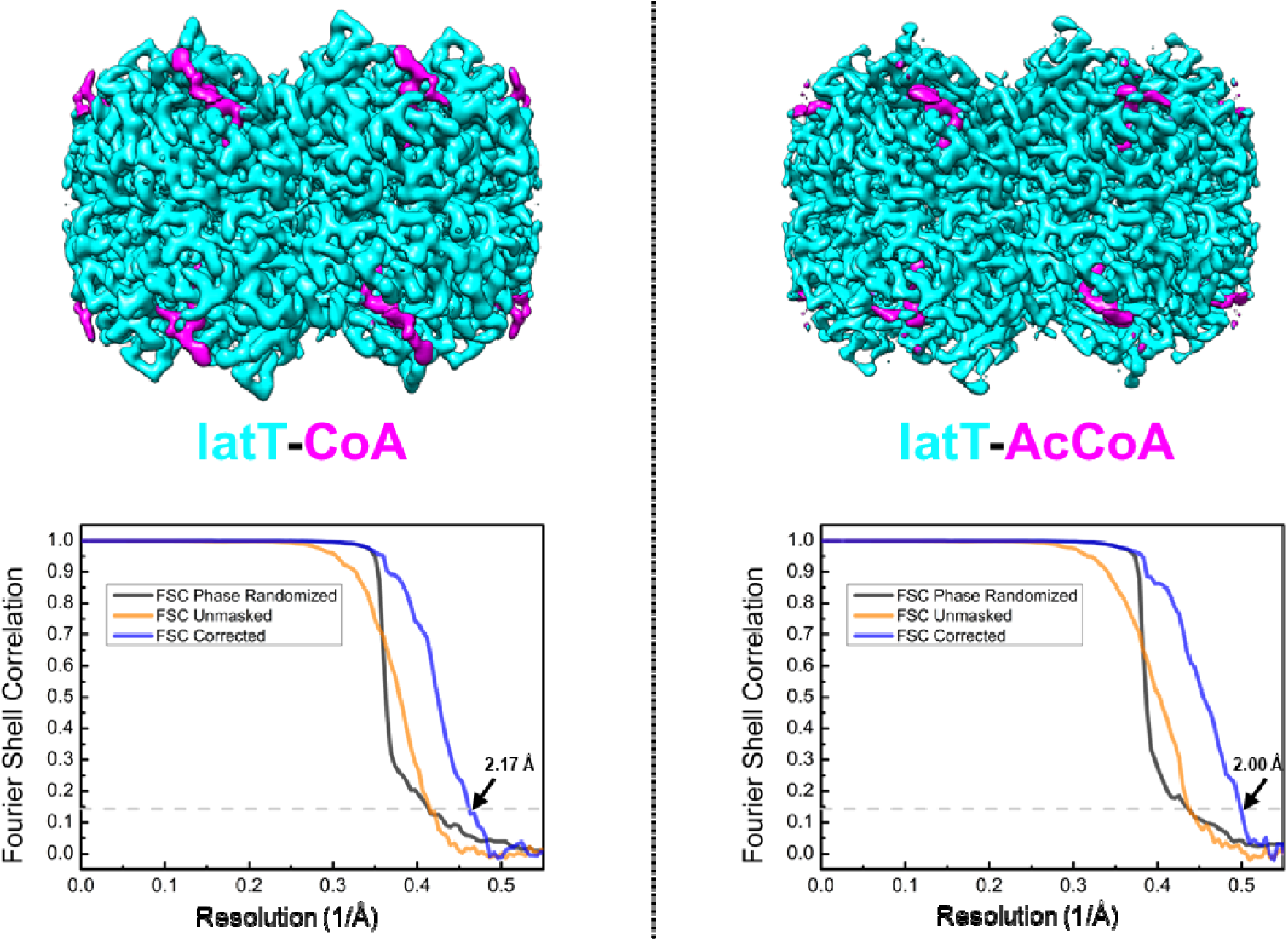
Resolution analysis of cryo-EM 3D reconstruction. Top, overall density maps of the final 3D reconstruction; bottom, the gold-standard Fourier shell correlation (FSC) curves for the 3D reconstruction calculated in cryoSPARC. FSCLJ=LJ0.143 is indicated.

**Extended Data Fig. 6.**
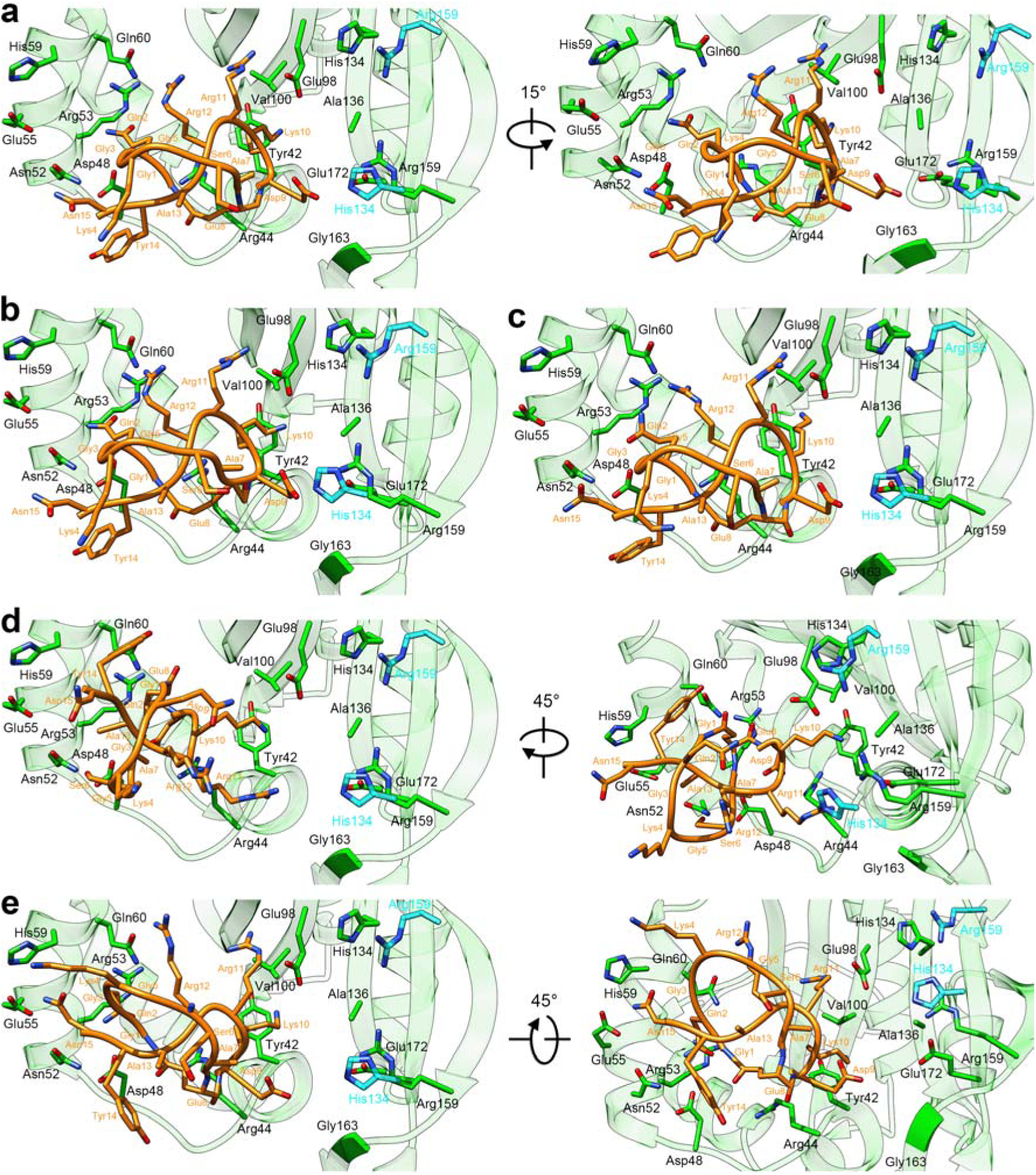
Conformations 1a (a), 1b (b), 1c (c), 1d (d), and 1e (e) of lasso peptide 1 in the IatT pocket. Conformation **1a** shown here is identical with that in Fig. 6a with slight movement for comparison purpose. The color codes are the same with Fig. 4f.

**Extended Data Fig. 7.**
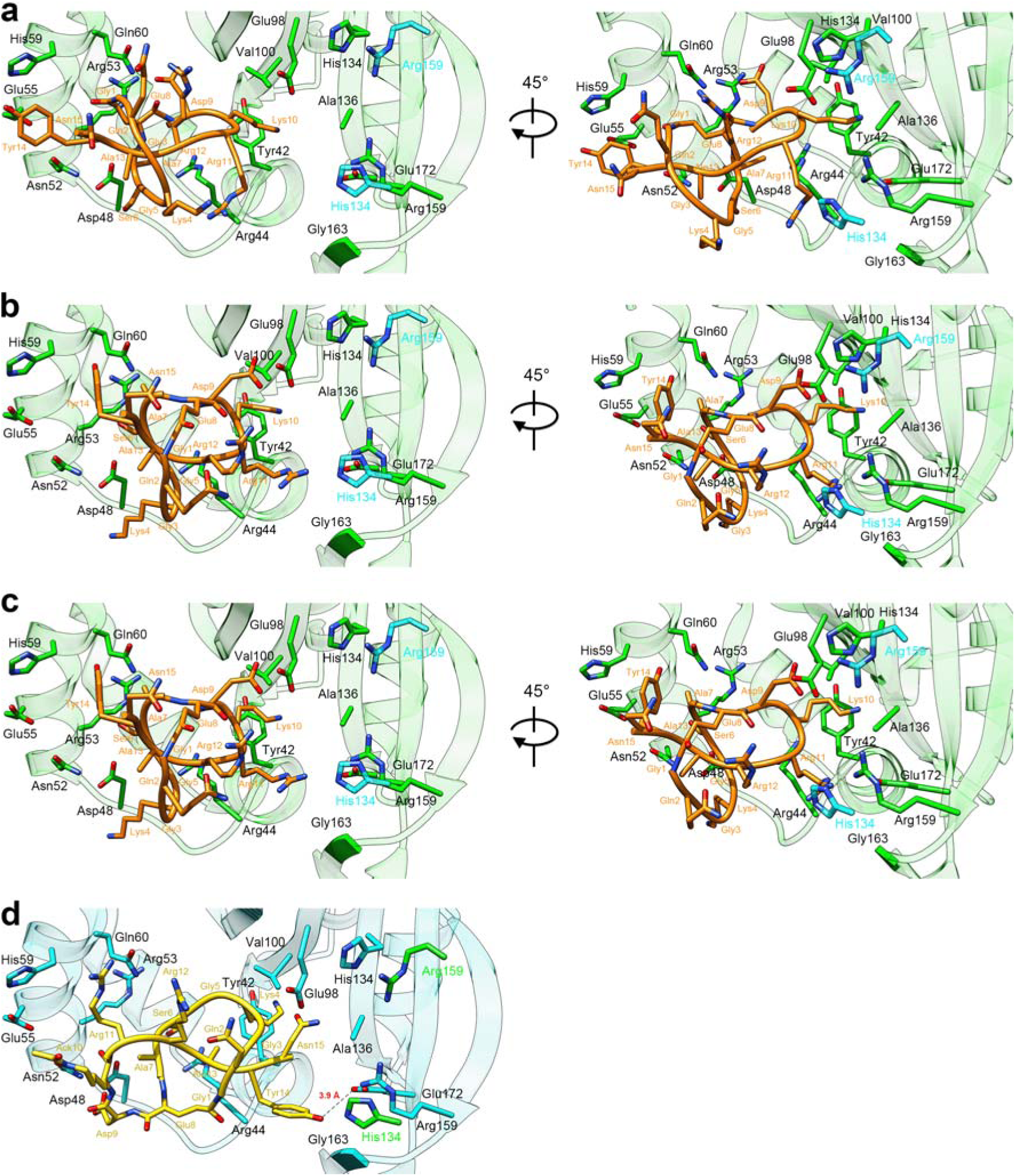
Conformations 1f (a), 1g (b), and 1h (c) of lasso peptide 1 and a different view of the conformation of lasso peptide 2 (d) in the IatT pocket. The color codes are the same with Figs. 4f and 4g.

