## Supplementary File for "Iterative acylation on mature lasso peptides by widespread acetyltransferases for lipolasso production"

^#^Denotes equal authorship contribution.

ORCIDs:

SL: 0000-0003-2162-2477;

DL: 0000-0003-1640-0693

SD: 0000-0002-1743-2163;

**Table of Contents:**

Table 1: Detailed information of the proteins used in this study	S4

Table 2: Oligonucleotide primers used in this study	S5

Table 3: Plasmids used in this study	S8

Table 4: ^1^H NMR data of **2***	S9

Table 5: Cryo-EM data collection and refinement statistics	S10

Table 6: Turnover ratios of IatT and its variants using **1** and **2** as substrates	S11

Fig. 1: Bioinformatic analysis of GNAT-coding lasso peptide BGCs	S12

Fig. 2: MS analysis of heterologous expression products of *iat* BGC	S13

Fig. 3: Comparison of ^1^H NMR spectra of **2***, **2**, and **3**	S14

Fig. 4: ^1^H-^1^H COSY NMR spectra of **2** and **3**	S15

Fig. 5: HRMS/MS analysis of **2***	S16

Fig. 6: ^1^H-^1^H COSY NMR spectra of **2***	S17

Fig. 7: Confirmation of the presence of lasso structure in **2** and **3**	S18

Fig. 8: HRMS/MS analysis of **2**	S19

Fig. 9: HRMS/MS analysis of **3**	S20

Fig. 10: LC-HRMS analysis of heterologous expression products of *ven* BGC containing *venABCT*, *venABC*, and *venA(G4K)BCT* genes	S21

Fig. 11: SDS-PAGE analysis of purified proteins	S22

Fig. 12: LC-HRMS analysis of the TEV-digested SUMO-IatA	S23

Fig. 13: LC-HRMS analysis of heterologous expression products of *iat*-HM1 BGC	S24

Fig. 14: DALI search results of IatT-AcCoA cryo-EM structure	S25

Fig. 15: Binding modes of CoA and AcoA with IatT	S26

Fig. 16: Sequence logo plot of the 112 IatT-like GNATs	S27

Fig. 17: Enzymatic activity assays of IatT-A136T, A136L, and A136E variants in vitro via LC-HRMS analysis	S28

Fig. 18: Enzymatic activity assays of IatT-Y42A, D48A, N52A, R53A, R53W, H59A, Q60A, E98A, V100W, H134A, H134Y, and R159A variants in vitro via LC-HRMS analysis	S29

Fig. 19: Enzymatic activity assays of IatT-R44A, E55A, A136Y, G163F, and E172A variants in vitro via LC-HRMS analysis	S30

Fig. 20: Enzymatic activity assays of IatT-Y42W, A136H, and A136K variants in vitro via LC-HRMS analysis	S31

Fig. 21: Enzymatic activity assays of IatT-E98W and E172W variants in vitro via LC-HRMS analysis	S32

Fig. 22: HRMS/MS analysis of **8**	S33

Fig. 23: HRMS/MS analysis of **9**	S34

Fig. 24: HRMS/MS analysis of **10**	S35

Fig. 25: HRMS/MS analysis of **17**	S36

Fig. 26: HRMS/MS analysis of **18**	S37

Fig. 27: HRMS/MS analysis of **10**	S38

**Supplementary Table 1. Detailed information of the proteins used in this study.**

| **name** | **annotation** | **UniProt ID** |
| --- | --- | --- |
| IatA | precursor peptide (PP), class II | C6WJ42 |
| IatB | RRE and trans-glutaminase | C6WJ44 |
| IatC | lasso cyclase | C6WJ43 |
| IatT | GCN5-related N-acetyltransferase (GNAT), class II | C6WJ45 |
| VenA | PP | F2R8U3 |
| VenB | RRE and trans-glutaminase | F2R8U5 |
| VenC | lasso cyclase | F2R8U4 |
| VenT | GNAT, class I | F2R8U6 |
| AlbT | GNAT, class I | A0A0B5EM53 |
| EmbT | GNAT, class I | A0A1T3NW30 |
| NocT | GNAT, class I | A0A7X6RMX2 |
| XiaT | GNAT, class II | A0A0F7FZ09 |
| JiaA | PP, class II | A0A1H5PE46 |
| JiaT | GNAT, class II | A0A1H5PGQ2 |
| AlkA | PP, class II | A0A1H2GJP3 |
| AlkT | GNAT, class II | A0A1H2GJZ4 |

**Supplementary Table 2. Oligonucleotide primers used in this study.** Nucleotide sequences are given in the 5’ to 3’ direction. F, forward primer; R, reverse primer; capital letters, homologous sequence with vector or mutagenized codon.

| **Primer name** | **Oligonucleotide sequence** |
| --- | --- |
| iat-KasO-F | CTGCATGCATACGTACTAGTCTGACAcacgacaggtccagctacccag |
| iat-KasO-R | CACAGGAAACAGCTATGACATGATTACGtcaccgcccaccctccacgaac |
| iat-KasO-R2 | CACAGGAAACAGCTATGACATGATTACGtcattgcctgatgtcttcccctc |
| IatT-LIC-F | CCTGTACTTCCAATCCAATGCAatgagccctcaagacgacctgatc |
| IatT-LIC-R | GATCCGTTATCCACTTCCAATGTTATTAccgcccaccctccacgaac |
| IatA-LIC-F | CCTGTACTTCCAATCCAATGCAatggcgcttctgccggaacagtg |
| IatA-LIC-R | GATCCGTTATCCACTTCCAATGTTATTAgttgtaggcccggcgcttgt |
| IatT-NdeI-F | GTTAAGTATAAGAAGGAGATATACatgagccctcaagacgacctg |
| IatT-NdeI-R | GATATCCAATTGAGATCTGCCAttaccgcccaccctccacgaacc |
| IatB1codon-LICF | CCTGTACTTCCAATCCAATGCAatgaaaccgacttgccgtgttgtggt |
| IatB1codon-LICR | GATCCGTTATCCACTTCCAATGTTATTAccaggacggttcccacggcggacca |
| IatB1codon-NcoIF | GTTTAACTTTAATAAGGAGATATACatgaaaccgacttgccgtgttgtggt |
| IatB1codon-NcoIR | GATGATGGTGATGGCTGCTGCCTTAccaggacggttcccacggcggacca |
| iat-KasO HpaI-R2 | CACAGGAAACAGCTATGACATGATTACGttaacgagtccggtgtctggcggccgtaa |
| iat-KasO HpaI-F2 | acggccgccagacaccggactcgttggagaac |
| IatT-T86Q-R | gatcagcactgcCTGggttccgaccggtaccggatc |
| IatT-T86Q-F  IatT -T86E-R | ccggtcggaaccCAGgcagtgctgatcgaccaccac  gatcagcactgcCTCggttccgaccggtaccggatc |
| IatT-T86E-F | ccggtcggaaccGAGgcagtgctgatcgaccaccac |
| IatT-T86A-R | gatcagcactgcTGCggttccgaccggtaccggatc |
| IatT-T86A-F | ccggtcggaaccGCAgcagtgctgatcgaccaccac |
| IatT-F99S-I101S-R | cgccgagctgAGAcacAGactcgccggtgcgcacgtggtg |
| IatT-F99S-I101S-F | caccggcgagtCTgtgTCTcagctcggcgcgggacaccgg |
| IatT-F99A-I101A-R | cgccgagctgAGCcacAGCctcgccggtgcgcacgtggtg |
| IatT-F99A-I101A-F | caccggcgagGCTgtgGCTcagctcggcgcgggacaccgg |
| IatT-T117R-R | gagcgtcaaccgCCtcgcctcggtgcccagccccct |
| IatT-T117R-F | caccgaggcgaGGcggttgacgctcgactacgcgtt |
| IatT-T117A-R | gagcgtcaaccgAgCcgcctcggtgcccagccccct |
| IatT-T117A-F | caccgaggcgGcTcggttgacgctcgactacgcgtt |
| IatT-L135S-R | cgtgagcaccgcAGAgtgcacgcaggcgagcgcgga |
| IatT-L135S-F | gcctgcgtgcacTCTgcggtgctcacgcccaacacc |
| IatT-L135A-R | cgtgagcaccgcAGCgtgcacgcaggcgagcgcgga |
| IatT-L135A-F | gcctgcgtgcacGCTgcggtgctcacgcccaacacc |
| IatT-Y148S-R | cccggcccgctcAGacgccgcgatagccccggtgtt |
| IatT-Y148S-F | gctatcgcggcgtCTgagcgggccgggtttcgcagg |
| IatT-Y148A-R | cccggcccgctcAGCcgccgcgatagccccggtgtt |
| IatT-Y148A-F | gctatcgcggcgGCTgagcgggccgggtttcgcagg |
| XiaB1NcoI-F | GTTTAACTTTAATAAGGAGATATACatgtctgatccaccggctcatgcta |
| XiaB1NcoI-R | GATGATGGTGATGGCTGCTGCCTTAacgagccagtgctgcaccagcacg |
| XiaTNdeI-F | GTTAAGTATAAGAAGGAGATATACatgattggttacggtcgtcaggcac |
| XiaTNdeI-R | GATATCCAATTGAGATCTGCCATTAagacggctgcgccggatcggagata |
| XiaB1-LIC-F | AAAACCTGTACTTCCAATCCAATGCAatgtctgatccaccggctcatgcta |
| XiaB1-LIC-R | GATCCGTTATCCACTTCCAATGTTATTAacgagccagtgctgcaccagcacg |
| XiaXiaT-LIC-F | AAAACCTGTACTTCCAATCCAATGCAatgattggttacggtcgtcaggcac |
| XiaXia-LIC-R | GATCCGTTATCCACTTCCAATGTTATTAagacggctgcgccggatcggagata |
| IatT-Y42W-F | gtgctgatcggttGGggccgccagacaccggactcgttg |
| IatT-Y42W-R | tgtctggcggccCCaaccgatcagcacggaggggtc |
| IatT-H134Y-F | ctcgcctgcgtgTacctggcggtgctcacgcccaacac |
| IatT-H134Y-R | gagcaccgccaggtAcacgcaggcgagcgcggacacgt |
| IatT-A136H-F | tgcgtgcacctgCATgtgctcacgcccaacaccggggctat |
| IatT-A136H-R | gggcgtgagcacATGcaggtgcacgcaggcgagcgcgga |
| IatT-G163F-F | cggcgcgattccTTCttctggctgggccggcgagtcag |
| IatT-G163F-R | gcccagccagaaGAAggaatcgcgccgttccccgatc |
| IatT-E98A-F  IatT-E98A-R | gtgcgcaccggcgCAttcgtgatccagctcggcgcgggaca  ctggatcacgaaTGcgccggtgcgcacgtggtggtcgatc |
| IatT-E98W-R | gtgcgcaccggcTGGttcgtgatccagctcggcgcgggaca |
| IatT-E98W-R | ctggatcacgaacCAgccggtgcgcacgtggtggtcgatc |
| IatT-E172W-R1 | ggaaagtcctcgggcaccgcgtccatcaaggtcCAgctgactcgccggcccagccagaa |
| IatT-E172W-R2 | ccgcccaccctccacgaacccgcgcacgacagacggaccgggaaagtcctcgggcac |
| IatT-R159G-F | caggatcggggaaGGTcgcgattccggtttct |
| IatT-R159G-R | cagaaaccggaatcgcgACCttccccgatcctg |
| IatT-R159A-F | aggatcggggaaGCAcgcgattccggtttctggct |
| IatT-R159A-R | accggaatcgcgTGCttccccgatcctgcgaaac |
| IatT-E55A-F | gttggagaaccggcgtgCAgggtacggccaccag |
| IatT-E55A-R | ctgctggtggccgtacccTGcacgccggttctcca |
| IatT -D48A-F | ccagacaccggcAtcgttggagaaccggcgtga |
| IatT -D48A-R | tctccaacgaTgccggtgtctggcggccgtaac |
| IatT -N52A-F | tcgttggagGCacggcgtgaggggtacggccacca |
| IatT -N52A-R | cctcacgccgtGCctccaacgagtccggtgtct |
| IatT -R53A-F | gactcgttggagaacGCAcgtgaggggtacggc |
| IatT -R53A-R | gccgtacccctcacgTGCgttctccaacgagtc |
| IatT -Q60A-F | gtacggccacGcagctcgcggtaccgacgacca |
| IatT -Q60A-R | taccgcgagcctggtggccgtacccctcacgccggtt |
| IatT -H134A-F | cgcgctcgcctgcgtgGCActggcggtgctcacgc |
| IatT -H134A-R | gcgtgagcaccgccagTGCcacgcaggcgagcgcg |
| IatT -R44A-F | ctgatcggttacggcGCAcagacaccggactcgt |
| IatT -R44A-R | caacgagtccggtgtctgTGCgccgtaaccgatcag |
| IatT -E172A-F | ctgggccggcgagtcagcgCAaccttgatggacgcgg |
| IatT -E172A-R | caccgcgtccatcaaggtTGcgctgactcgccggcccag |
| IatT-H59A-F | gtgaggggtacggcGCAcaggctcgcggtaccga |
| IatT-H59A-R | tcggtaccgcgagcctgTGCgccgtacccctcac |
| IatT-R53W-F | cgttggagaacTggcgtgaggggtacggccaccag |
| IatT-R53W-R | tacccctcacgccAgttctccaacgagtccggtgt |
| IatT-V100W-F | gcaccggcgagttcTGgatccagctcggcgcg |
| IatT-V100W-R | cgcgccgagctggatcCAgaactcgccggtgc |
| IatT-A136Y-F | gcctgcgtgcacctgTACgtgctcacgcccaa |
| IatT-A136Y-R | gtgttgggcgtgagcacGTAcaggtgcacgca |
| IatT-A136T-F | gcctgcgtgcacctgAcggtgctcacgcccaa |
| IatT-A136T-R | gttgggcgtgagcaccgTcaggtgcacgcaggc |
| IatT-A136K-F | gcctgcgtgcacctgAAggtgctcacgcccaa |
| IatT-A136K-R | gttgggcgtgagcaccTTcaggtgcacgcaggc |
| IatT-A136E-F | gcctgcgtgcacctggAggtgctcacgcccaa |
| IatT-A136E-R | gttgggcgtgagcaccTccaggtgcacgcaggc |
| IatT-A136L-F | gcctgcgtgcacctgCTggtgctcacgcccaa |
| IatT-A136L-R | gttgggcgtgagcaccAGcaggtgcacgcaggc |
| IatT-Y42A-F | ctccgtgctgatcggtGCTggccgccagacac |
| IatT-Y42A-R | gtgtctggcggccAGCaccgatcagcacggag |
| VenT-LIC-F | AAAACCTGTACTTCCAATCCAATGCAatgaacgaccacgcccccgacct |
| VenT-LIC-R | GATCCGTTATCCACTTCCAATGTTATTActtctcagggaacagcgccttca |
| ven-KasO-F1 | CAGCGTGCAGGACTGGGGGAGTTCaatattgggcgcgtttcccgtccgagtc |
| ven-KasO-R1 | CACAGGAAACAGCTATGACATGATTACGttaAcgcgcaggtcgggctggagat |
| ven-KasO-R2 | CACAGGAAACAGCTATGACATGATTACGctacttctcagggaacagcgccttc |
| ven-KasO-F2 | CAGCGTGCAGGACTGGGGGAGTTCaatgaccacagagaggaaccgtcatga |
| ven-KasO-R3 | gactcggacgggaaacgcgcccaatcgccaagcggacgacctggtgagag |
| ven-KasO-F3 | gagctgacccagggccagggcAAGggccagagcgaggacaagcgg |
| ven-KasO-R4 | ccgcttgtcctcgctctggccCTTgccctggccctgggtcagctc |

**Supplementary Table 3.** **Plasmids used in this study.**

| **Gene** | **Vector** | **Cloning site** | **Use** |
| --- | --- | --- | --- |
| **Co-expression experiments** | | | |
| *iatA* | pET His6 SUMO TEV LIC | SspI | PP expression |
| *xiaA* | pET His6 MBP TEV LIC |  |  |
| *iatT* | pRSFDuet | NdeI | Acetylation PTM |
| *xiaT* | pRSFDuet |  |  |
| **Protein overproduction experiments** | | | |
| *iatT* | pET N-terminal His6 TEV LIC | SspI | IatT expression |
| *xiaT* | pET His6 MBP TEV LIC | SspI | MBP-XiaT expression |
| *venT* | pET N-terminal His6 TEV LIC | SspI | VenT expression |
| *alkT* | pET N-terminal His6 TEV LIC | SspI | AlkT expression |
| *nocT* | pET N-terminal His6 TEV LIC | SspI | NocT expression |
| *jiaT* | pET N-terminal His6 TEV LIC | SspI | JiaT expression |
| *albT* | pET N-terminal His6 TEV LIC | SspI | AlbT expression |
| *embT* | pET N-terminal His6 TEV LIC | SspI | EmbT expression |
| *iatT-A136T* | pET N-terminal His6 TEV LIC | SspI | IatT-A136T expression |
| *iatT-A136L* | pET N-terminal His6 TEV LIC | SspI | IatT-A136L expression |
| *iatT-A136E* | pET N-terminal His6 TEV LIC | SspI | IatT-A136E expression |
| *iatT-E55A* | pET N-terminal His6 TEV LIC | SspI | IatT-E55A expression |
| *iatT-A136Y* | pET N-terminal His6 TEV LIC | SspI | IatT-A136Y expression |
| *iatT-G163F* | pET N-terminal His6 TEV LIC | SspI | IatT-G163F expression |
| *iatT-Y42W* | pET N-terminal His6 TEV LIC | SspI | IatT-Y42W expression |
| *iatT-A136H* | pET N-terminal His6 TEV LIC | SspI | IatT-A136H expression |
| *iatT-A136K* | pET N-terminal His6 TEV LIC | SspI | IatT-A136K expression |
| *iatT-Y42A* | pET N-terminal His6 TEV LIC | SspI | IatT-Y42A expression |
| *iatT-R53A* | pET N-terminal His6 TEV LIC | SspI | IatT-R53A expression |
| *iatT-R53W* | pET N-terminal His6 TEV LIC | SspI | IatT-R53W expression |
| *iatT-E98A* | pET N-terminal His6 TEV LIC | SspI | IatT-E98A expression |
| *iatT-V100W* | pET N-terminal His6 TEV LIC | SspI | IatT-V100W expression |
| *iatT-H134A* | pET N-terminal His6 TEV LIC | SspI | IatT-H134A expression |
| *iatT-H134Y* | pET N-terminal His6 TEV LIC | SspI | IatT-H134Y expression |
| *iatT-R159A* | pET N-terminal His6 TEV LIC | SspI | IatT-R159A expression |
| *iatT-E98W* | pET N-terminal His6 TEV LIC | SspI | IatT-E98W expression |
| *iatT-E172W* | pET N-terminal His6 TEV LIC | SspI | IatT-E172W expression |
| *iatT-D48A* | pET N-terminal His6 TEV LIC | SspI | IatT-D48A expression |
| *iatT-N52A* | pET N-terminal His6 TEV LIC | SspI | IatT-N52A expression |
| *iatT-Q60A* | pET N-terminal His6 TEV LIC | SspI | IatT-Q60A expression |
| *iatT-H134A* | pET N-terminal His6 TEV LIC | SspI | IatT-H134A expression |
| *iatT-R44A* | pET N-terminal His6 TEV LIC | SspI | IatT-R44A expression |
| *iatT-E172A* | pET N-terminal His6 TEV LIC | SspI | IatT-E172A expression |
| **Heterologous expression experiments** | | | |
| *iatABC* | pSET-KasO | AflII/EcoRI | Heterologous expression |
| *iatABCT* | pSET-KasO | AflII/EcoRI |  |
| *iatABCT-HM1* | pSET-KasO | AflII/EcoRI |  |
| *iatABCT-HM2* | pSET-KasO | AflII/EcoRI |  |
| *venABC* | pSET-KasO | AflII/EcoRI |  |
| *venABCT* | pSET-KasO | AflII/EcoRI |  |
| *venA(G4K)BCT* | pSET-KasO | AflII/EcoRI |  |

**Supplementary Table 4.** **^1^H NMR data of 2*.**

| Residue | NH | *α*H | *β*H | *γ*H | *δ*H | *ε*H | Ac |
| --- | --- | --- | --- | --- | --- | --- | --- |
| Gly^1^ | 8.379 | 3.948, 3.579 |  |  |  |  |  |
| Gln^2^ | 7.717 | 3.363 | 2.728 |  | NH 7.163 |  |  |
| Gly^3,5^ | 8.303 | 3.999, 3.541 |  |  |  |  |  |
| Lys^4^ | 8.567 | 3.942 | 1.953 | - | 1.800 | 2.105 |  |
| Ser^6^ | 8.494 | 5.104 | 4.113 | OH3.693 |  |  |  |
| Ala^7^ | - | 4.189 | 1.563 |  |  |  |  |
| Glu^8^ | 8.178 | 4.494 | 2.512 |  |  |  |  |
| Asp^9^ | 7.908 | 4.532 | 2.702 | 2.474 |  |  |  |
| Lys^10^ | 7.602 | 4.214 | 2.232 | - | 1.813 | 3.083 | 2.080 |
| Arg^11^ | 7.913 | 4.267 | 1.686 | 1.482 | 3.083 |  |  |
| Arg^12^ | 7.819 | 4.392 | 1.653, 1.529 | 1.330 | 2.969, 2.664 |  |  |
| Ala^13^ | 7.875 | 4.240 | 1.292 |  |  |  |  |
| Tyr^14^ | - | 4.126 | 3.707, 3.363 | 6.947 | 6.794 |  |  |
| Asn^15^ | - | 4.392 | 2.664 |  |  |  |  |

**Supplementary Table 5. Cryo-EM data collection and refinement statistics.**

|  | latT-CoA | latT-AcCoA |
| --- | --- | --- |
| Data collection and processing |  |  |
| Magnification | 105,000 | 105,000 |
| Voltage (kV) | 300 | 300 |
| Electron exposure (e^–^/Å^2^) | 60 | 60 |
| Defocus range (μm) | -2.5 to -0.8 | -2.5 to -0.8 |
| Pixel size (Å) | 0.83 | 0.83 |
| Total micrographs (no.) | 14724 | 9060 |
| Reconstruction |  |  |
| Software | cryoSPARC | cryoSPARC |
| Final particle images (no.) | ~1,440,000 | ~4,350,000 |
| Symmetry imposed | D7 | D7 |
| Map resolution (Å) | 2.2 | 2.0 |
| FSC threshold | 0.143 | 0.143 |
| Map sharpening B factor (Å^2^) | -99.7 | -85.3 |
| Model Building and validation |  |  |
| MolProbity score | 1.05 | 1.12 |
| Clash score | 2.61 | 3.34 |
| Rotamer outliers (%) | 0.00 | 0.00 |
| R.M.S.D. of bond lengths (Å) | 0.003 | 0.002 |
| R.M.S.D. of angles (∘) | 0.703 | 0.454 |
| Favored (%) | 100.00 | 100.00 |
| Allowed (%) | 0.00 | 0.00 |
| Outliers (%) | 0.00 | 0.00 |

**Supplementary Table 6. Turnover ratios of IatT and its variants using 1 and 2 as substrates.**

| **1** as substrate | | | | **2** as substrate | | |
| --- | --- | --- | --- | --- | --- | --- |
| IatT  variants | **1** | **2** | **3** | IatT  variants | **2** | **3** |
| IatT | 0.00%±0.00% | 82.94%±4.42% | 17.06%±4.42% | IatT | 26.80%±3.97% | 73.20%±3.97% |
| A136T | 22.13%±2.64% | 75.25%±1.87% | 2.63%±0.83% | A136T | 28.74%±3.49% | 71.26%±3.49% |
| A136L | 22.59%±6.17% | 70.88%±2.55% | 6.53%±3.63% | A136L | 34.32%±3.31% | 65.68%±3.31% |
| A136E | 18.57%±3.34% | 70.52%±1.49 | 10.91%±1.86% | A136E | 21.29%±4.81% | 78.71%±4.81% |
| Y42A | 9.62%±5.26% | 78.49%±2.89% | 11.89%±3.64% | Y42A | 91.66%±0.62% | 8.347%±0.62% |
| D48A | 0.00%±0.00% | 82.96%±1.62% | 17.04%±1.62% | D48A | 67.25%±3.60% | 32.75%±3.60% |
| N52A | 1.66%±0.69% | 78.91%±3.45% | 19.42%±1.05% | N52A | 60.28%±3.03% | 39.72%±3.03% |
| R53A | 3.98%±0.28% | 62.92%±0.80% | 33.10%±1.03% | R53A | 70.66%±2.76% | 29.34%±2.76% |
| R53W | 0.83%±0.62% | 85.77%±1.88% | 13.40%±1.72% | R53W | 89.45%±1.80% | 10.55%±1.80% |
| H59A | 0.94%±1.78% | 73.94%±4.54% | 25.12%±1.05% | H59A | 38.02%±1.70% | 61.98%±1.70% |
| Q60A | 6.20%±0.49% | 72.49%±5.43% | 26.57%±5.77% | Q60A | 64.87%±3.38% | 35.13%±3.38% |
| E98A | 6.20%±3.62% | 87.56%±1.74 | 6.25%±2.25% | E98A | 91.49%±0.74% | 8.51%±0.74% |
| V100W | 0.10%±0.10% | 99.83%±0.16% | 0.07%±0.07% | V100W | 78.17%±6.21% | 21.83%±6.21% |
| H134A | 8.78%±1.47% | 75.11%±2.59 | 16.11%±1.13% | H134A | 63.39%±0.87% | 36.61%±0.87% |
| H134Y | 5.15%±0.83 | 94.64%±0.61% | 0.21%±0.29% | H134Y | 79.35%±3.65% | 20.65%±3.65% |
| R159A | 1.59%±1.11% | 86.06%±3.07% | 12.35%±1.97% | R159A | 94.93%±0.59% | 5.07%±0.59% |
| R44A | 63.04%±3.71% | 26.07%±0.99% | 10.89%±4.66% | R44A | 87.47%±2.19% | 12.53%±2.19% |
| E55A | 10.67%±0.99% | 57.02%±2.00% | 30.65%±2.03% | E55A | 79.75%±1.30% | 20.25%±1.30% |
| A136Y | 21.30%±0.54% | 75.22%±0.90% | 3.49%±0.92% | A136Y | 58.08%±4.75% | 41.921%±4.75% |
| G163F | 31.38%±7.77% | 57.97%±4.08% | 10.65%±3.72% | G163F | 86.61%±2.42% | 13.39%±2.42% |
| E172A | 19.27%±1.29% | 63.31%±1.56% | 17.42%±0.34% | E172A | 89.04%±2.68% | 10.96%±2.68% |
| Y42W | 31.44%±5.08% | 56.56%±5.22% | 12.00%±5.60% | Y42W | 10.74%±0.60% | 89.26%±0.60% |
| A136H | 23.06%±6.39% | 35.35%±4.56% | 41.59%±2.60% | A136H | 10.41%±1.95% | 89.59%±1.95% |
| A136K | 11.98%±3.22% | 79.54%±1.35% | 8.48%±3.06% | A136K | 1.92%±1.92% | 89.73%±1.92% |
| E98W | 3.40%±4.61% | 40.92%±3.76% | 55.68%±2.60% | E98W | 5.37%±0.49% | 94.63%±0.49% |
| E172W | 3.77%±3.02% | 23.99%±4.84% | 72.24%±4.84% | E172W | 7.50%±0.57% | 92.50%±0.57% |

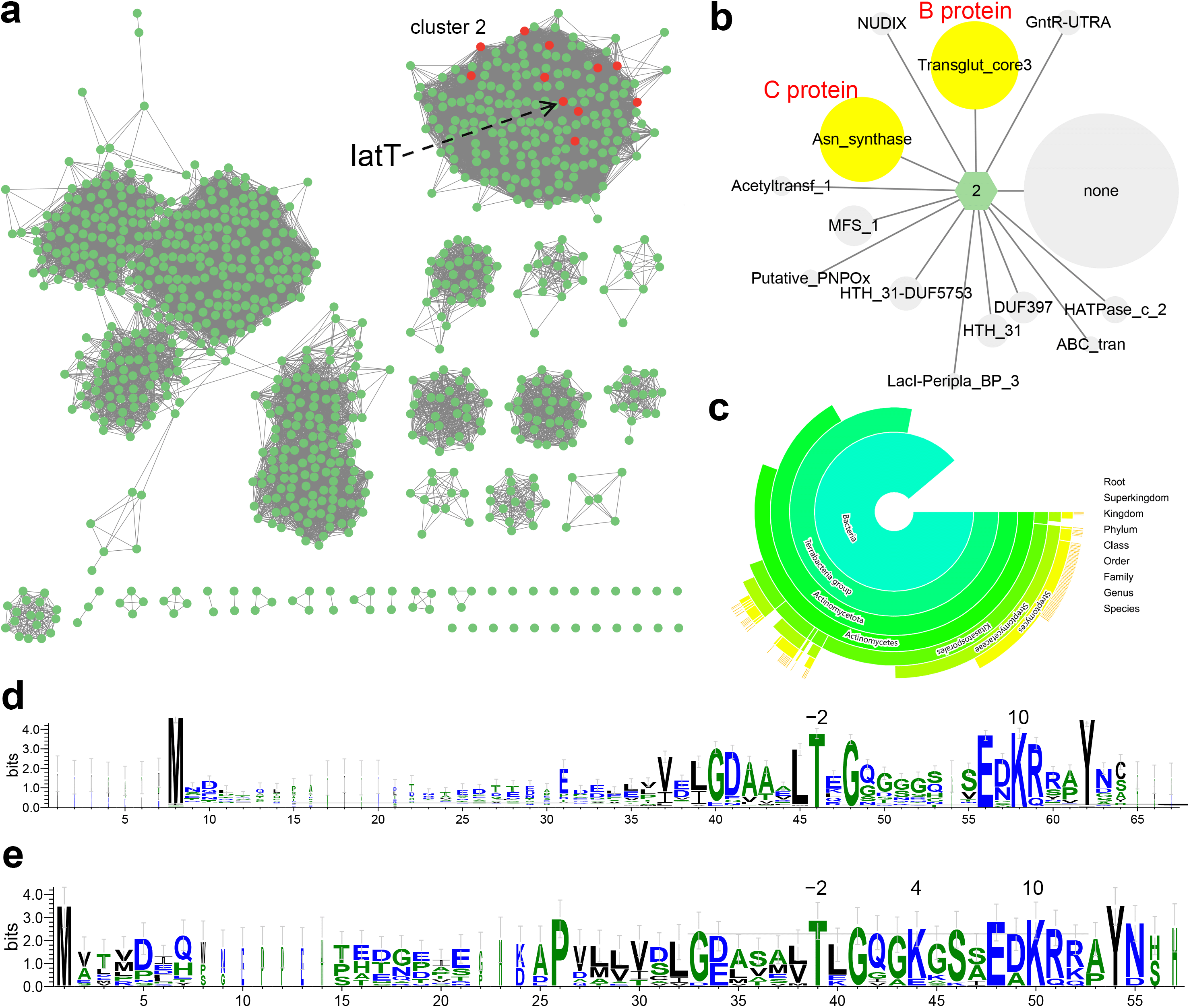

**Supplementary Fig. 1. Bioinformatic analysis of GNAT-coding lasso peptide BGCs.** **a**, Complete SSN of IatT with class II GNATs shown in red. **b**, GNN of the IatT-containing SSN cluster 2, showing the cooccurrence with lasso peptide biosynthetic proteins B and C. **c**, The taxonomy distribution for the lasso peptide GNATs. **d**,**e**, Sequence logo plots of the PPs from class I (**d**) and II (**e**) BGCs show the conservation of the Lys residues.

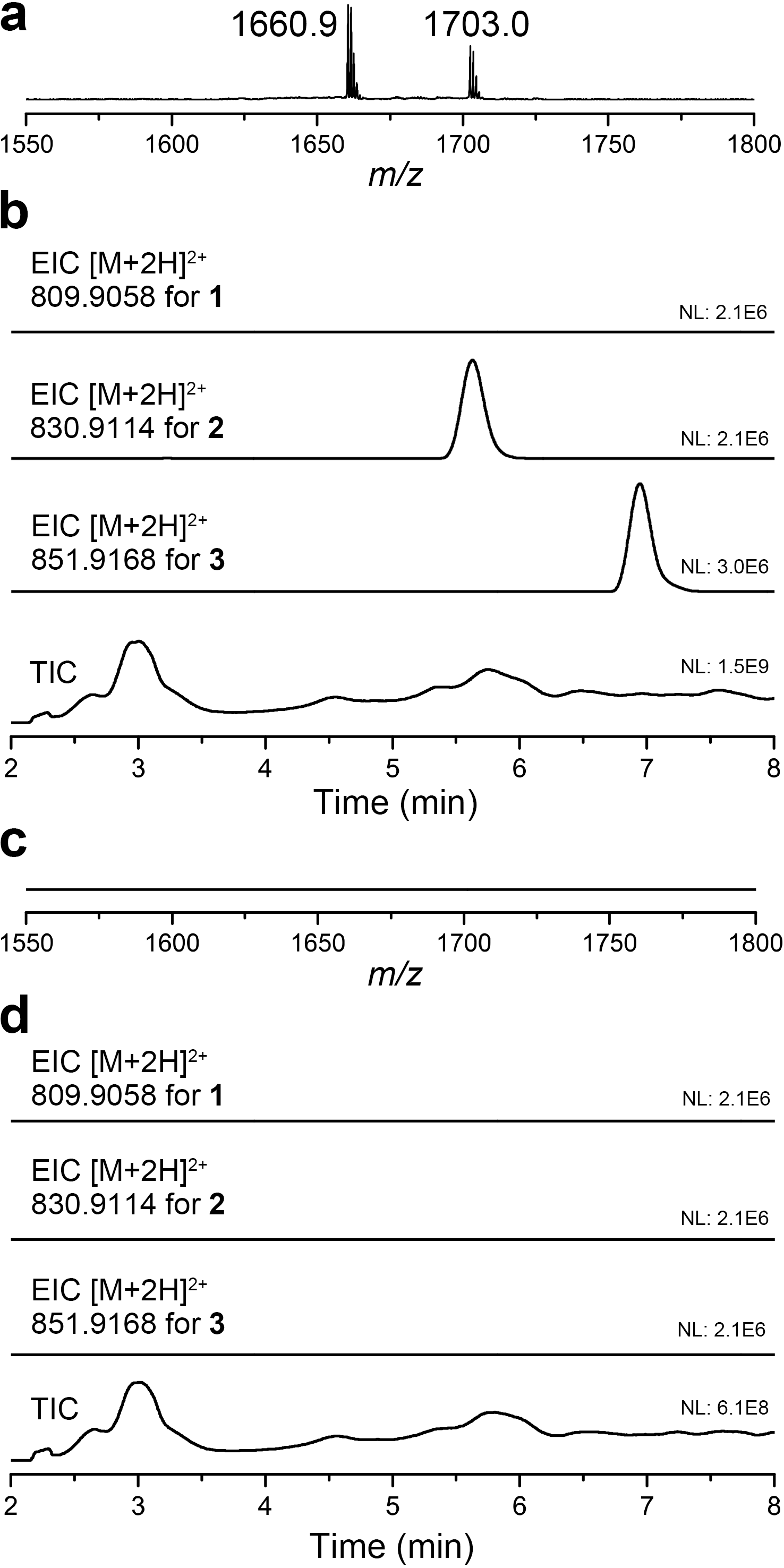

**Supplementary Fig. 2. MS analysis of heterologous expression products of *iat* BGC.** **a**, MALDI-TOF MS analysis of M1154-*iat* products. **b**, LC-HRMS analysis of M1154-*iat* products. **c**, MALDI-TOF MS analysis of M1154-*iat*-△*iatT* products. **d**, LC-HRMS analysis of M1154-*iat-*△*iatT* products.

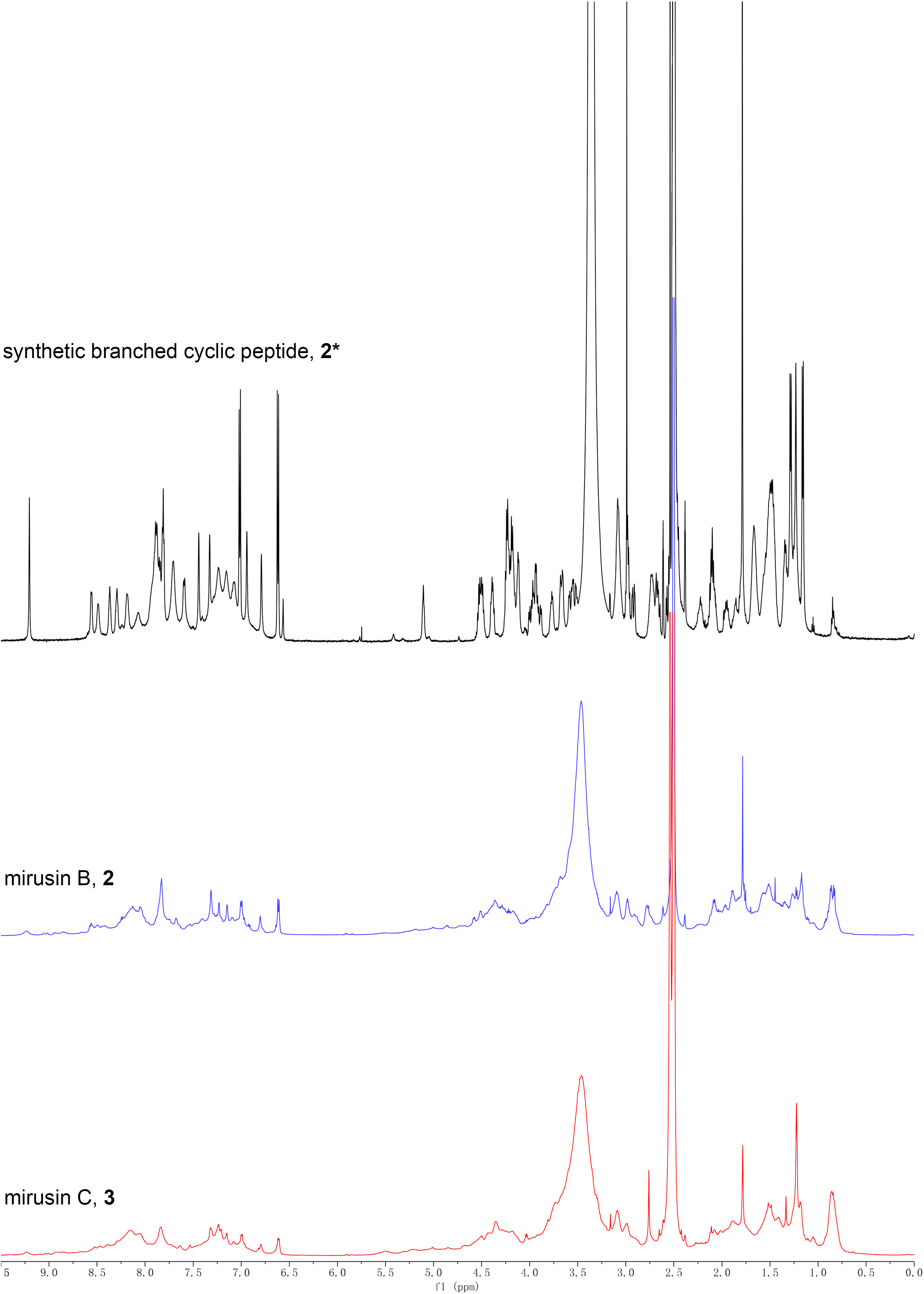

**Supplementary Fig. 3.** **Comparison of ^1^H NMR spectra of 2*, 2, and 3.**

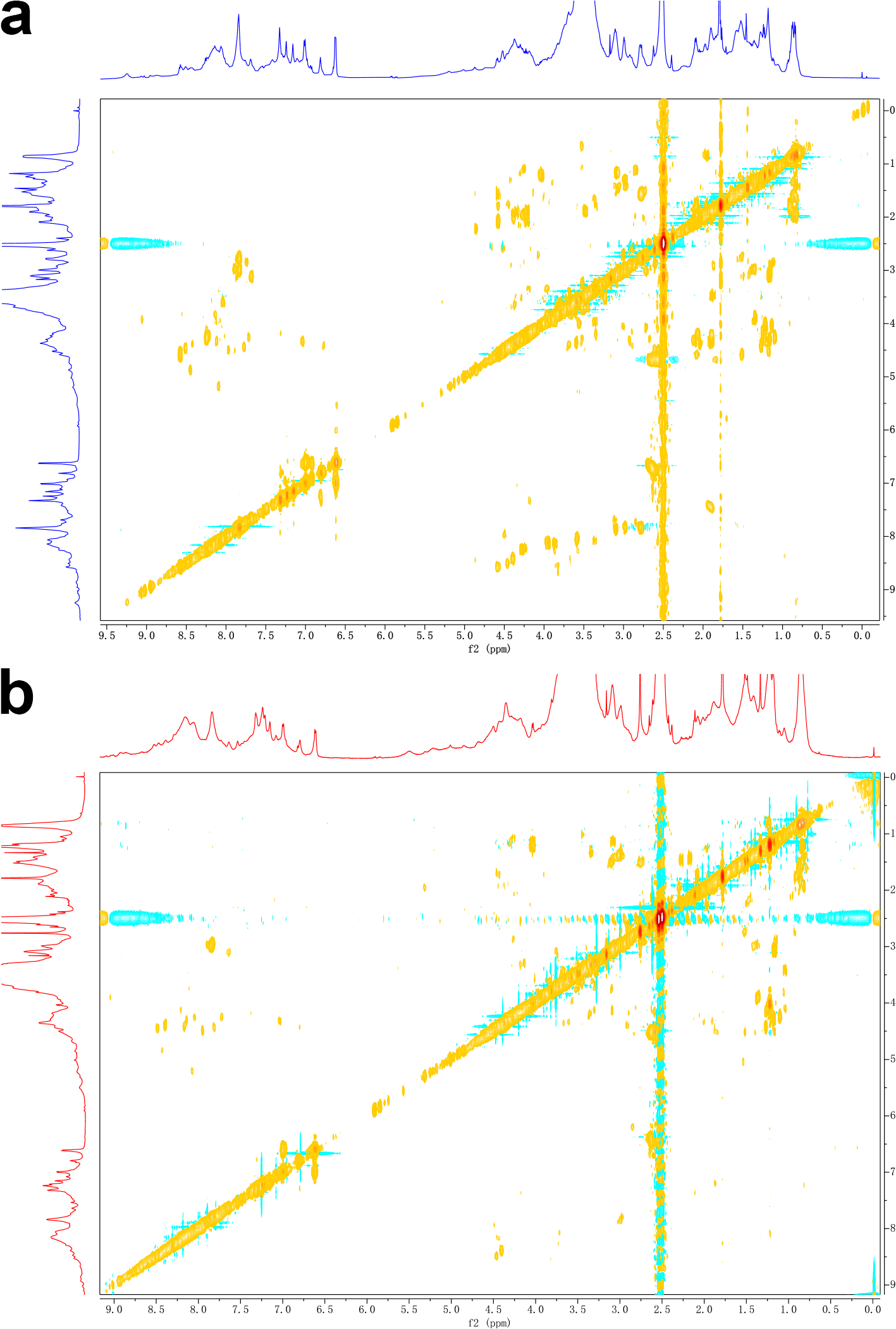

**Supplementary Fig. 4. ^1^H-^1^H COSY NMR spectra of 2 (a) and 3 (b).**

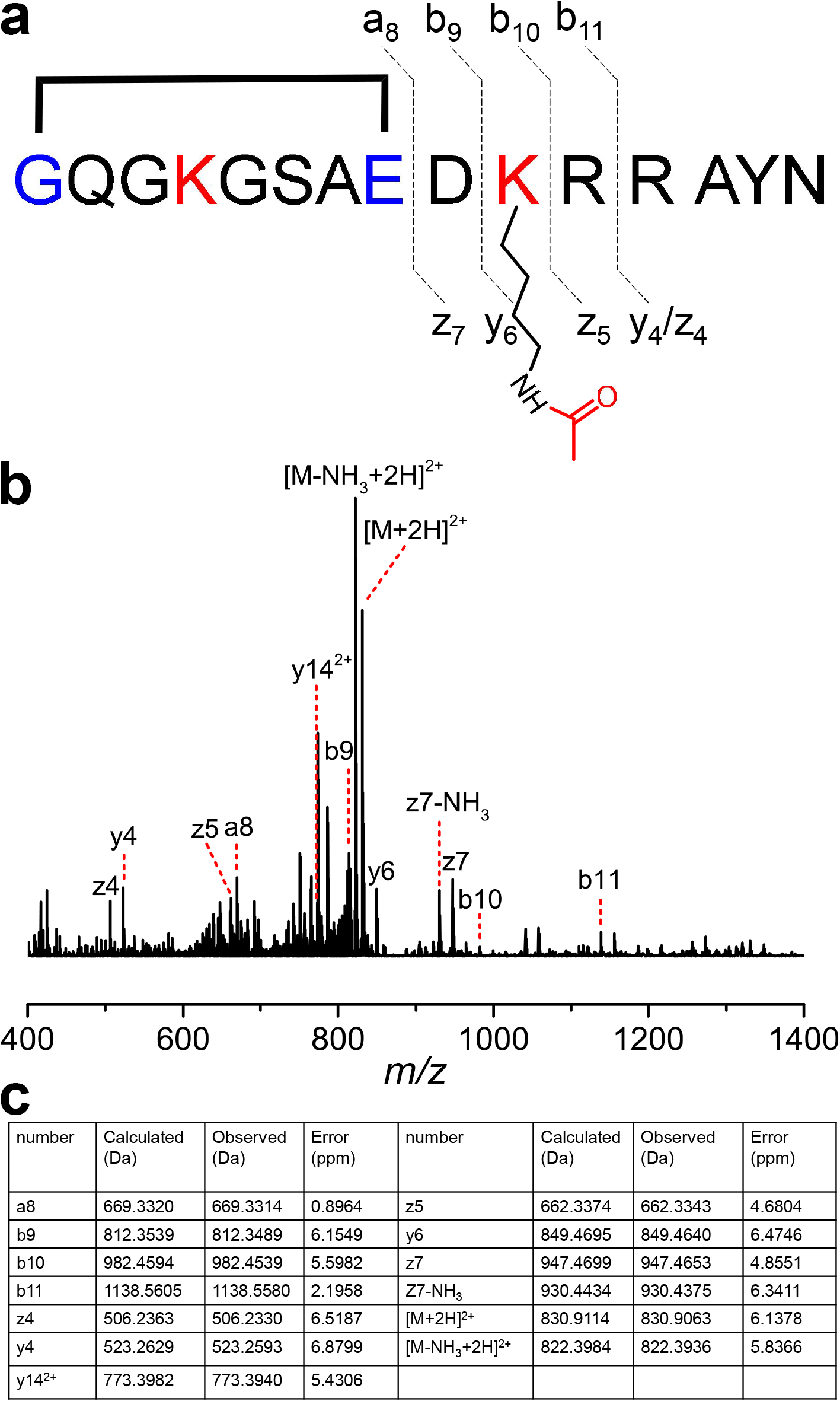

**Supplementary Fig. 5. HRMS/MS analysis of 2*.** **a**, Annotation of fragment ions of **2***. **b**, HRMS/MS spectrum of **2***. **c**, Detailed information of identified fragment ions.

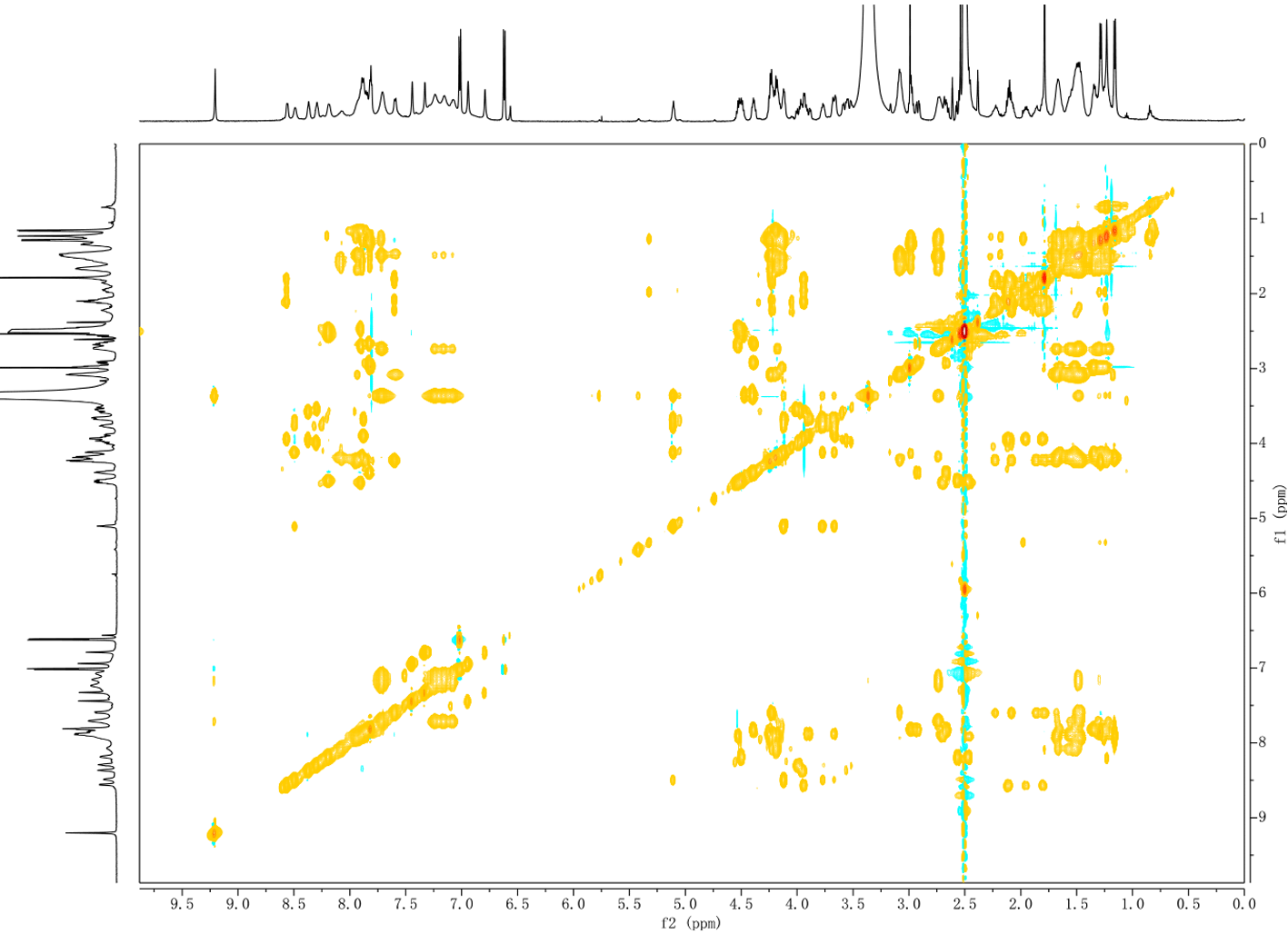

**Supplementary Fig. 6. ^1^H-^1^H COSY NMR spectra of 2*.**

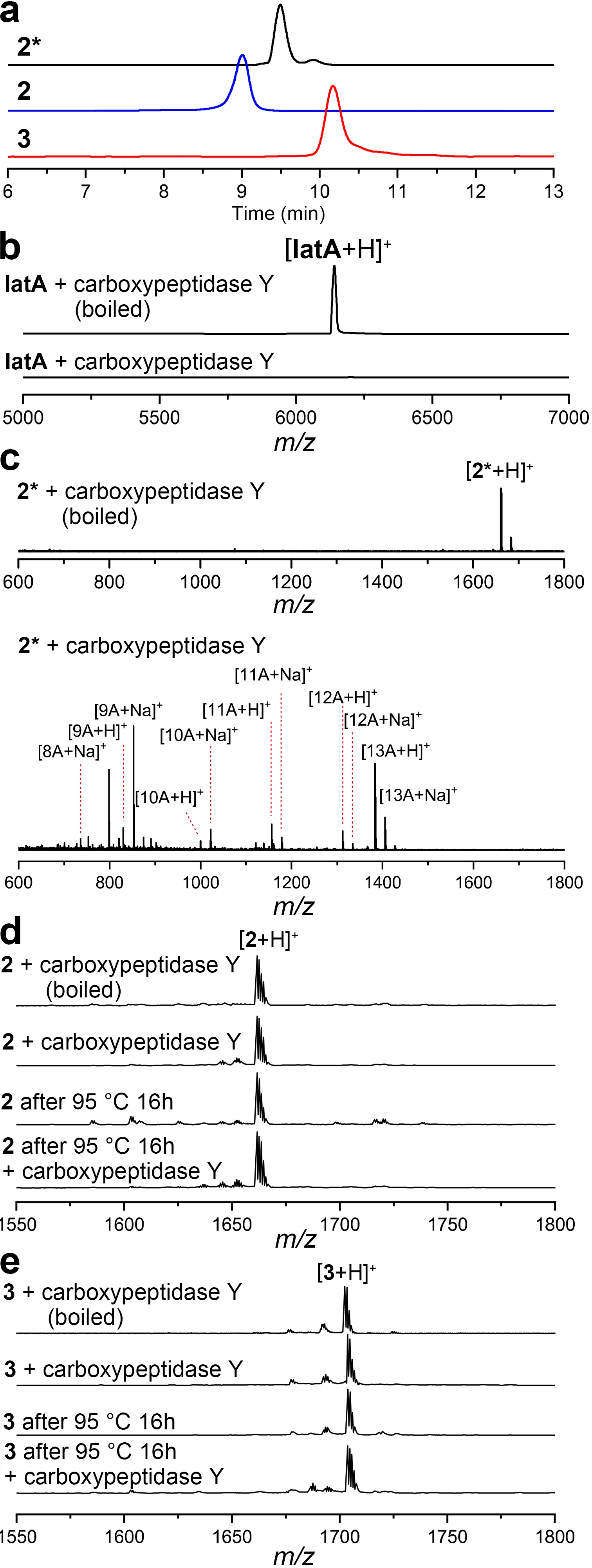

**Supplementary Fig. 7. Confirmation of the presence of lasso structure in 2 and 3.** **a,** comparison of the elution profiles of **2***, **2**, and **3**. MALDI-TOF MS analysis of IatA (**b**), **2*** (**c**), **2** (**d**), and **3** (**e**) after carboxypeptidase Y digestion. The mass signals of hydrolysis products of **2*** were indicated by the leftover residue numbers.

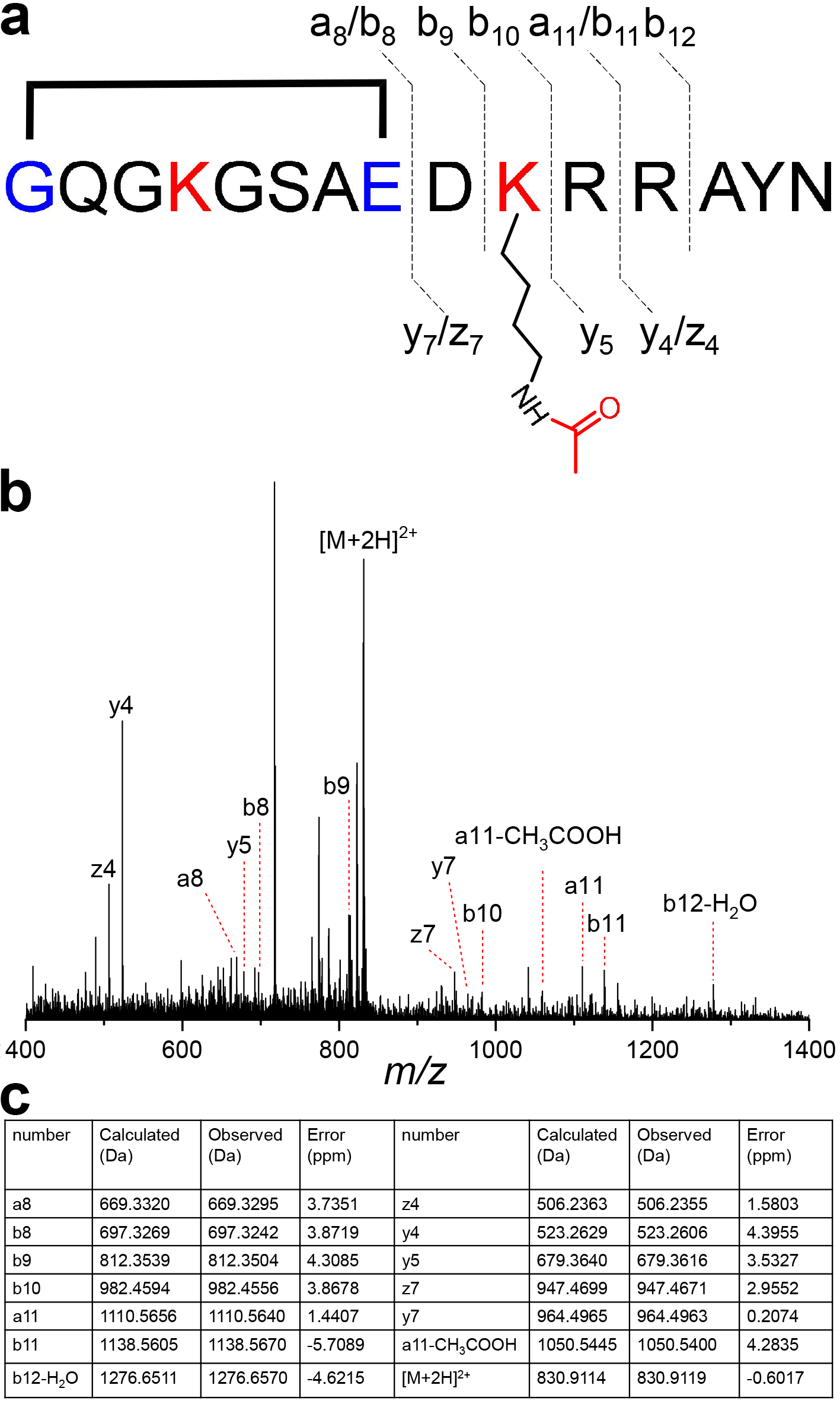

**Supplementary Fig. 8.** **HRMS/MS analysis of 2.** **a**, Annotation of fragment ions of **2**. **b**, HRMS/MS spectrum of **2**. **c**, Detailed information of identified fragment ions. Structure is shown in simplified model with unthreaded conformation for clarity.

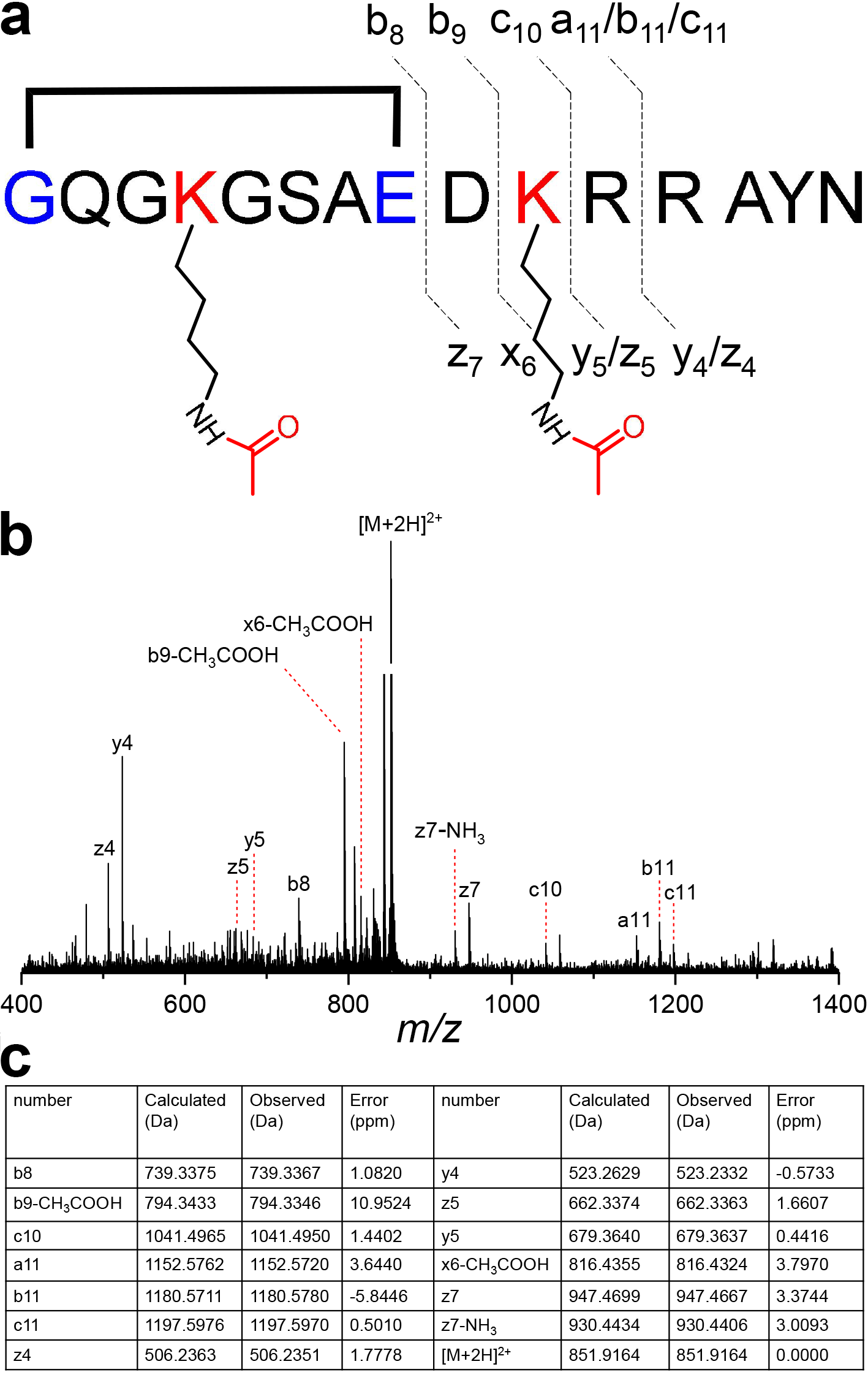

**Supplementary Fig. 9. HRMS/MS analysis of 3.** **a**, Annotation of fragment ions of **3**. **b**, HRMS/MS spectrum of **3**. **c**, Detailed information of identified fragment ions. Structure is shown in simplified model with unthreaded conformation for clarity.

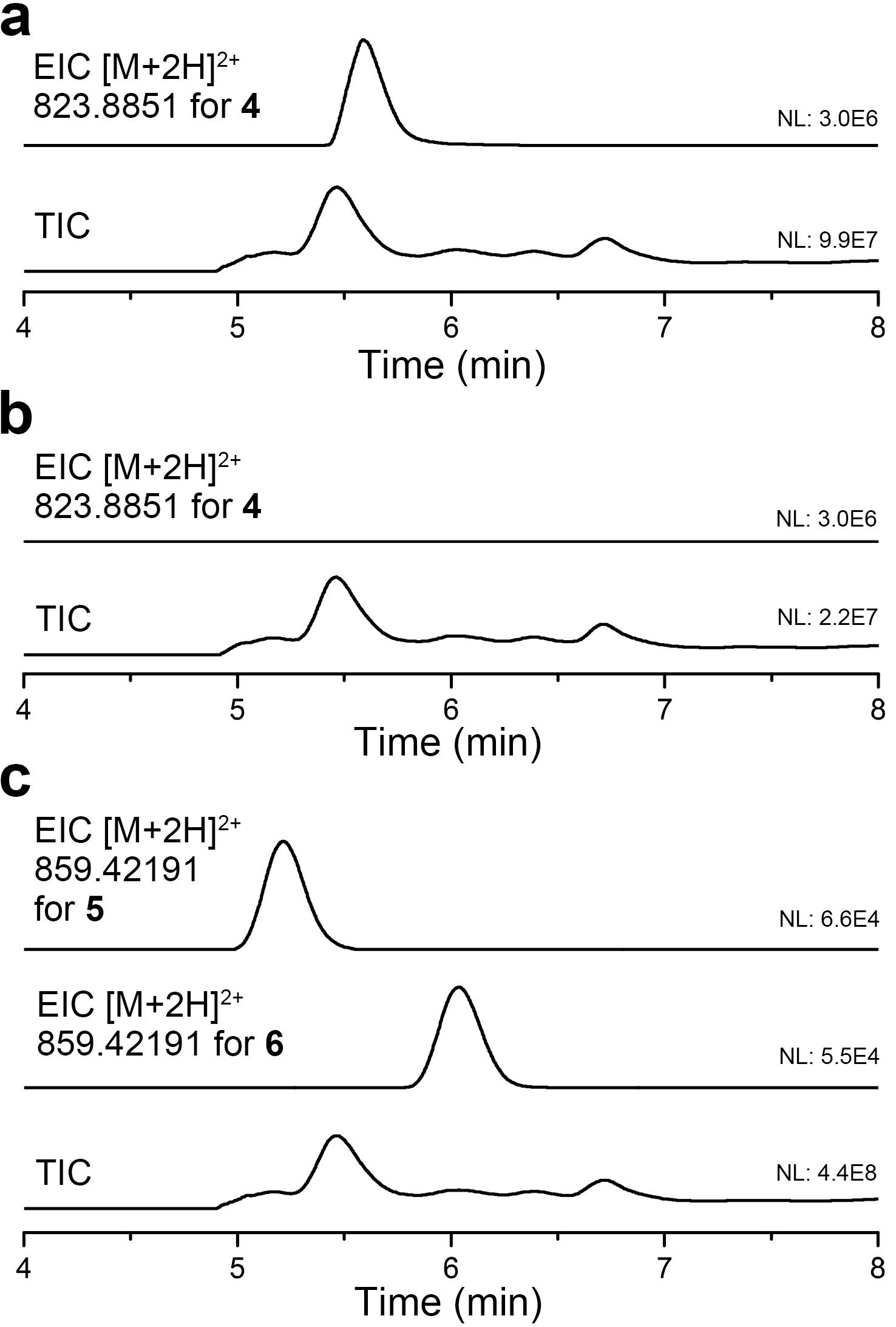

**Supplementary Fig. 10. LC-HRMS analysis of heterologous expression products of *ven* BGC containing *venABCT* (a), *venABC* (b), and *venA(G4K)BCT* (c) genes.**

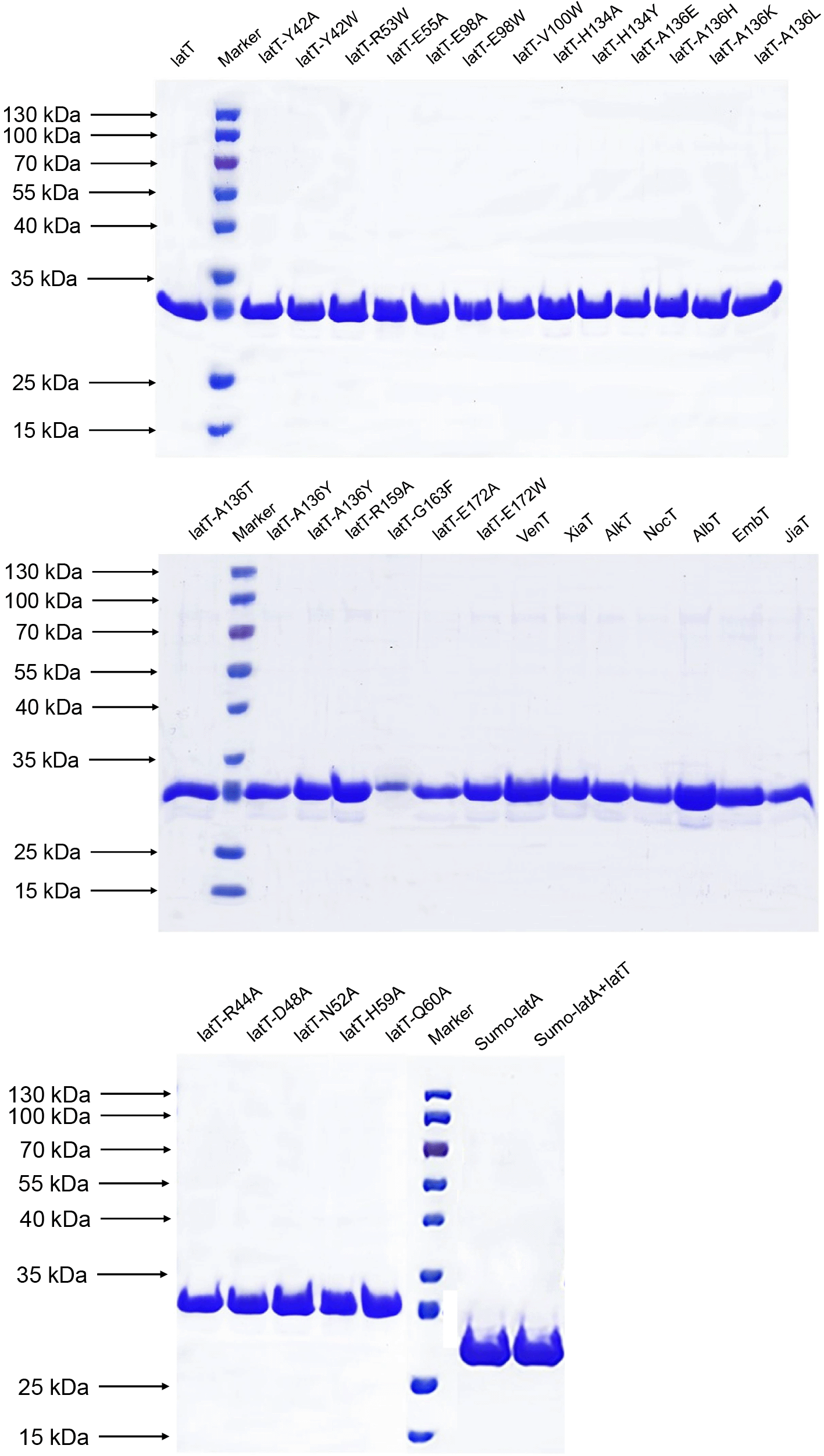

**Supplementary Fig. 11. SDS-PAGE analysis of purified proteins.**

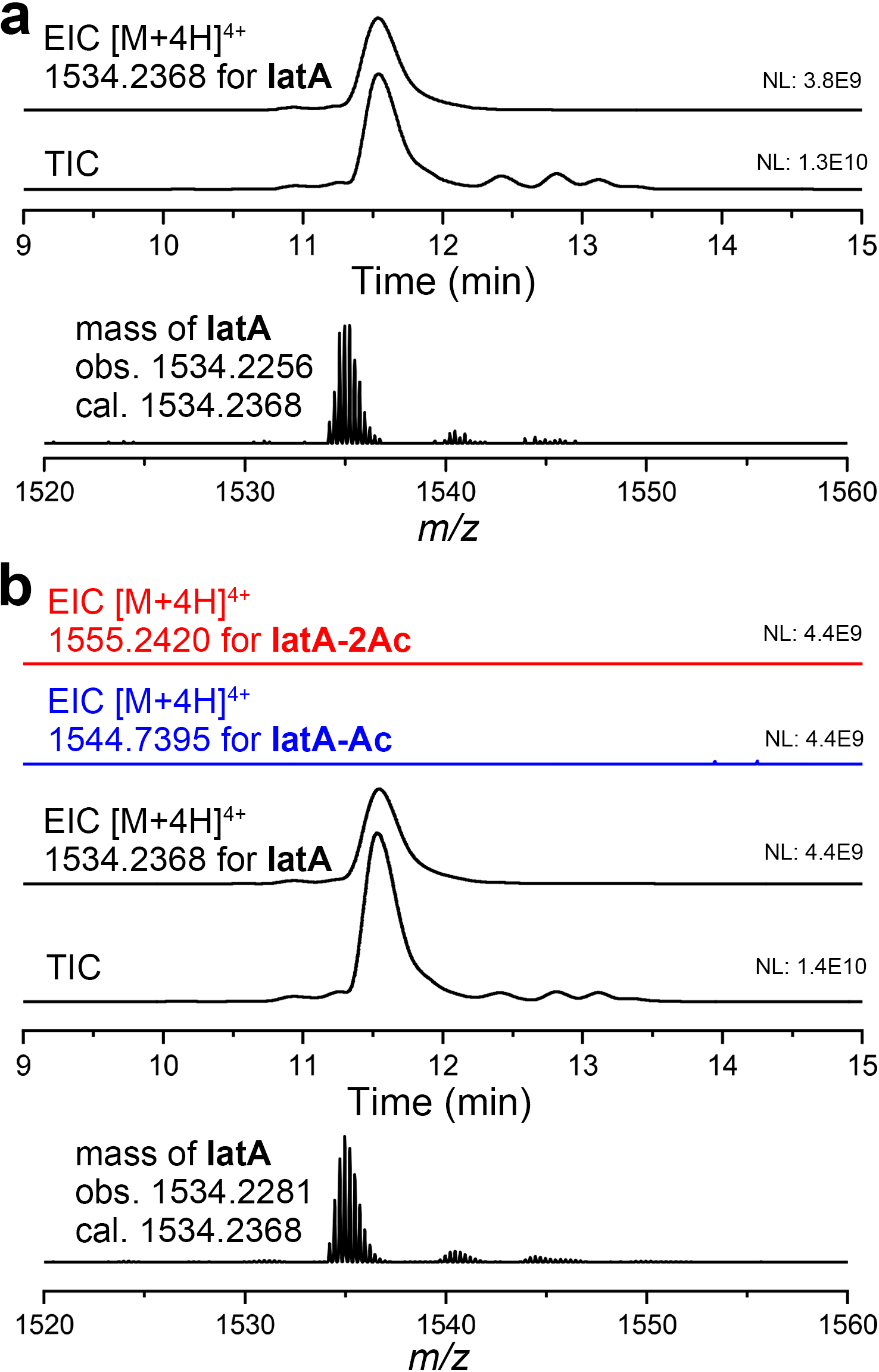

**Supplementary Fig. 12.** **LC-HRMS analysis of the TEV-digested SUMO-IatA. a**, His_6_-SUMO-IatA expressed in *E. coli* BL21(DE3). **b**, His_6_-SUMO-IatA co-expressed with IatT in *E. coli* BL21(DE3).

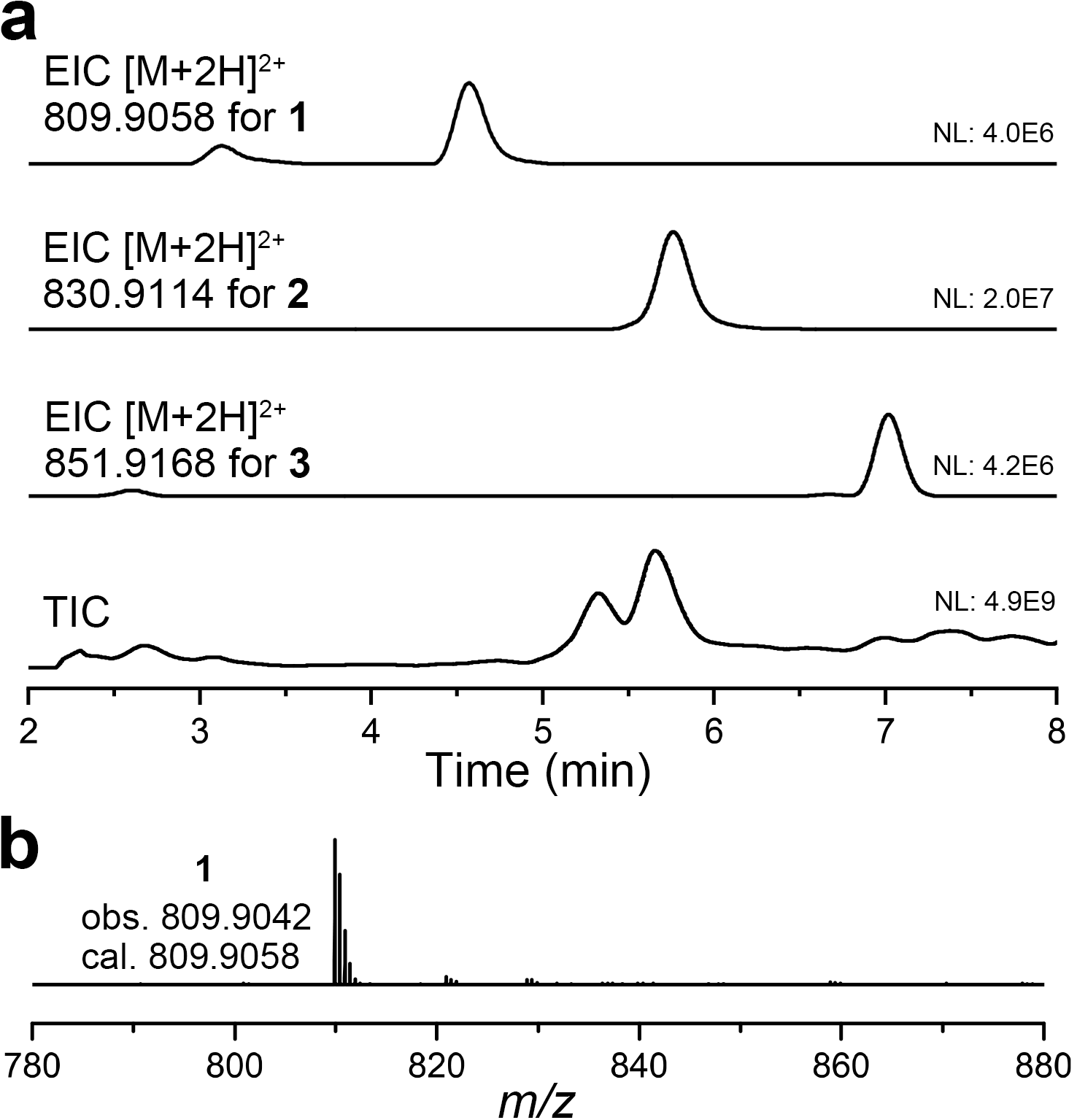

**Supplementary Fig. 13. LC-HRMS analysis of heterologous expression products of *iat*-HM1 BGC.** **a**, LC-HRMS analysis of M1154-*iat*-HM1 products. **b**, HRMS spectrum of compound **1**.

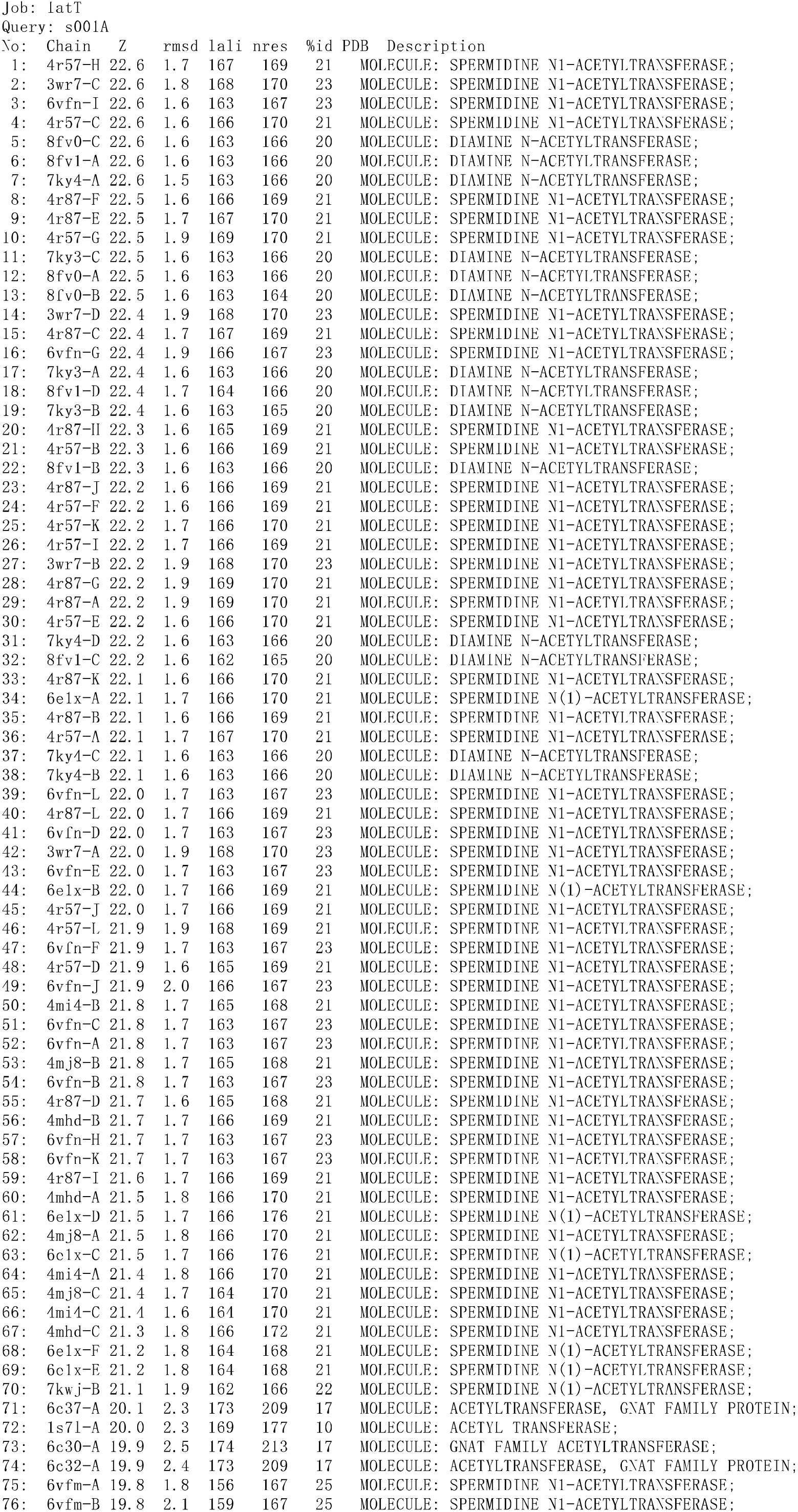

**Supplementary Fig. 14.** **DALI search results of IatT-AcCoA cryo-EM structure.**

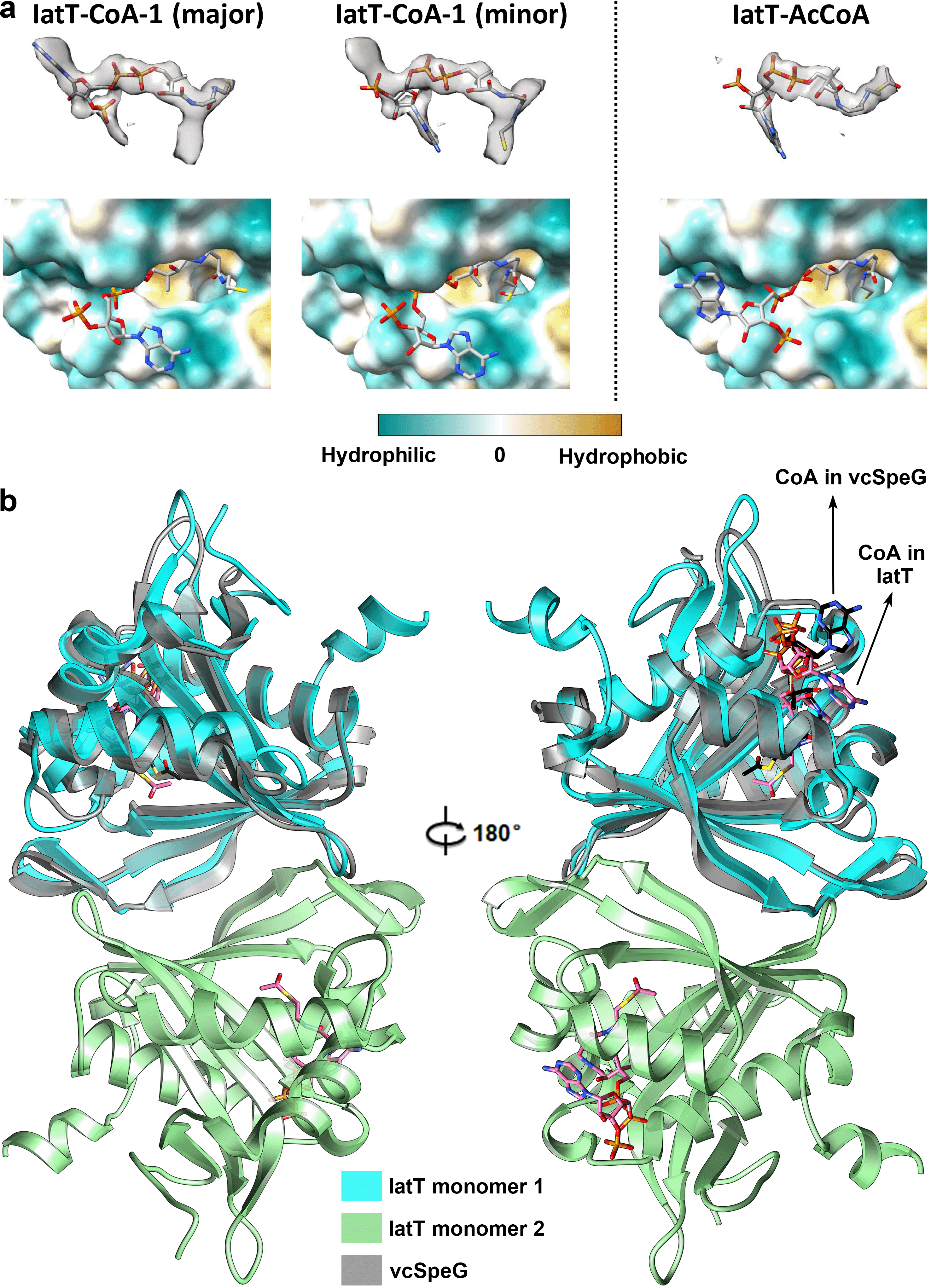

**Supplementary Fig. 15.** **Binding modes of CoA and AcoA with IatT.** **a**, Density maps of CoA and AcCoA in the binding pocket of IatT. Modelling of CoA into the density map indicates the presence of two conformations. CoA and AcCoA are shown in sticks, and IatT is shown as surface model. **b**, Comparison of the binding poses of AcCoA in the complex structures of IatT and vcSpeG.
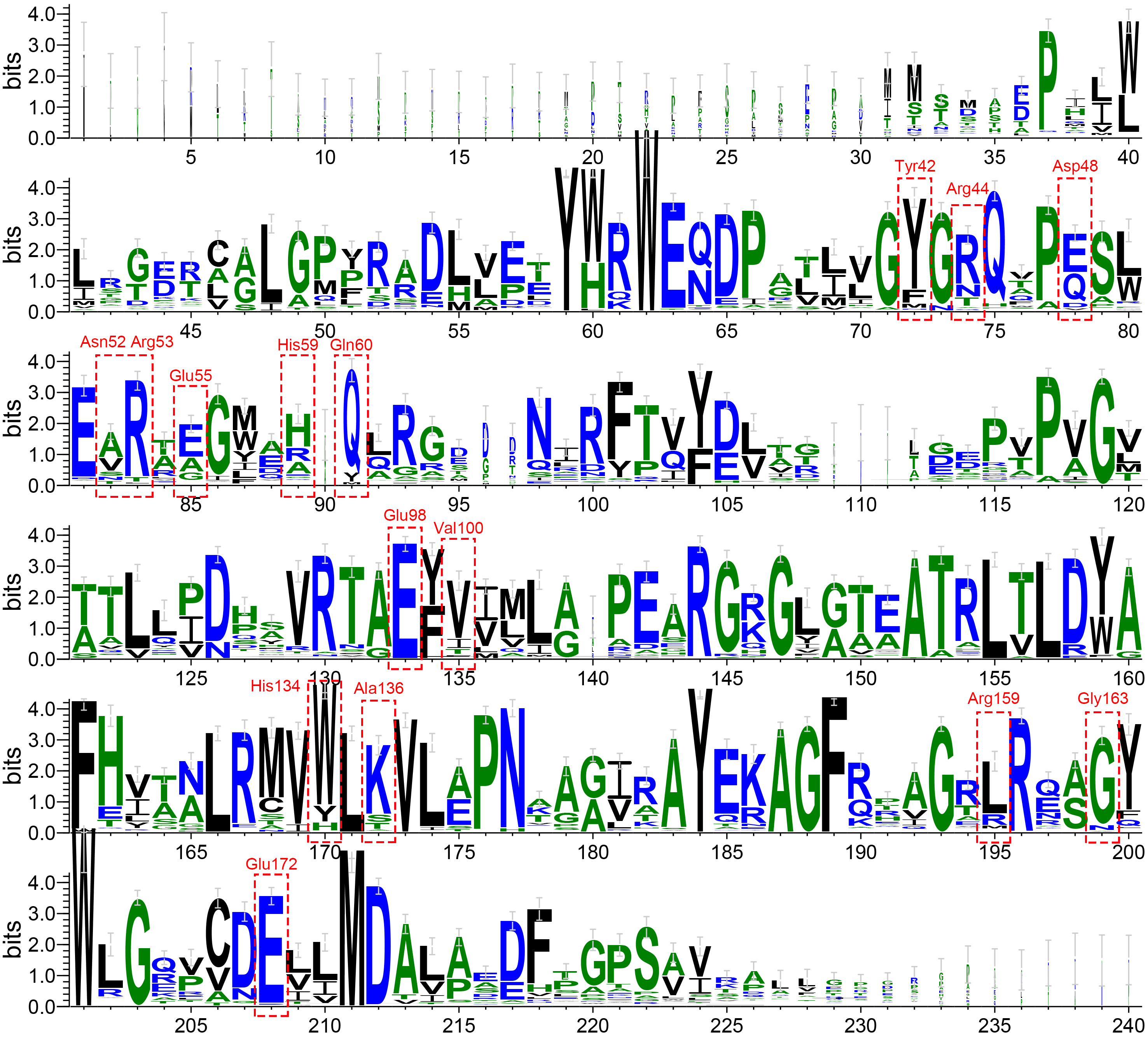

**Supplementary Fig. 16. Sequence logo plot of the 112 IatT-like GNATs.** The 15 residues surrounding the potential lasso peptide binding pocket are highlighted with red dashed rectangles. Residue identities are indicated in red characters, corresponding to the IatT amino acid numbers.

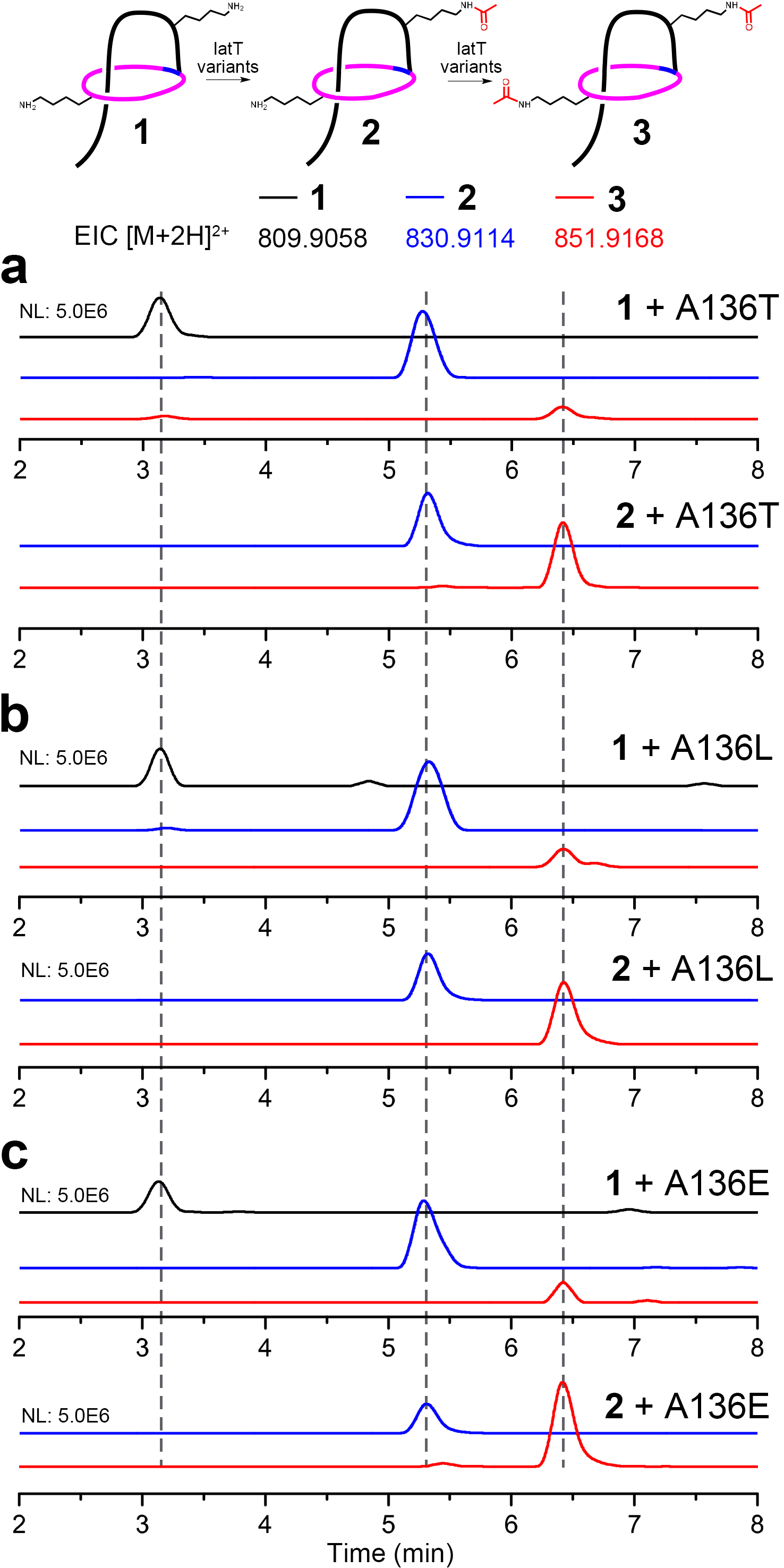

**Supplementary Fig. 17.** **Enzymatic activity assays of IatT-A136T, A136L, and A136E variants in vitro via LC-HRMS analysis.** EICs for assays of IatT-A136T (**a**), A136L (**b**), and A136E (**c**).

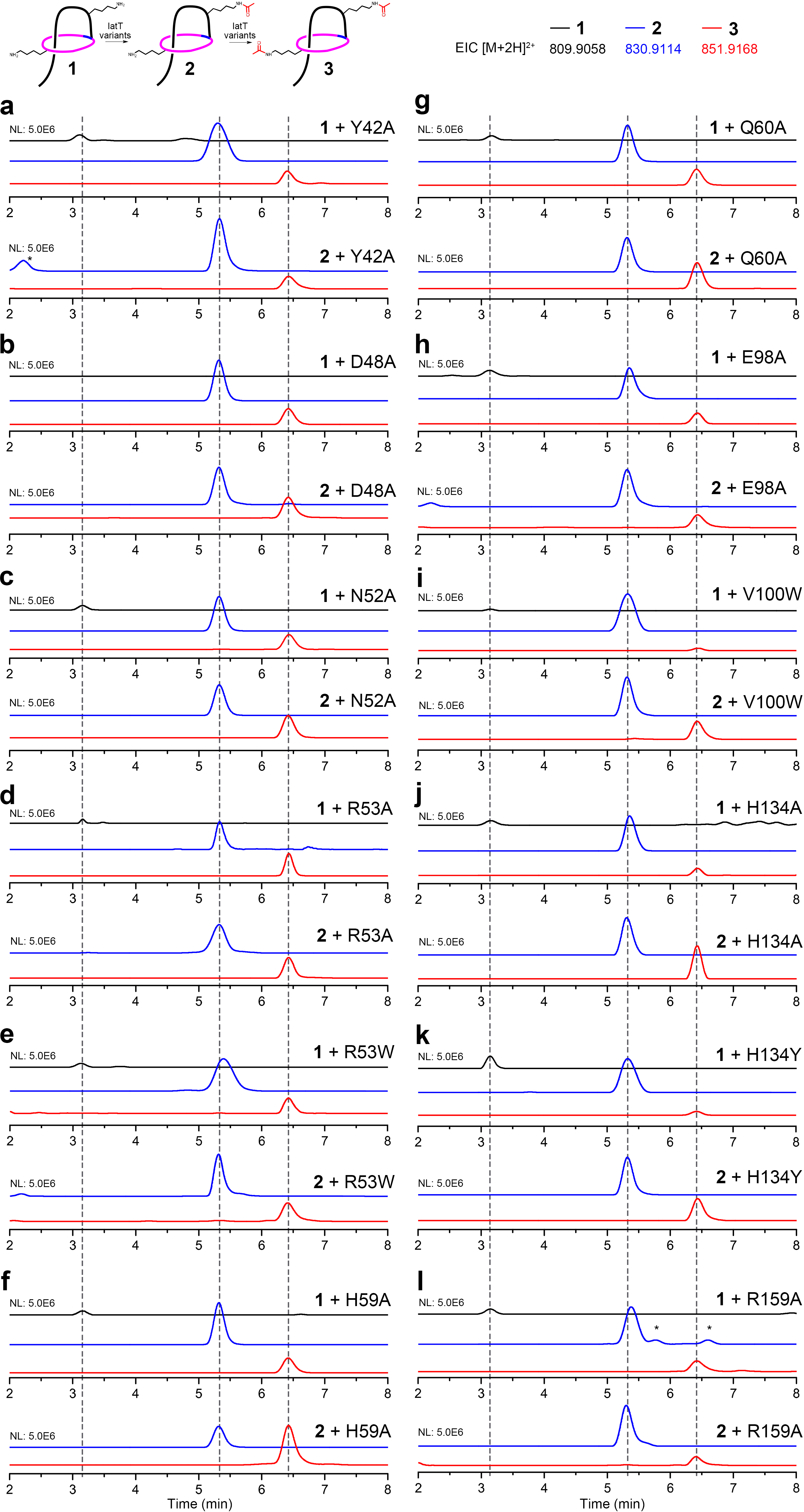

**Supplementary Fig. 18. Enzymatic activity assays of IatT-Y42A, D48A, N52A, R53A, R53W, H59A, Q60A, E98A, V100W, H134A, H134Y, and R159A variants in vitro via LC-HRMS analysis.** EICs for assays of IatT-Y42A (**a**), D48A (**b**), N52A (**c**), R53A (**d**), R53W (**e**), H59A (**f**), Q60A (**g**), E98A (**h**), V100W (**i**), H134A (**j**), H134Y (**k**), and R159A (**l**). * indicates unrelated impurity showing identical mass with [M+H]^+^ instead of [M+H]^2+^ ion.

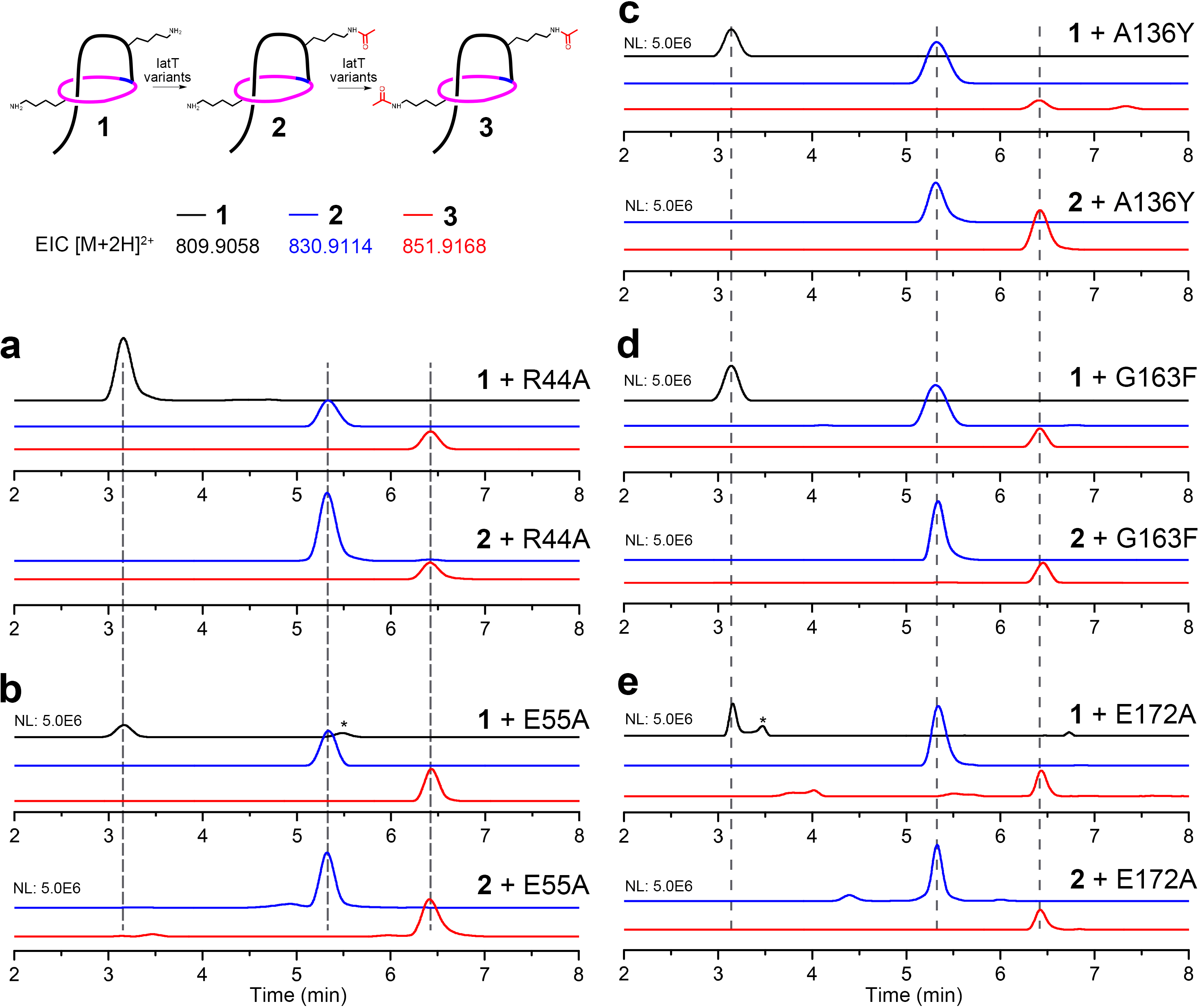

**Supplementary Fig. 19. Enzymatic activity assays of IatT-R44A, E55A, A136Y, G163F, and E172A variants in vitro via LC-HRMS analysis.** EICs for assays of IatT-R44A (**a**), E55A (**b**), A136Y (**c**), G163F (**d**), and E172A (**e**). * indicates unrelated impurity showing identical mass with [M+H]^+^ instead of [M+H]^2+^ ion.

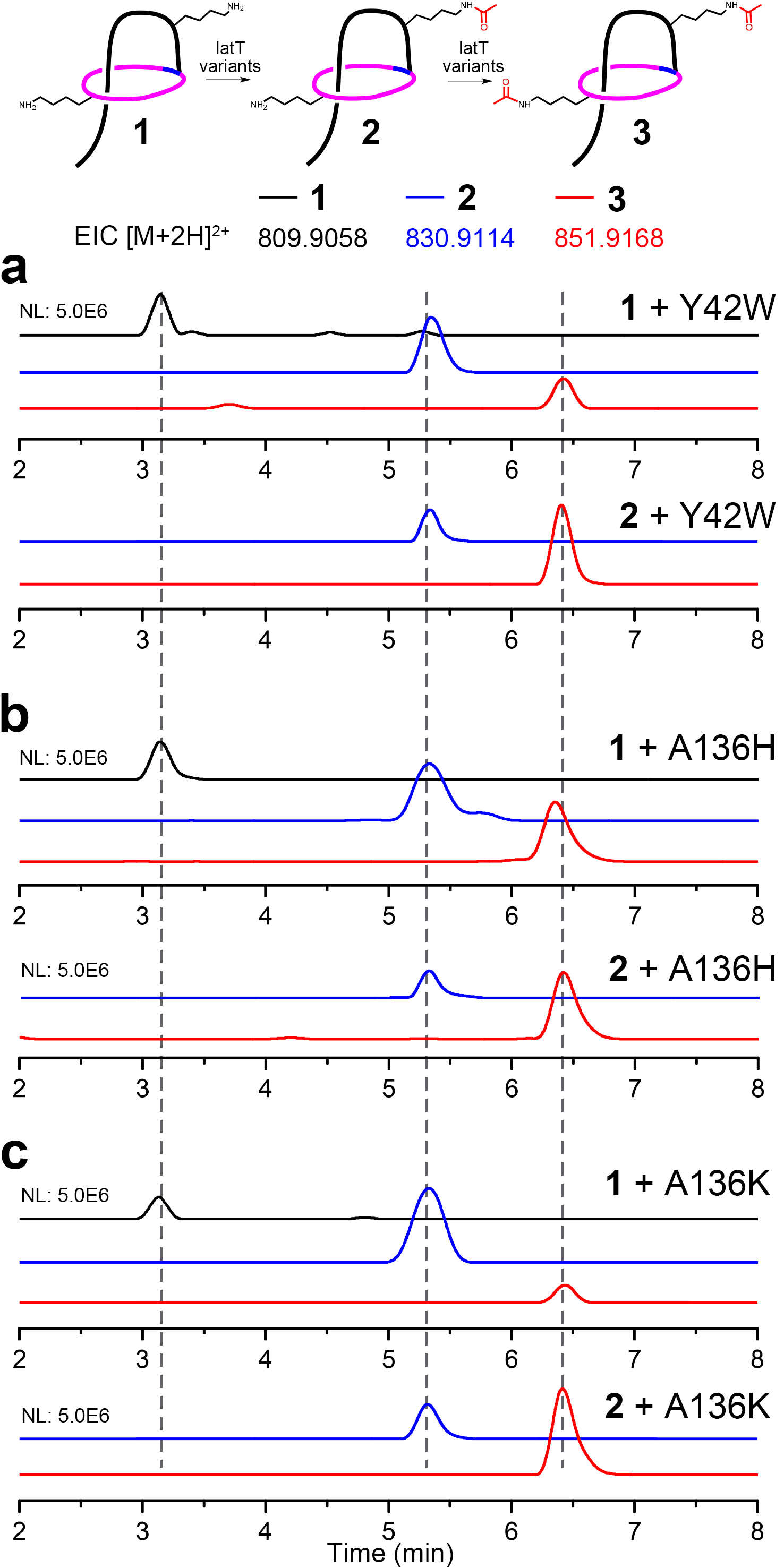

**Supplementary Fig. 20.** **Enzymatic activity assays of IatT-Y42W, A136H, and A136K variants in vitro via LC-HRMS analysis.** EICs for assays of IatT-Y42W (**a**), A136H (**b**), and A136K (**c**).

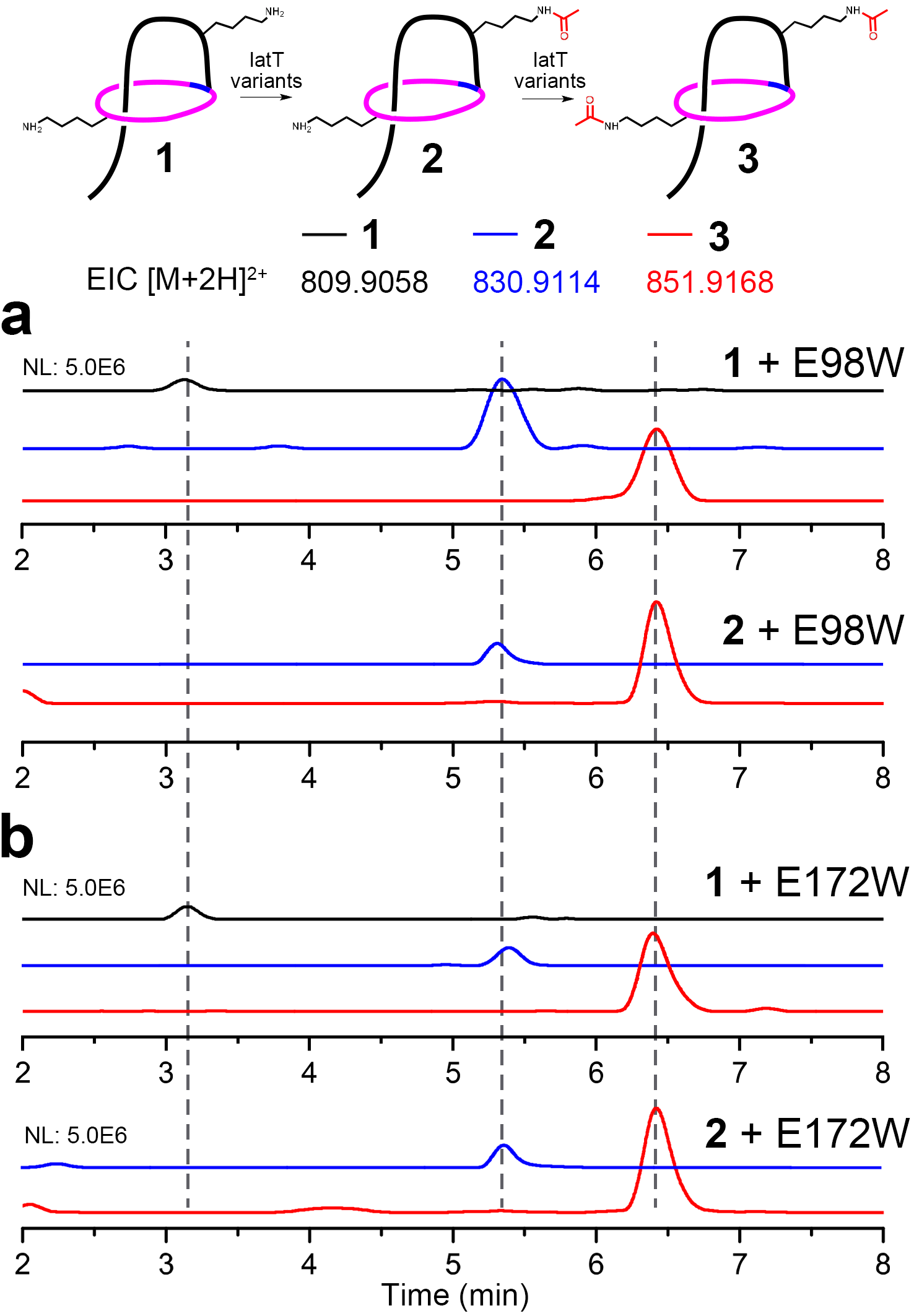

**Supplementary Fig. 21. Enzymatic activity assays of IatT-E98W and E172W variants in vitro via LC-HRMS analysis.** EICs for assays of IatT-E98W (**a**) and E172W (**b**).

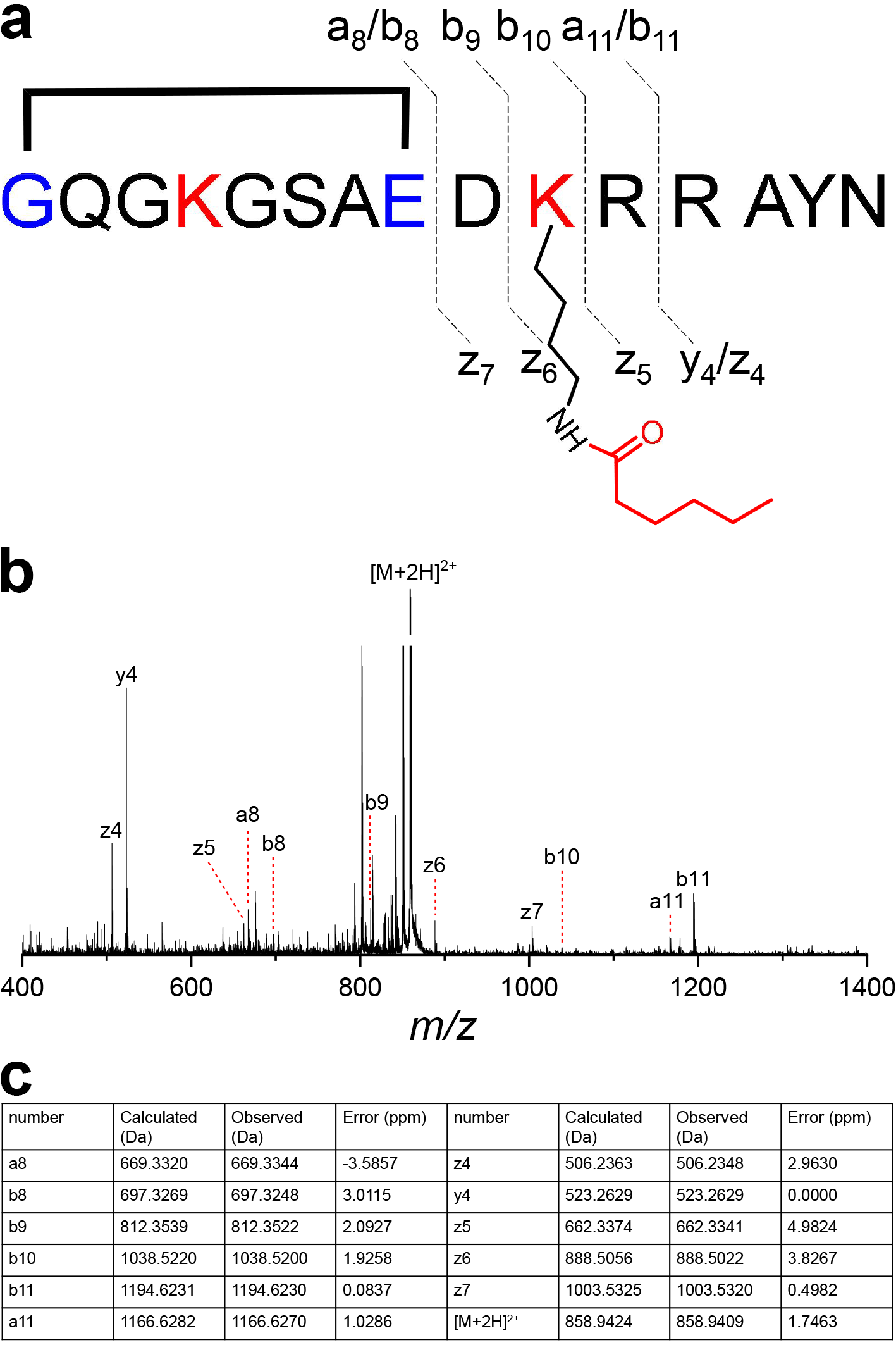

**Supplementary Fig. 22. HRMS/MS analysis of 8.** **a**, Annotation of fragment ions of **8**. **b**, HRMS/MS spectrum of **8**. **c**, Detailed information of identified fragment ions. Structure is shown in simplified model with unthreaded conformation for clarity.

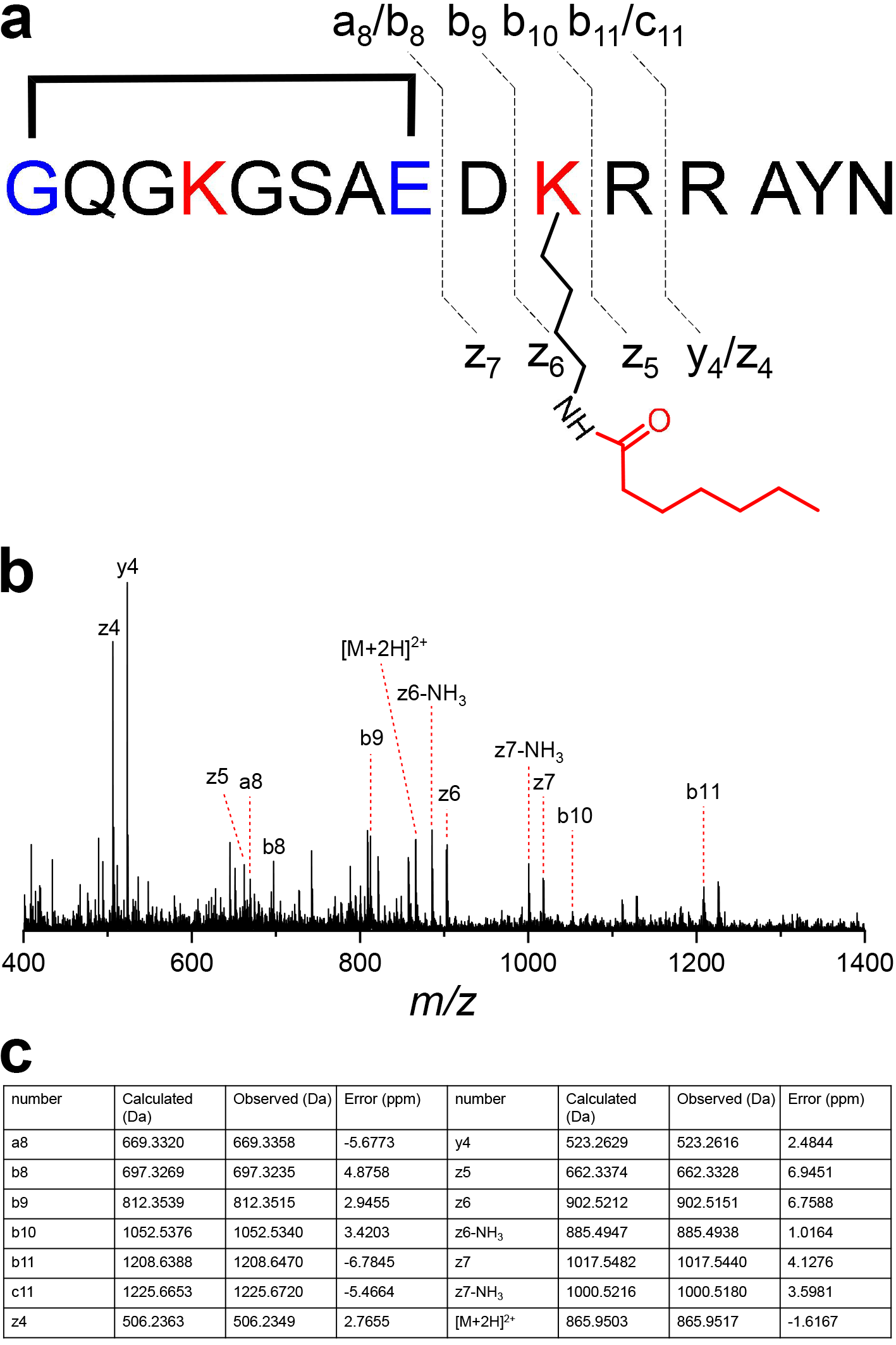

**Supplementary Fig. 23. HRMS/MS analysis of 9.** **a**, Annotation of fragment ions of **9**. **b**, HRMS/MS spectrum of **9**. **c**, Detailed information of identified fragment ions. Structure is shown in simplified model with unthreaded conformation for clarity.

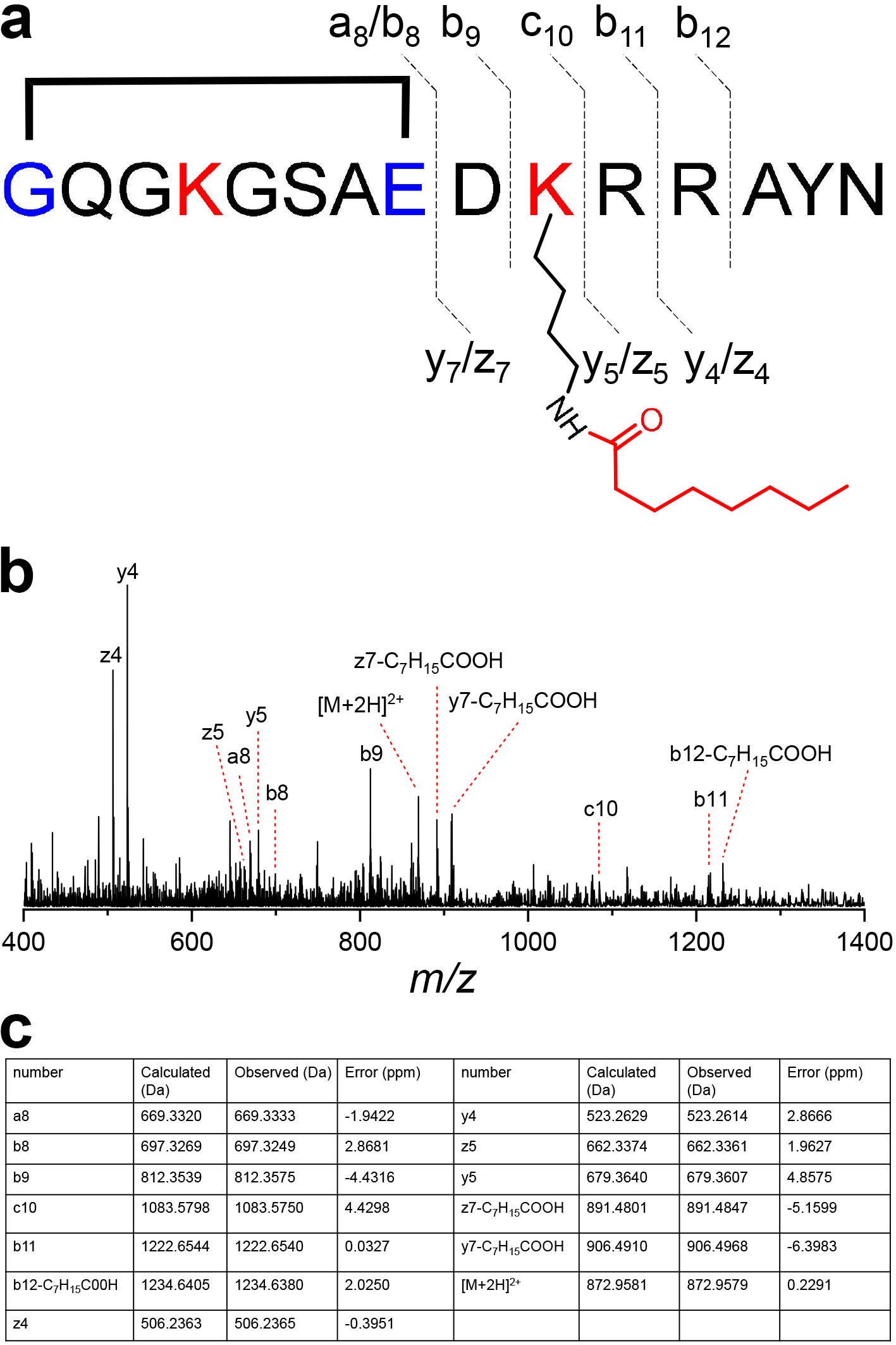

**Supplementary Fig. 24. HRMS/MS analysis of 10.** **a**, Annotation of fragment ions of **10**. **b**, HRMS/MS spectrum of **10**. **c**, Detailed information of identified fragment ions. Structure is shown in simplified model with unthreaded conformation for clarity.

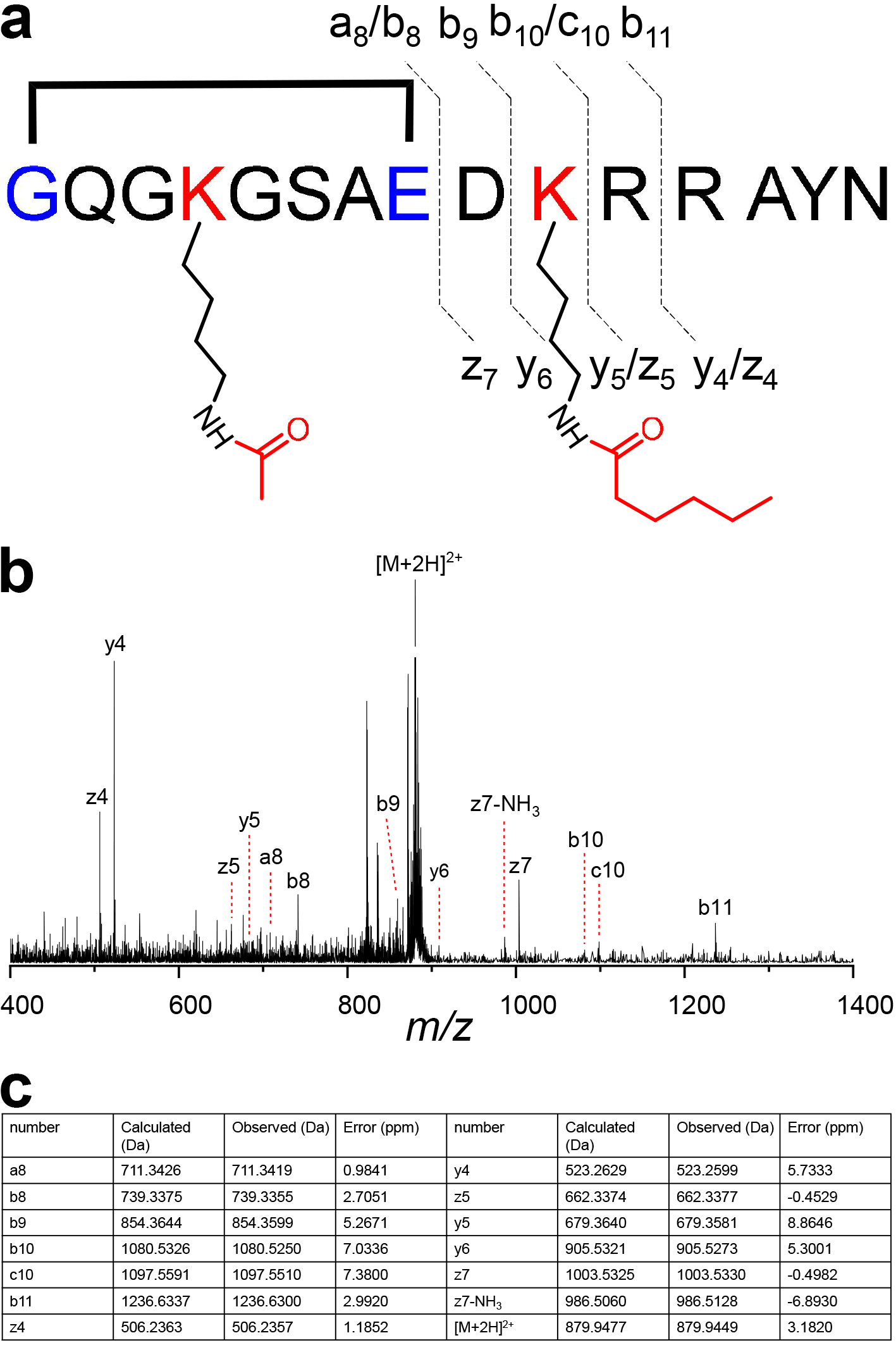

**Supplementary Fig. 25. HRMS/MS analysis of 17.** **a**, Annotation of fragment ions of **17**. **b**, HRMS/MS spectrum of **17**. **c**, Detailed information of identified fragment ions. Structure is shown in simplified model with unthreaded conformation for clarity.

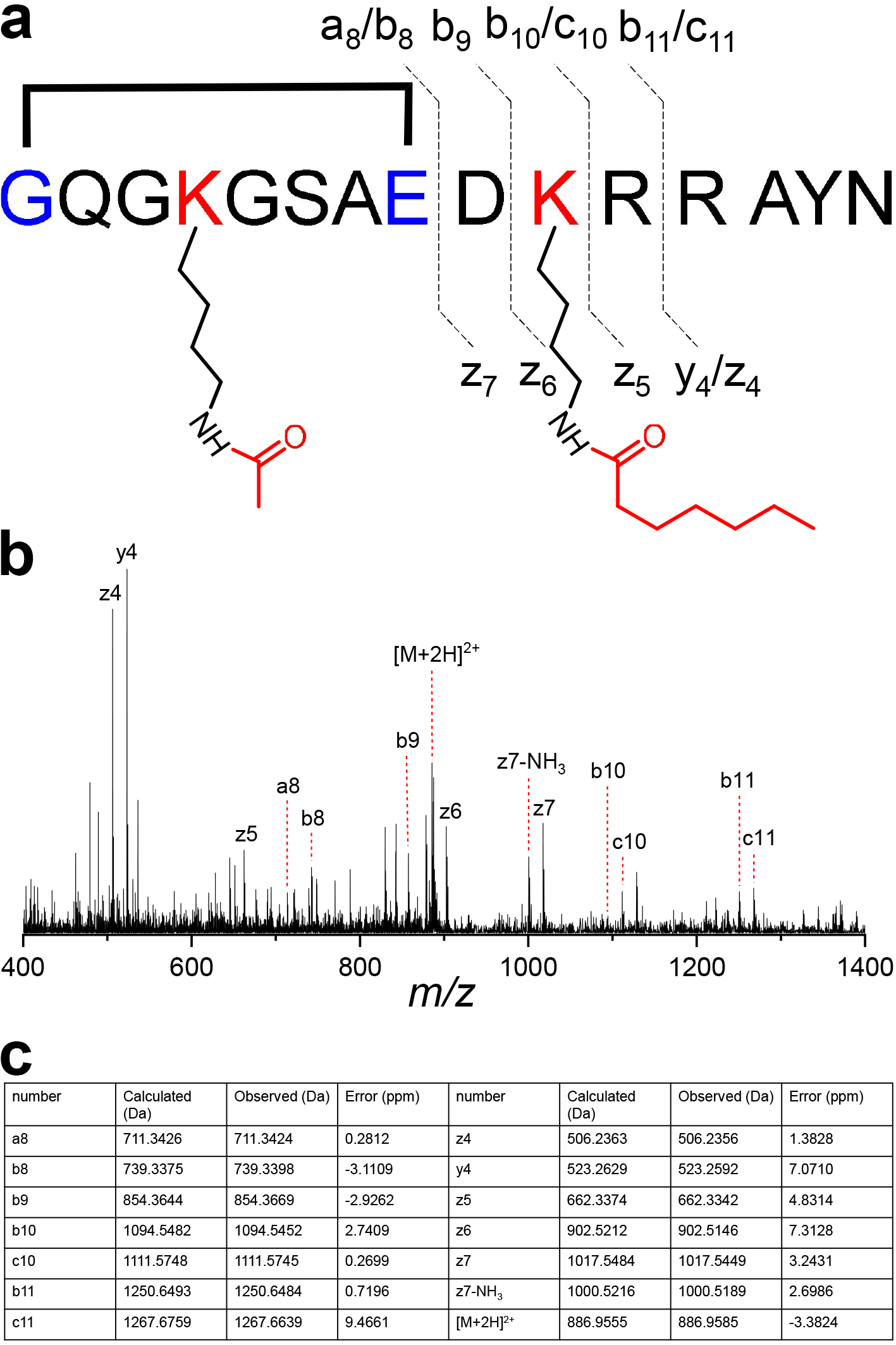

**Supplementary Fig. 26. HRMS/MS analysis of 18.** **a**, Annotation of fragment ions of **18**. **b**, HRMS/MS spectrum of **18**. **c**, Detailed information of identified fragment ions. Structure is shown in simplified model with unthreaded conformation for clarity.

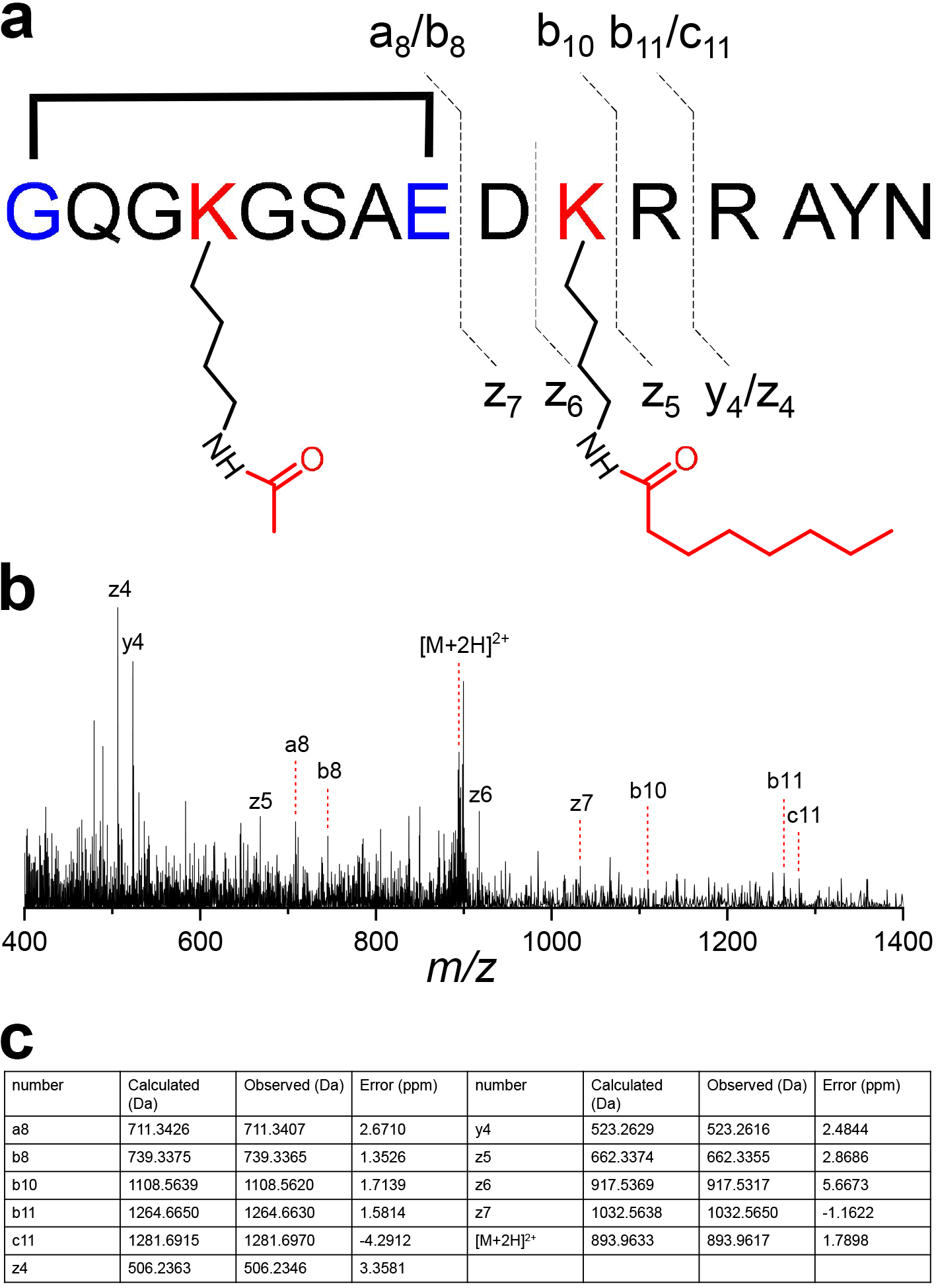

**Supplementary Fig. 27. HRMS/MS analysis of 19.** **a**, Annotation of fragment ions of **19**. **b**, HRMS/MS spectrum of **19**. **c**, Detailed information of identified fragment ions. Structure is shown in simplified model with unthreaded conformation for clarity.
